# A specific, glycomimetic Langerin ligand for human Langerhans cell targeting

**DOI:** 10.1101/286021

**Authors:** Eike-Christian Wamhoff, Jessica Schulze, Lydia Bellmann, Gunnar Bachem, Felix F. Fuchsberger, Juliane Rademacher, Martin Hermann, Barbara Del Frari, Rob van Dalen, David Hartmann, Nina M. van Sorge, Oliver Seitz, Patrizia Stoitzner, Christoph Rademacher

**Author notes:** These authors contributed equally (Jessica Schulze, Lydia Bellmann).

## Abstract

Langerhans cells are a subset of dendritic cells residing in the epidermis of the human skin. As such, they are key mediators of immune regulation and have emerged as prime targets for novel transcutaneous cancer vaccines. Importantly, the induction of protective T cell immunity by these vaccines requires the efficient and specific delivery of both tumor-associated antigens and adjuvants. Langerhans cells uniquely express Langerin (CD207), an endocytic C-type lectin receptor. Here, we report the discovery of a specific, glycomimetic Langerin ligand employing a heparin-inspired design strategy that integrated NMR spectroscopy and molecular docking. The conjugation of these glycomimetics to liposomes enabled the specific and efficient targeting of Langerhans cells in the human skin. This delivery platform provides superior versatility and scalability over antibody-based approaches and thus addresses current limitations of dendritic cell-based immunotherapies.

## Introduction

The human skin is an attractive vaccination site due to the high density of immune cells compared to other organs such as the muscle^1^. The highly efficacious and cost-effective small pox vaccine was first used via this administration route and has thus proven its feasibility^2^. The skin contains several subsets of dendritic cells (DCs), immune cells that are specialized in the internalization of pathogens and the presentation of antigens to induce T cell responses^3^. Langerhans cells (LCs) constitute a subset of DCs residing in the epidermis of the stratified as well as the mucosal skin. Following their activation, LCs migrate to the draining lymph nodes to elicit systemic immune responses^4^. Due to their localization in the epidermis and their ability to cross-present exogenous antigens to cytotoxic T cells, LCs have emerged as promising targets for transcutaneous vaccination strategies^5–7^. Various approaches avoiding hypodermic injections have been explored to improve patient care and vaccination costs^1^. For example, microneedles and thermal ablation overcome the stratum corneum and thereby facilitate antigen delivery to the skin.

Sipuleucel-T, an adoptive cell therapy for prostate cancer, has provided proof of concept for the induction of protective cytotoxic T cell responses against tumor-associated antigens (TAAs) by myeloid immune cells^8^. Moreover, the adoptive transfer of monocyte-derived DCs into melanoma patients has been demonstrated to elicit TAA-specific T cell immunity^9^. As *ex vivo* strategies remain laborious and expensive, the focus has shifted towards the delivery of antigens *in situ*^10^. Intriguingly, DCs express several endocytic receptors including chemokine receptors, scavenger receptors and C-type lectin receptors (CLRs) that promote the internalization and cross-presentation of antigens^11–13^. Pioneered by Steinman et al., the use of antibody-antigen conjugates targeting CLRs such as DEC-205, DC-SIGN and DNGR-1 represents an established strategy to deliver antigens to DCs and has translated into clinical trials^14–17^. These investigations helped identify several parameters that shape cytotoxic T cell immunity and guide the development of next-generation cancer vaccines. First, the activation of DCs by co-administration of adjuvants such as Toll-like receptor (TLR) or Rig-I-like receptor agonists is required to avoid tolerance induction^18^. Furthermore, the choice of delivery platform and targeting ligand influence the efficiency of antigen internalization, processing and cross-presentation by DCs^19–22^. Finally, the specific targeting of individual DC subsets is essential as of off-target delivery of antigens and adjuvants may result in adverse effects or compromised cytotoxic T cell immunity^23,24^. Consequently, DC subset-specific receptors such as the CLRs Langerin and DNGR-1 as well as the chemokine receptor XCR1 have become a focal point for the development of novel immunotherapies^13,17^.

Langerin (CD207), an endocytic CLR exclusively expressed on human LCs, has been shown to promote the cross-presentation of antigens to prime cytotoxic T cells^4,25^. The trimeric CLR thus represents an attractive target receptor for transcutaneous vaccination strategies^26^. In this study, we pursued the development of targeted nanoparticles as an antigen delivery platform for LCs to address the limitations of antibody-based approaches. Liposomes represent versatile nanoparticles that have been approved for the delivery of chemotherapeutics in Kaposi’s sarcoma and allow for the co-formulation of adjuvants^27,28^. They can be targeted to glycan-binding proteins (GBPs) including CLRs or sialic acid-binding immunoglobulin-like lectins (Siglecs) expressed on immune cells using glycans or glycomimetic ligands^29–31^.

Glycan recognition by Langerin is Ca^2+^ - as well as pH-dependent and consequently abrogated in the early endosome, thereby limiting lysosomal antigen degradation^32^. This release mechanism simultaneously increases the internalization capacity of LCs as unbound Langerin has been shown to recycle to the plasma membrane^33^. Hence, the use of glycans or glycomimetics provides advantages over antibody-based approaches which potentially suffer from inefficient ligand release^20,21^. Furthermore, antibodies are prone to denaturation and their large-scale production is expensive. Despite recent advances in antibody engineering such as humanization and defined glycosylation profiles, antibody-based therapies remain limited by Fc receptor-mediated immunogenicity and clearance^34^.

As glycans are typically recognized by several CLRs or other GBPs, they do not provide the specificity required to target individual DC subsets^35^. Additionally, glycan-Langerin interactions display low affinities insufficient to promote the endocytosis of liposomes^36–39^. This renders the design of potent and specific glycomimetic ligands essential for the development of an antigen delivery platform for LCs. The carbohydrate binding sites of CLRs are hydrophilic and solvent-exposed which has impeded the discovery of drug-like molecule ligands^40,41^. While mono- and oligosaccharides represent attractive scaffolds, the synthesis of carbohydrates and structural glycomimetics is generally considered onerous^42,43^. Nevertheless, individual reports have demonstrated the feasibility of ligand design for these challenging target receptors and other GBPs^44–47^. Many of these reports highlight the utility of concepts from rational and fragment-based drug discovery for glycomimetic ligand design.

Here, we present the discovery of the first micromolar glycomimetic ligand for Langerin. Combining NMR spectroscopy and molecular docking, we rationally designed heparin-derived monosaccharide analogs. The targeting ligand facilitated the endocytosis of liposomes by LCs and provides remarkable specificity over other GBPs in a physiologically relevant *ex vivo* skin model. Hence, our findings demonstrate the CLR-mediated targeting of nanoparticles to individual immune cell subsets using glycomimetics.

## Results

### Heparin-derived monosaccharides represent favorable scaffolds for glycomimetic ligand design

Aside from its function as a pathogen recognition receptor, Langerin interacts with self-antigens such as glycosaminoglycans including heparin^39,48,49^. These linear polysaccharides are composed of disaccharide repeating units consisting of galactose or uronic acids and differentially sulfated *N*-acetyl glucosamine (GlcNAc). Prompted by the 10-fold affinity increase (K_D_ = 0.49±0.05 mM) over mannose (Man) disaccharides (K_D_ = ca. 4 mM) recently reported for a heparin-derived trisaccharide, we employed ligand-observed ^19^F R_2_-filtered NMR experiments to determine K_I_ values for a set of differentially sulfated GlcNAc derivatives (Figure 1a)^37,39,50^. Interestingly, the affinities of glucosamine-2-sulfate (GlcNS) (K_I_ = 1.4±0.2 mM), *N*-acetyl glucosamine-6-sulfate (GlcNAc-6-OS) (K_I_ = 0.6±0.1 mM) and glucosamine-2-sulfate-6-sulfate (GlcNS-6-OS) (K_I_ = 0.28±0.06 mM) were comparable or higher than those observed for heparin-derived oligosaccharides and other monosaccharides including Glc (K_I_ = 21±4 mM), GlcNAc (K_I_ = 4.1±0.7 mM) and Man (K_I_ = 4.5±0.5 mM) (Supplementary Figure 1, Supplementary Table 1)^48^.

**Table 1.** Structure-activity relationship and specificity against DC-SIGN.

| Structure | Langerin |  |  | DC-SIGN |  |
| --- | --- | --- | --- | --- | --- |
|  | K<sub>I</sub> [mM] | K<sub>D</sub> [mM] | Relative potency<sup>a</sup> | K<sub>I</sub> [mM] | Specificity |
| **2** | 0.32±0.05 | 0.46±0.04<br>0.5±0.2<sup>b</sup> | 31 | 17±1 | 53 |
| **16** | 0.24±0.03 | 0.23±0.07<br>0.3±0.1<sup>b</sup> | 42 | 15±3 | 63 |
| Man | 4.5±0.5<sup>c</sup> | 6.5±0.2<sup>c</sup> | 2.2 | 3.0±0.3 | 0.67 |
| **21** | 10±1 | 12±1 | 1.0 | 2.7±0.3 | 0.27 |

<sup>a</sup> The relative potency was calculated utilizing the K<sub>I</sub> value determined for **21**.
<sup>b</sup> The value was determined from integrals V of resonances observed to be in slow exchange.
<sup>c</sup> The value was previously published<sup>50</sup>.

**Figure 1.**
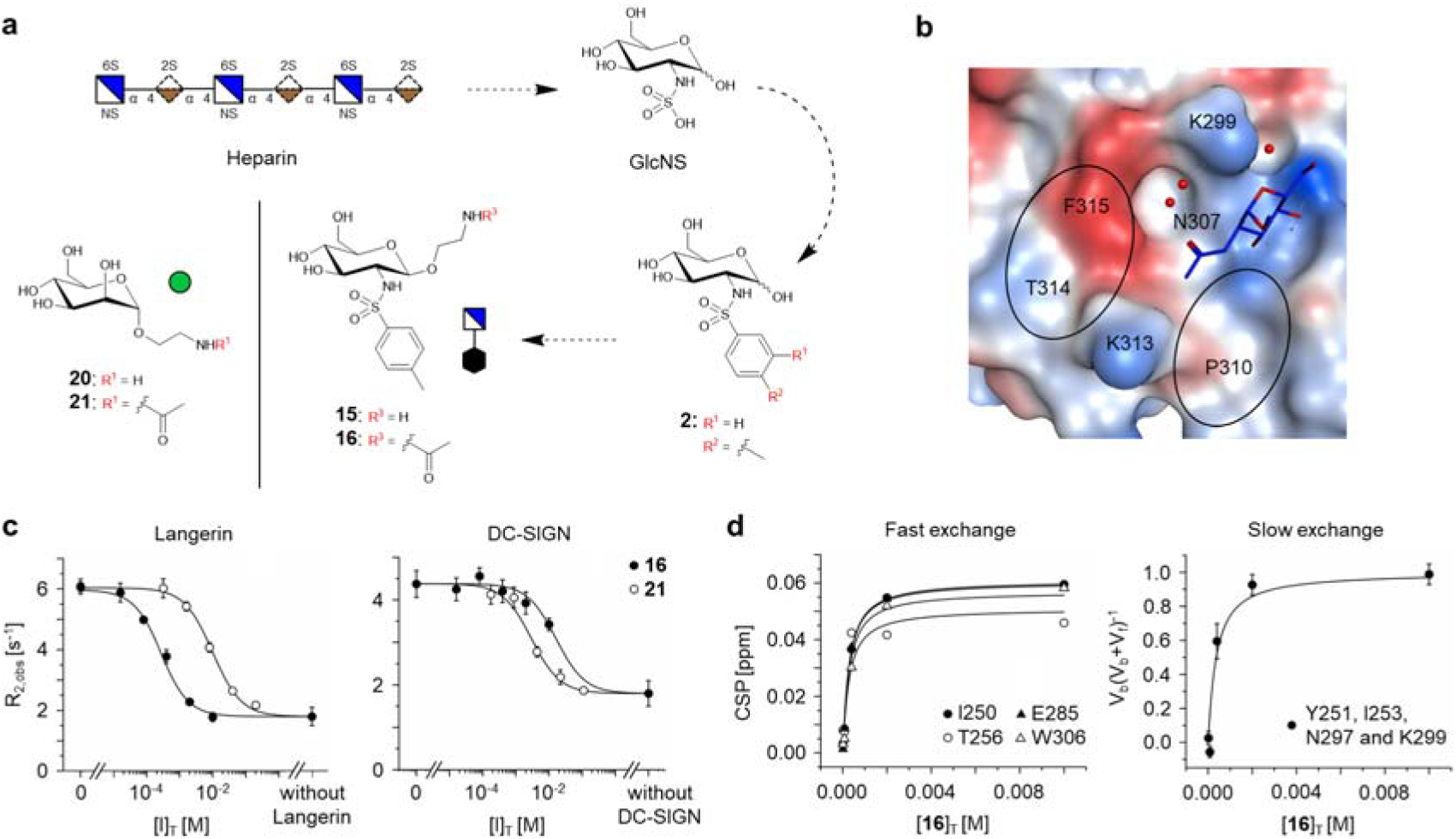
Heparin-inspired design of glycomimetic targeting ligands for Langerin. **a.** The heparin-derived monosaccharide GlcNS was identified as a favorable scaffold for glycomimetic ligand design. The design of GlcNS analogs lead to the discovery of glycomimetic targeting ligand **15**. **15** bears an ethylamino linker in β-orientation of C1 for conjugation to the delivery platform. **20** served as a Man-based reference molecule throughout this study. **b.** Based on the binding mode of GlcNAc (PDB code: 4N32), the small aromatic substituents in C2 were hypothesized to increase the affinity by the formation of cation-π interactions with K299 and K313 or π-π and H-π interactions with F315 and P310^38^. The receptor surface is colored according to its lipophilicity (lipophilic: red, hydrophilic: blue). **c.** ^19^F R_2_-filtered NMR experiments revealed a 42-fold affinity increase for model ligand **16** (K_I_ = 0.24±0.03 mM) over Man-based reference molecule **21** (K_I_ = 10±1 mM). Additionally, **16** displayed an encouraging specificity against DC-SIGN (K_I,DC-SIGN_ = 15±3 mM). **d.** The affinity of **16** for Langerin was validated in ^15^N HSQC NMR experiments analyzing resonances in the fast (K_D,fast_ = 0.23±0.07 mM) and the slow (K_D,slow_ = 0.3±0.1 mM) exchange regime.

GlcNS-6-OS, representing the most potent monosaccharide identified, displayed an additive structure-activity relationship (SAR) for the sulfation in C2 and C6. This affinity increase is based on the formation of salt bridges with K299 and K313 as previously shown by X-ray crystallography^51^. GlcNS-6-OS displayed an altered orientation of the Glc scaffold compared to the Langerin-GlcNAc complex (Figure 1b)^38^. The affinity increase observed over Glc observed for GlcNAc, on the other hand, is the result of an H_2_O-mediated hydrogen bond with K299. Importantly, either of these interactions might be leveraged for glycomimetic ligand design via the bioisosteric substitution of the sulfate groups with a sulfonamide linker. In particular, the synthesis of GlcNS analogs represents a feasible fragment growing approach to explore the carbohydrate binding site for favorable interactions (Figure 1a). These characteristics render sulfated GlcNAc derivatives favorable scaffolds for the design of glycomimetic Langerin ligands.

### Small aromatic sulfonamide substituents render glycomimetics potent targeting ligands for Langerin and provide specificity against DC-SIGN

Assuming the conservation of the Glc scaffold orientation observed for GlcNAc, small aromatic substituents in C2 were hypothesized to increase the affinity by the formation of cation-π interactions with K299 and K313 or π-π and H-π interactions with F315 and P310, respectively (Figure 1b). Accordingly, a panel of GlcNS analogs **1** to **5** bearing differentially substituted phenyl rings was prepared and followed by the determination of K_I_ values (Figure 1a, Supplementary Scheme 1, Supplementary Figure 2). Increased affinities over GlcNAc were observed for all analogs, with a 13-fold affinity increase for **2** (K_I_ = 0.32±0.05 mM), the most potent panel member (Table 1, Supplementary Figure 3, Supplementary Table 2). The analog bears a methyl group in para position of the phenyl ring that does not contribute substantially to the affinity increase, as exemplified by the K_I_ value obtained for **1** (K_I_ = 0.37±0.04 mM).

**Figure 2.**
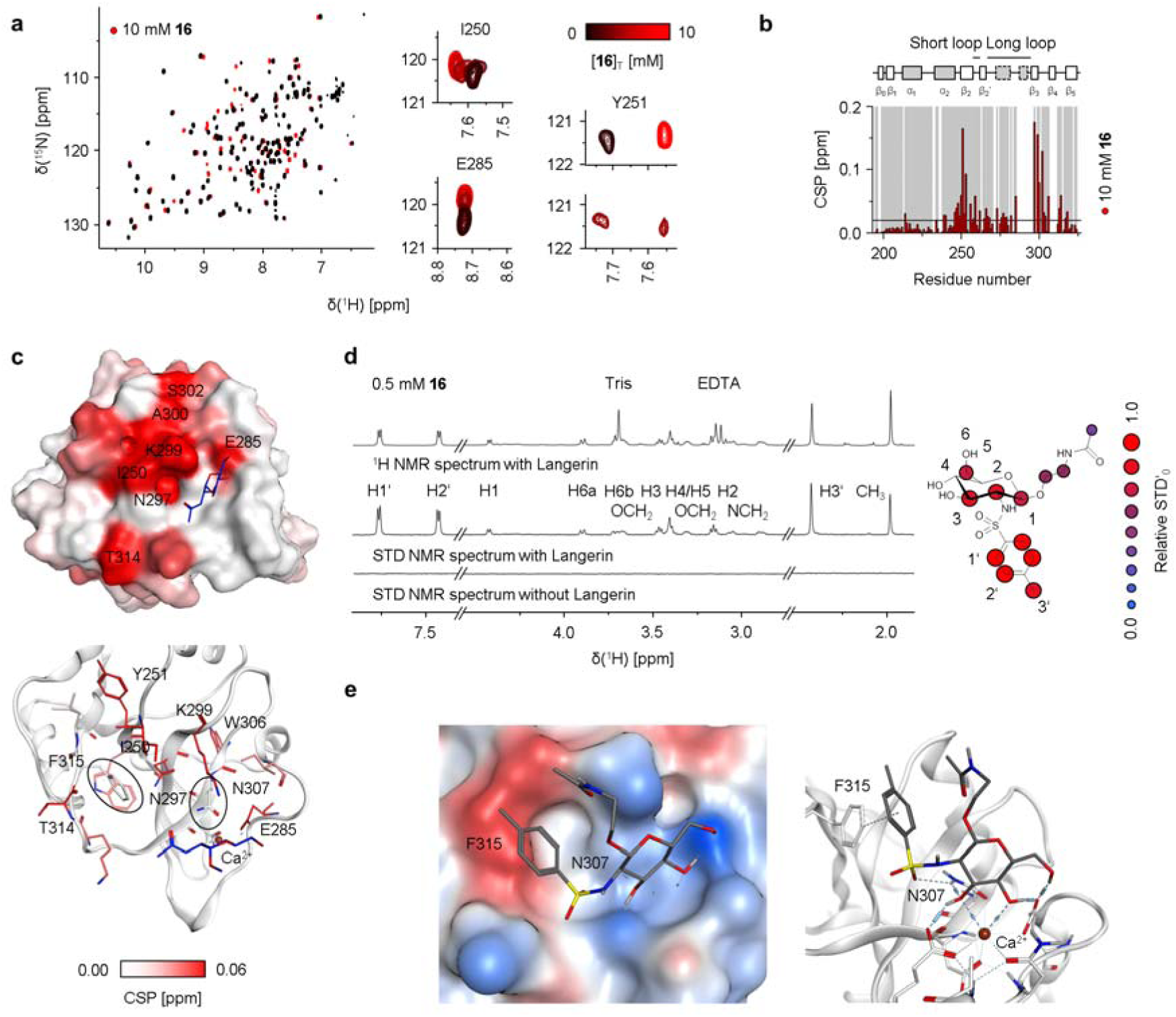
Binding mode analysis for the glycomimetic targeting ligand. **a. and b.** ^15^N HSQC NMR experiments revealed the CSP pattern for **16**. Upon titration, fast exchanging resonances such as I250 and E285 as well as slow exchanging resonances including Y251 were observed. **c.** Mapping the CSPs on the X-ray structure of Langerin in complex with GlcNAc (PDB code: 4N32) validated a Ca^2+^-dependent binding mode as indicated by CSPs observed for E285 and K299^38^. Compared to titrations with **21,** Y251, I250 and T314 displayed a relative CSP increase, while a decrease was observed for K313 (Supplementary Figure 6). Overall, the majority of residues displaying increased CSPs can be associated with N307 and F315, which could not be assigned^32^ **d.** STD NMR experiments served to further validate the interaction formed between **16** and Langerin. STD NMR spectra were recorded at saturation times t_sat_ of 0.4 s and are magnified 8-fold. Epitopes determined from build-up curves suggest strong interactions formed by the phenyl substituent (Supplementary Figure 10). By contrast, low relative STD’_0_ values were observed for the acetylated ethylamino linker, consistent with a solvent exposed orientation. **e. 16** was docked into the carbohydrate binding site to rationalize the observations from ^15^N HSQC and STD NMR experiments. The selected docking pose predicted the formation of π-π interactions between the phenyl ring and F315 as well as the formation of a hydrogen bond between the sulfonamide group and N307. The linker displays high solvent exposure. The receptor surface is colored according to its lipophilicity (lipophilic: red, hydrophilic: blue).

**Figure 3.**
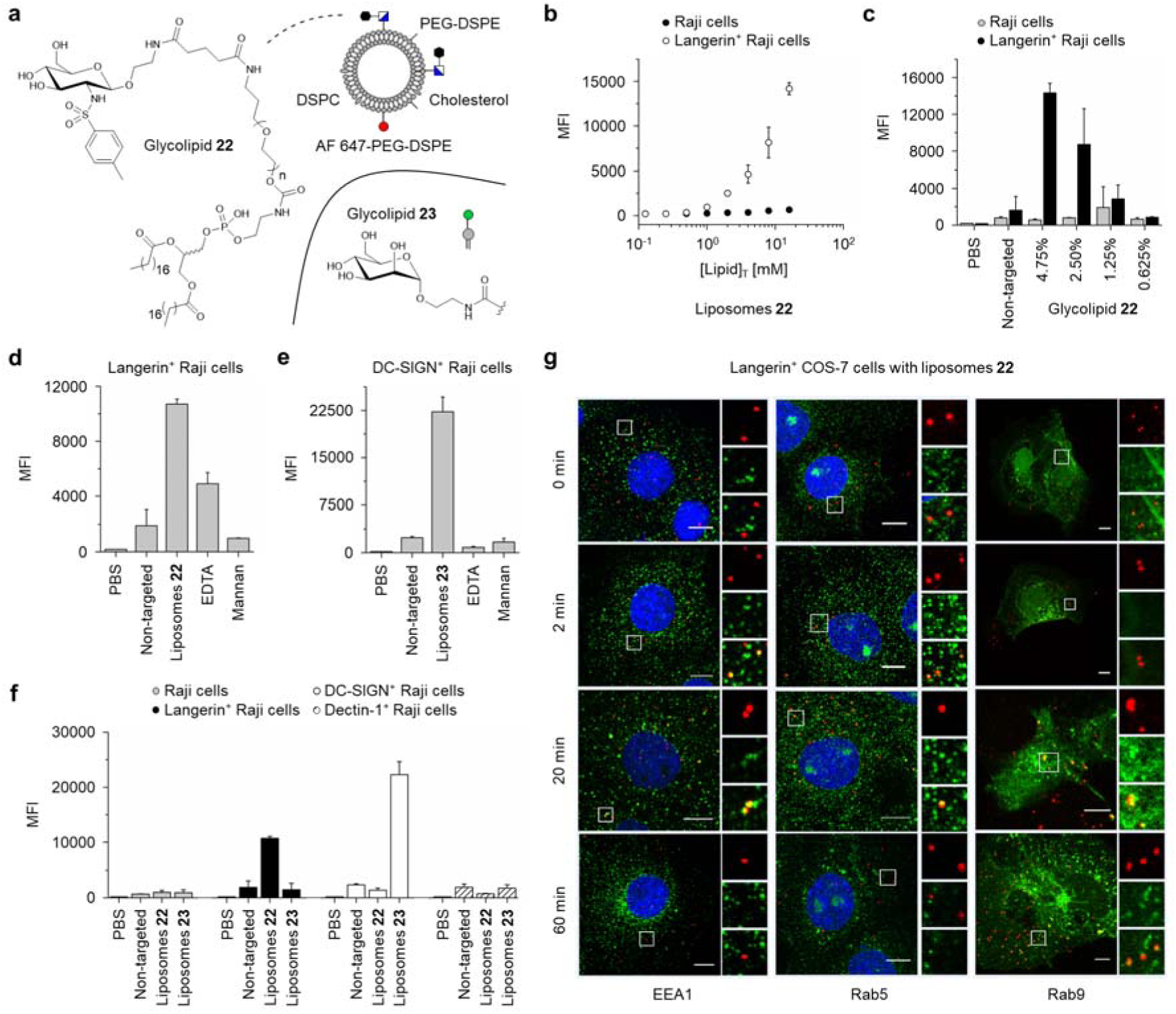
*In vitro* targeting of Langerin^+^ human model cells. **a.** Targeted liposomes were prepared via thin film hydration and pore extrusion by incorporation glycolipids **22** or **23**. The binding of the liposomes at 4°C to human Raji cells expressing different CLRs was investigated by flow cytometry. **b.** Dose-dependent binding of liposomes **22** was observed for Langerin^+^ cells. The number of liposomes used is expressed as the total concentration of lipids [Lipid]_T_. All subsequent experiments were conducted at a concentration [Lipid]_T_ of 16 µM. **c.** Binding of liposomes **22** to Langerin^+^ cells was furthermore dependent on the amount of the incorporated glycolipid. Only negligible unspecific binding of non-targeted liposomes to Raji cells was observed. All subsequent experiments were conducted at 4.75% of glycolipids **22** or **23**. **d.** Competition experiments with EDTA and mannan validated Ca^2+^-and carbohydrate binding site-dependent binding to Langerin^+^ cells for liposomes **22**.**e.** Analogously, liposomes **23**, bearing the Man on their surface, were observed to bind DC-SIGN^+^ Raji cells via the Ca^2+^-dependent carbohydrate binding site. **f.** Among the set of analyzed CLRs including Langerin, DC-SIGN and Dectin-1, liposomes **22** were found to be specific for Langerin^+^ cells, while liposomes **23** exclusively bound to DC-SIGN^+^ cells. MFI values were determined from three independent experiments. **g.** The intracellular trafficking of liposomes **22** (red) in Langerin^+^ COS-7 cells was analyzed by confocal microscopy. Co-localization with the early endosomal compartment was observable 2 min after incubation at 37°C using either the EEA-1 or the Rab5 marker (green). Liposomes remained associated with this compartment for at least 20 min. At this time point, a subset of liposomes was trafficked into the late endosomal compartment as indicated by the co-localization with the Rab9 marker (green). The scale bars indicate 10 µm.

Despite its low complexity, **2** displays an affinity superior to that of glycans previously applied as targeting ligands for DC subsets distinct from LCs^29^. Here, the blood group antigen Le^X^ (K_D,DC-SIGN_ = ca. 1 mM) was demonstrated to promote the DC-SIGN-dependent internalization of liposomes by isolated dermal DCs to activate T cells *in vitro*^52^. Encouraged by these reports, we advanced **2** towards targeted delivery applications via the introduction of an ethylamino linker in β-orientation of C1 of the Glc scaffold to yield targeting ligand **15** (Figure 1a, Supplementary Scheme 2, Supplementary Figure 2).

After acetylation of the amino group, we obtained model ligand **16** (Figure 1a, Supplementary Scheme 2, Supplementary Figure 2). The K_I_ value determination for **16** (K_I_ = 0.24±0.03 mM) revealed a 42-fold affinity increase over the Man-based reference molecule **21** (K_I_ = 10±1 mM) (Figure 1a and c, Table 1, Supplementary Scheme 2 and 3, Supplementary Figure 2 and 4). To validate these affinities and to expand our insight into the recognition process, orthogonal protein-observed ^15^N HSQC NMR experiments were performed (Figure 1d and 2a, Table 1, Supplementary Figure 5). Notably, a considerable fraction of the resonances displaying chemical shift perturbations (CSPs) upon the addition of **16** also displayed line broadening Δν_0.5_ of more than 10 Hz, indicative of intermediate exchange phenomena. Accordingly, these resonances were not considered for K_D_ determination. Simultaneously, slow exchange phenomena were observed for a set of resonances corresponding to Y251, I253, N297 and K299 (Figure 2a). Analysis of both fast- and slow-exchanging peaks revealed affinities comparable to the K_I_ values obtained for **16** (K_D,fast_ = 0.23±0.07 mM, K_D,slow_ = 0.3±0.1 mM) as well as **21** (K_D_ = 12±1 mM) (Figure 1d, Table 1, Supplementary Figure 5). Likewise, the affinity of **2** was validated using ^15^N HSQC NMR (K_D,fast_ = 0.46±0.04 mM, K_D,slow_ = 0.5±0.2 mM) (Table 1, Supplementary Figure 6).

**Figure 4.**
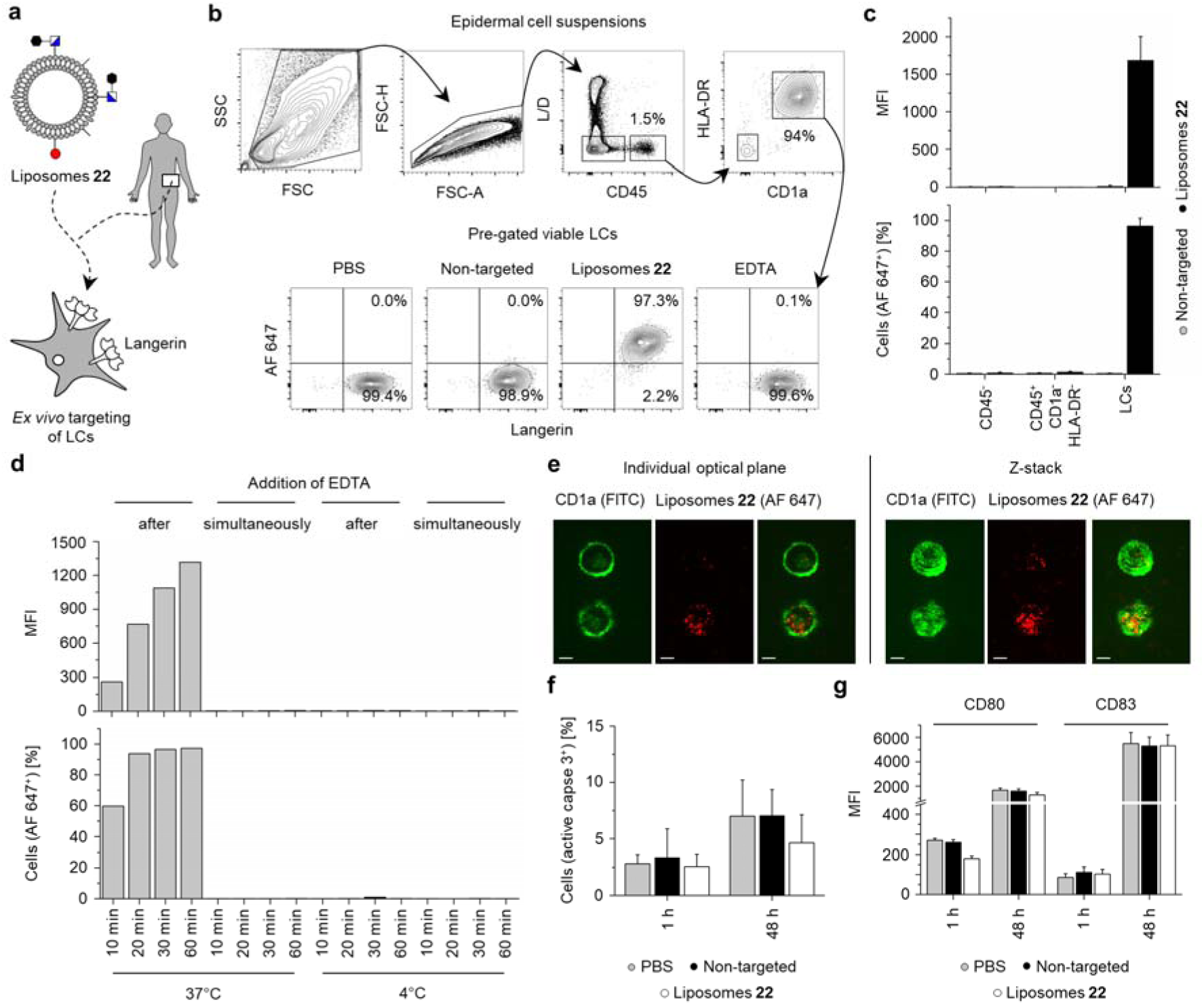
*Ex vivo* targeting of human LCs in epidermal cell suspensions. **a.** LC targeting by liposomes **22** was investigated *ex vivo* using flow cytometry. To this end, epidermal cell suspensions were prepared as previously described and incubated at 37°C^53^. **b. and c.** LCs were identified as viable HLA-DR^+^-CD45^+^-CD1a^high^ cells. The binding and endocytosis of liposomes **22** by human LCs was detected via the fluorescence signal of AF 647. Selectivity for LCs over keratinocytes and T cells was reproducibly demonstrated in seven independent experiments and quantified via the fraction of AF 647^+^ cells. Additionally, the corresponding MFI for representative subset of experiments is depicted. **d.** The kinetics of endocytosis by LCs were analyzed at different temperatures in three independent experiments. Simultaneous incubation with liposomes **22** and EDTA resulted in complete inhibition of endocytosis. By contrast, the addition of EDTA 20 min after incubation at 37°C did not alter the fraction of AF 647^+^ LCs, indicating efficient endocytosis. As expected, endocytosis was abrogated at 4°C. One representative experiment is depicted. **e.** LCs in epidermal cell suspensions were identified by addition of a fluorescently labeled anti-CD1a antibody and internalization of liposomes **22** was visualized by confocal microscopy at 37°C. The scale bars indicate 4 µm. **f.** The cytotoxicity of liposomes **22** for LCs was monitored in four independent experiments. No significant increase in active caspase 3 levels due to incubation with liposomes was observed after 1 h or 48 h. **g.** Furthermore, the incubation with liposomes **22** for 2 h and 48 h in four independent experiments did not significantly increase the expression levels of CD80 or CD83, indicating the absence liposome-mediated LC activation *ex vivo*.

**Figure 5.**
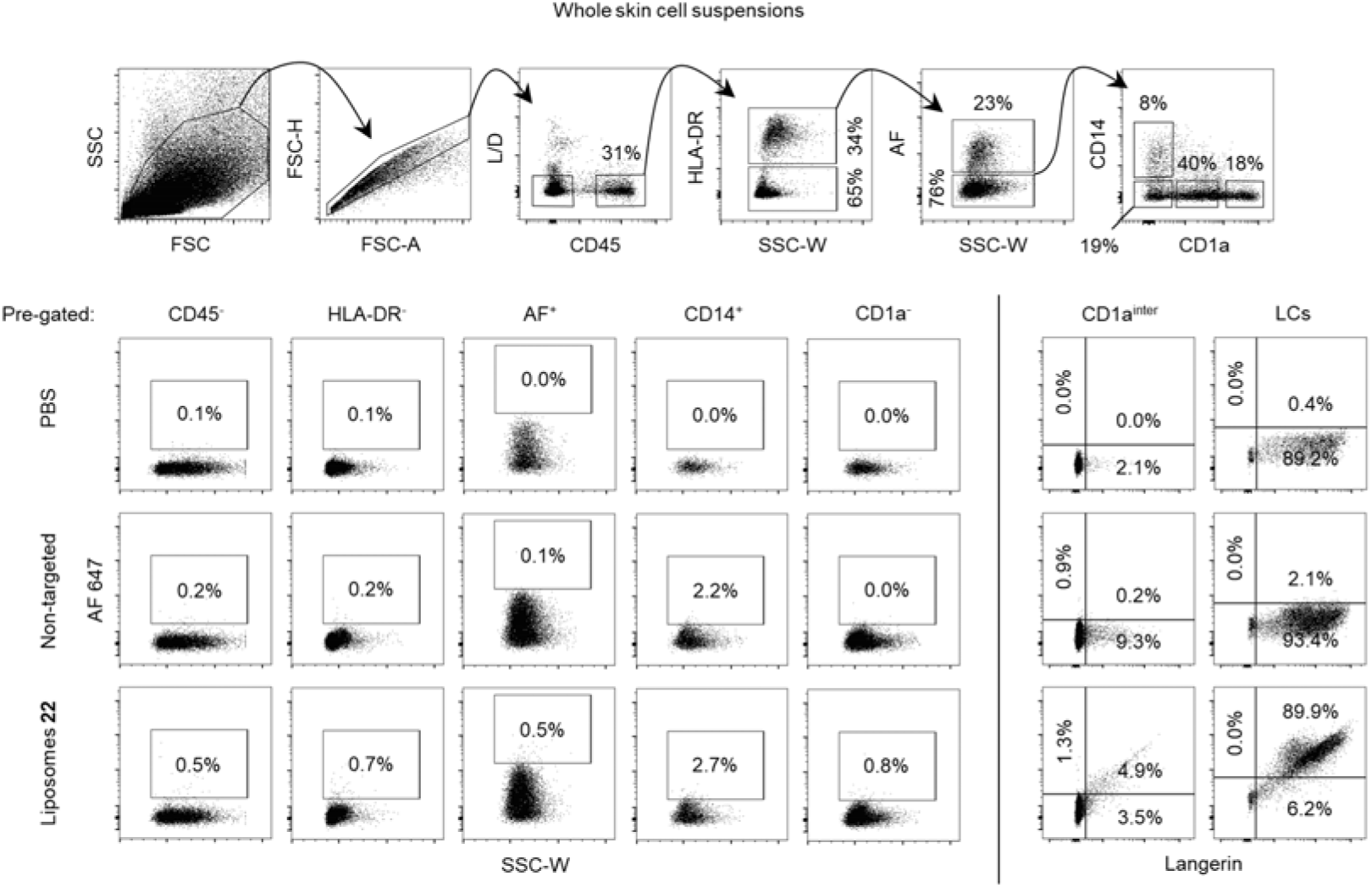
*Ex vivo* targeting of human LCs in whole skin cell suspensions. The specificity of liposomes **22** for LCs in the context of the human skin was evaluated at 37°C by flow cytometry. To this end, whole skin suspensions were prepared as previously published^96^. LCs were identified as viable HLA-DR^+^-CD45^+^-CD1a^high^ cells while dermal DCs were identified as viable HLA-DR^+^-CD45^+^-CD1a^intermediate^ cells. Monocytes and macrophages were characterized by the expression of CD14. Strikingly, binding and endocytosis of liposomes **22** was exclusively observed for LCs. The depicted plots are representative of three independent experiments.

Next, we explored the specificity of targeting ligand **16** against DC-SIGN as such off-target affinity would imply a reduced efficiency of the approach and the potential induction of adverse effects. For this purpose, we transferred the ^19^F R_2_-filtered NMR reporter displacement assay to DC-SIGN (Supplementary Figure 7, Supplementary Table 3). Strikingly, **16** (K_I,DC-SIGN_ = 15±3 mM) displayed a considerably decreased K_I_ for DC-SIGN compared to Langerin corresponding to 63-fold specificity (Figure 1c, Table 1). At the same time **21** displayed 3.7-fold specificity for DC-SIGN over Langerin (K_I,DC-SIGN_ = 2.7±0.3 mM). A comparison with the affinities determined for **2** (K_I,DC-SIGN_ = 17±1 mM) and Man (K_I,DC-SIGN_ = 3.0±0.3 mM) revealed that the differential recognition of α- and β-glycosides by these CLRs contributes to specificity (Table 1, Supplementary Figure 8).

### The formation of **π**-**π** interactions and hydrogen bonds by aromatic sulfonamide substituents mediate an affinity increase for Langerin

To investigate the binding mode of model ligand **16**, ^15^N HSQC and STD NMR experiments were combined with molecular docking studies (Figure 2a to e). Here, the orientation of the linker was of particular interest to evaluate the compatibility of the binding mode with the presentation of targeting ligand **15** on liposomes.

Titration of **16** induced CSPs for E285 and K299 providing further evidence for a canonical Ca^2+^-dependent binding mode of the Glc scaffold of the glycomimetic (Figure 2b and c). These protein-observed NMR experiments additionally revealed strong CSPs for residues in proximity of F315 and N307. Notably, both residues could not be assigned, likely due to their association with the flexible long loop^32^. This effect is accompanied by a decreased the CSP for K313 compared to titrations with Man analog **21** (Supplementary Figure 5 and 9). Both observations are conserved in titrations with **2** and indicate an orientation of the phenyl ring towards F315 or K299 rather than K313 or P310 (Supplementary Figure 6 and 9). Interestingly, additional CSPs were induced for residues remote from the carbohydrate binding, suggesting the modulation of an allosteric network involved in the regulation of Ca^2+^ recognition by Langerin (Supplementary Note 1)^32^.

To complement the protein-observed NMR experiments and to investigate the orientation of the acetylated ethylamino linker, STD NMR epitope mapping with **16** and **21** was conducted. The binding epitope of **16** was dominated by uniformly high STD effects for the phenyl ring and thus supports a model in which favorable secondary interactions are formed between this substituent and the Langerin surface (Figure 2d, Supplementary Figure 10). The acetylated ethylamino linker did, by contrast, display uniformly low STD effects indicating a solvent exposed orientation and validating the developed conjugation strategy for GlcNS analogs. Similarly, the ethylamino linker of **21** received decreased STD effects compared to the Man scaffold (Supplementary Figure 11 and 12).

Finally, molecular docking was performed utilizing the X-ray structure of the Langerin complex with GlcNAc (Figure 2e, Supplementary Figure 13)^38^. Generated docking poses were evaluated in the context of the NMR experiments and representative poses were selected to visualize the formation of potential secondary interactions. Indeed, orienting the phenyl ring towards F315 resulted in the formation of π-π interactions. This orientation also coincided with the formation of a weak hydrogen bond between the sulfonamide linker and N307. Both interactions explain the pronounced CSP values observed for residues that are associated with F315 and N307 including I250, Y251, N297 and K299. Furthermore, the phenyl ring received high STD effects indicating the formation of secondary interaction and high proximity to the Langerin surface. Conversely, the acetylated ethylamino linker displayed high solvent exposure and no conserved secondary interactions for the majority of docking poses. This observation was in accordance with the low STD effects and thus validated the developed conjugation strategy for GlcNS analogs. Overall, we propose a binding mode for **16** that displays a conserved orientation of the Glc scaffold, consistent with both STD and ^15^N HSQC NMR experiments. The affinity increase can be rationalized by the formation of π-π interactions between the phenyl substituent and F315 as well as a hydrogen bond between the sulfonamide linker and N297.

### Targeted liposomes specifically bind to Langerin^+^ cells *in vitro*

Next, monosaccharide analogs **15** or **20** were utilized to synthesize glycolipids **22** and **23**, respectively (Figure 3a, Supplementary Scheme 4). Their affinity for Langerin was evaluated in a plate-based enzyme-linked lectin assay (ELLA) (Supplementary Figure 14)^29^. While a dose-dependent interaction could be demonstrated for **22**, no interaction was detected for the immobilization of **23**. This validates the determined affinity increase of model ligand **16** over the Man-based reference molecule **21**. Encouraged by these findings, we prepared targeted liposomes labeled with Alexa Fluor (AF) 647 with a diameter d of 160±60 nm that were stable over several months when stored at 4°C in PBS (Figure 3a, Supplementary Figure 14). ^1^H NMR experiments were employed to probe the accessibility of targeting ligand **15** on the surface of the liposomes. Interestingly, two states were observed for the resonances corresponding to H1’ and H2’ of the phenyl ring (Supplementary Figure 14). Both states displayed linewidths ν_0.5_ smaller than 30 Hz, suggesting residual flexibility due to the presentation of the targeting ligand on an extended polyethylene glycol linker. The alternative state potentially corresponds to targeting ligands oriented towards the lumen of the liposomes. In summary, **15** is likely presented favorably on the surface of the liposomes to enable interactions with Langerin, further validating the developed conjugation strategy.

The binding of the targeted liposomes to Langerin^+^ Raji model cells was evaluated via flow cytometry (Supplementary Figure 15). Indeed, initial titration experiments revealed dose- and Langerin-dependent binding of liposomes **22**, as well as negligible cytotoxicity (Figure 3b, Supplementary Figure 15 and 16). The avidity of the interaction was furthermore dependent on the fraction of glycolipid **22** incorporated into the liposomal formulation, with negligible unspecific interactions observed for non-targeted liposomes (Figure 3c). As expected, binding of the targeted liposomes could be abrogated via the addition of EDTA or the Man-based polysaccharide mannan to inhibit Ca^2+^-dependent glycan recognition (Figure 3d). Analogously, liposomes **23**, bearing Man on their surface bound to DC-SIGN^+^ Raji cells (Figure 3e). Strikingly, binding of these liposomes was not detected for Langerin^+^ or Dectin-1^+^ cells, suggesting an avidity threshold for liposomal targeted delivery. Most importantly, Langerin-targeted liposomes specifically bound to Langerin^+^ cells, not to DC-SIGN or Dectin-1 expressing cells (Figure 3f). The intracellular trafficking of liposomes **22** was followed in Langerin^+^ COS-7 cells. Upon internalization, the liposomes co-localized with the early endosomal markers EEA1 and Rab5 within 2 minutes lasting up to at least 20 minutes (Figure 3g). At this later time point, a subset of liposomes was trafficked into the late endosomal compartment as demonstrated by co-staining with Rab9 as a marker.

### Langerhans cells of the human epidermis efficiently internalize targeted liposomes

To explore the binding and subsequent internalization of the delivery platform by primary cells, we prepared epidermal cell suspensions from skin biopsies (Figure 4a)^53^. The cells were incubated with targeted and non-targeted liposomes for 1 h at 37°C and analyzed by flow cytometry. Upon incubation with liposomes **22**, more than 95% of gated HLA-DR^+^-CD45^+^-CD1a^high^ LCs were found to display AF 647^+^ fluorescence (Figure 4b). As for the Raji cells, binding was dependent on the targeting ligand and could be abrogated by simultaneous incubation with EDTA. The interaction was highly specific in the context of the human epidermis as neither keratinocytes, melanocytes nor T cells were targeted (Figure 4c).

Next, the kinetics of endocytosis by LCs were evaluated by adding EDTA at different times after the incubation with the delivery platform (Figure 4d). From these experiments, it can be inferred that more than 95% of gated LCs had internalized targeted liposomes after 20 min. The continuous increase in AF 647^+^ fluorescence was monitored for up to 60 min, further highlighting the efficient endocytosis by LCs that was expectedly abrogated at 4°C. The internalization of liposomes 22 was additionally demonstrated via confocal microscopy (Figure 4e). Similar to the Langerin^+^ Raji cells, the liposomal formulations displayed no cytotoxicity with LCs as indicated by the analysis of active caspase3 levels (Figure 4f, Supplementary Figure 17). Finally, we evaluated whether liposomes 22 would activate LCs *ex vivo* (Figure 4g, Supplementary Figure 17). The expression level of neither CD80 nor CD83 was significantly increased after incubation with non-targeted or targeted liposomes for 1 h. LCs in epidermal cell suspension matured within 48 h. However, this process was not affected by liposomes 22.Targeted liposomes exclusively address Langerin^+^ cells of the human skin

As an alternative to epicutaneous administration, intradermal injection represents an attractive vaccination strategy for the skin^26,54^. However, the human dermis contains additional antigen-presenting cells including dermal DCs, macrophages and monocytes. These cells express a variety of GBPs such as MR, Dectin-1, DC-SIGN and Siglec-10 and hence represent potential targets for glycomimetics^55^. In analogy to the experiments with epidermal skin cell suspensions, whole skin cell suspensions were utilized to analyze the specificity of the delivery platform in a physiologically relevant context (Figure 5)^53^. Again, targeted liposomes were efficiently endocytosed by LCs. Additionally, a minor population of CD1a^intermediate^-Langerin^+^ cells also capable of internalizing liposomes were identified. These cells might constitute a dermal DC subset but are most likely migrating LCs. Remarkably, endocytosis by CD1a^intermediate^-Langerin^−^ dermal DCs and other cell populations was negligible. Approximately 3% of CD14^+^ macrophages and monocytes were targeted by liposomes **22**, comparable to the population non-specifically internalizing non-targeted liposomes. Overall, the delivery platform was found to be highly specific for LCs in the context of the human skin.

## Discussion

Human LCs have been recognized for their capacity to internalize and cross-present exogenous antigens to elicit cytotoxic T cell responses, an established strategy for the development of novel cancer immunotherapies^5,6^. They reside in the epidermis of the skin and have consequently emerged as prime targets for transcutaneous vaccines^7,26^. However, the induction of protective T cell immunity remains challenging, requiring the efficient and specific delivery of antigens as well as adjuvants. In the absence of adjuvants, LCs induce tolerance while the off-target delivery of antigens or adjuvants potentially results in adverse effects and compromises the quality of the cytotoxic T cell response^23,24^. In this study, we present the development of a liposomal delivery platform that specifically addresses LCs in the context of the human skin to overcome these challenges.

Beyond their relevance for transcutaneous vaccination strategies, our findings provide the proof-of-concept for CLR-mediated targeting of nanoparticles to individual immune cell subsets using glycomimetics. The discovery of ligand **15** (K_I_ = 0.24±0.03 mM) with micromolar affinity for Langerin represents the essential innovation required to achieve efficient internalization of liposomes by LCs. Previous *ex vivo* studies have explored the use of natural glycans such as Le^Y^ for this purpose^29^. Interestingly, Le^Y^ did not promote endocytosis by LCs while the Le^X^-mediated (K_D,DC-SIGN_ = ca. 1 mM) targeting of DC-SIGN on dermal DCs succeeded^52^. At the time, the authors concluded that liposomal formulations are not suitable to address LCs. Here, we propose the concept of CLR-specific avidity thresholds to explain these findings. The affinities of the utilized natural glycans for Langerin and DC-SIGN were comparable. Yet, the targeted liposomes presumably displayed an increased avidity for dermal DCs due the tetrameric organization of the carbohydrate recognition domains, the formation of nanoclusters or increased expression levels for DC-SIGN^56^.

These characteristics renders LCs more difficult targets for glycan-mediated liposomal delivery compared to dermal DCs. Here, we have demonstrated that this difficulty can be overcome by glycomimetic ligand design. While generally considered challenging in itself, the design of mono- or oligosaccharide analogs has been successfully applied to target delivery platforms to other GBPs such as ASGPR and Siglec-2^57,58^. The 42-fold affinity increase over natural glycans observed for **15**, by proxy of model ligand **16**, exceeds that reported for other first-generation glycomimetics and highlights the success of our heparin-inspired rational design strategy^58–60^. Notably, **15** provides improved synthetic feasibility and metabolic stability over sulfated heparin-derived mono- (K_I_ = 0.28±0.06 mM) or trisaccharides (K_D_ = 0.49±0.05 mM) which display similar affinities^39^. The SAR obtained for the Glc scaffold suggests that the formation β-glucosides represents the superior conjugation strategy compared to the use of α-mannosides previously explored^50^.

Using NMR spectroscopy and molecular docking, we have proposed a Ca^2+^-dependent binding mode for **15** which likely resembles that of GlcNAc^38^. The conserved orientation of the Glc scaffold allows for the formation of π-π interactions between the phenyl ring and F315 as well as a hydrogen bond between the sulfonamide linker and N307. We conclude that these interactions contribute substantially to the affinity increase observed for **15**. The combination of the obtained SAR with our binding mode analysis will inform the design of next-generation glycomimetics. Attractive approaches to further optimize the affinity for Langerin include the introduction of sulfonamide substituents in C6 to mimic the interactions formed by GlcNS-6-OS and the introduction of electron-donating substituents on the phenyl ring^61^. Notably, our analysis does not account for conformational flexibility of the carbohydrate binding site and our analysis is furthermore limited by an incomplete resonance assignment for Langerin. Hence, X-ray crystallography will serve to validate the proposed binding mode for **15** moving forward.

In summary, liposomes bearing targeting ligand **15** were efficiently internalized both by model cells expressing Langerin as well as LCs in whole skin suspensions. Furthermore, we observed no cytotoxicity even upon exposure over several days. The kinetics of endocytosis were fast and the majority of LCs was successfully addressed within 20 minutes while internalization by off-target cells was negligible. Notably, the epidermis predominantly consists of keratinocytes while LCs only amount to approximately 2% of epidermal cells^62^. Additional skin-resident immune cells such as dermal DCs and macrophages are present in the dermis and many of these off-target cells express GBPs including CLRs such as MR, Dectin-1 and DC-SIGN or Siglec-10^55^. Accordingly, the required specific delivery of antigens or adjuvants to LCs in the human skin is particularly challenging. Yet, **15** provided remarkable specificity for Langerin, despite its low complexity. While residual off-target affinity can be expected for glycomimetic ligands, the proposed avidity threshold for liposomal targeting likely prevents endocytosis by non-LC skin-resident cells^63^. Intriguingly, this observation might be leveraged to infer general design principles for specific nanoparticle-based delivery platforms. Overall, our findings highlight not only the therapeutic potential of the targeted liposomes but also their value as molecular probes for basic research where they will potentially contribute to studying the role of LCs in skin homeostasis or to elucidate the mechanisms of antigen cross-presentation^64^.

In contrast to other CLRs, Langerin-dependent signaling has not been reported to date^65^. Our findings support this hypothesis as the binding and endocytosis of targeted liposomes did not activate immature LCs *ex vivo*. This expands the therapeutic scope of the liposomal delivery platform. On the one hand, the co-administration of adjuvants, preferably TLR-3 or MDA5 agonists, will promote the induction of cytotoxic T cell immunity required for cancer vaccines^25,66,67^. As both are intracellular pattern recognition receptors, facilitating the internalization of these agonists is of particular importance. On the other hand, antigen delivery to LCs in absence of adjuvants has been shown to result in the expansion of regulatory T cells and can be leveraged to treat autoimmune diseases^68^. In this context, liposomes are superior to antibody-antigen conjugates as they enable the co-formulation of antigens and adjuvants. While the systemic administration of adjuvants generally induces adverse effects and compromises cytotoxic T cell immunity, their targeted delivery to LCs allows for reduced adjuvant doses and tailored immune responses.

Moreover, many CLRs including Langerin recycle between the plasma membrane and the endosomal compartment^33^. In this context, the Ca^2+^- and pH-dependent release of glycomimetic ligands in the early endosome increases the endocytic capacity of LCs and other DCs^48^. It can be argued that this intracellular trafficking mechanism evolved to promote antigen cross-presentation^69^. The observed fast internalization kinetics and the prolonged co-localization of targeted liposomes with early endosomal markers suggest that this mechanism is efficiently exploited. By contrast, antibodies have been demonstrated to recycle back to the plasma membrane, thereby limiting the dose of internalized and processed antigens^20,21^. These characteristics highlight another potential advantage of the liposomal delivery platform over antibody-based approaches.

Future investigations will advance the developed liposomal delivery platform for LCs towards *in vivo* studies. Here, it will be essential to demonstrate the LC-mediated induction of cytotoxic T cell responses. Important parameters of this process that remain to be investigated are the intracellular trafficking and the efficient cross-presentation of delivered antigen. Finally, the feasibility of transcutaneous vaccinations using targeted liposomes will be evaluated to pave the way for therapeutic applications.

## Methods

Methods, including statements of data availability and any associated accession codes and references, are available in the online version of the paper.

### Synthetic Chemistry

Synthetic procedures and characterization for monosaccharide analogs and glycolipids are provided as **Supplementary Information**.

### Receptor Expression and Purification

#### General remarks

Codon-optimized genes for the expression of Langerin and DC-SIGN in *E. coli* were purchased from GenScript and Life Technologies, respectively. All growth media or chemicals used for receptor expression and purification were purchased from Carl Roth if not stated otherwise.

#### Langerin extracellular domain

Expression and purification were conducted as previously published^50^. Briefly, the trimeric Langerin extracellular domain (ECD) was expressed insolubly in *E. coli* BL21* (DE3) (Invitrogen). Following enzymatic cell lysis, inclusion bodies were harvested and subsequently solubilized. The sample was centrifuged and the Langerin ECD was refolded overnight via rapid dilution. Next, the sample was dialyzed overnight, centrifuged and purified via mannan-agarose affinity chromatography (Sigma Aldrich). For ^19^F R_2_-filtered NMR and lipid-enzyme-linked lectin assay (Lipid-ELLA) experiments, the buffer was exchanged to 25 mM Tris with 150 mM NaCl and 5 mM CaCl_2_ at pH 7.8 using 7 kDa size-exclusion desalting columns (Thermo Scientific). For STD NMR experiments, Langerin ECD samples were dialyzed five times for at least 8 h against H_2_O. Subsequently, the H2O was removed via lyophilization and the residue was stored at −80° C. Prior to STD NMR experiments, the Langerin ECD was dissolved in in 25 mM Tris-d11 (Eurisotope) with 100% D2O, 150 mM NaCl and 5 mM CaCl2 at pH 7. The concentration of Langerin ECD was determined via UV spectroscopy (A_280, 0.1%_ = 2.45). Purity and monodispersity of Langerin ECD samples were analyzed via SDS PAGE and DLS.

#### Langerin and DC-SIGN carbohydrate recognition domain

Expression and purification were conducted as previously published. Briefly, the monomeric ^15^N-labeled Langerin and DC-SIGN carbohydrate recognition domains (CRDs) were expressed insolubly in *E. coli* BL21* (DE3) (Invitrogen). Following enzymatic cell lysis, inclusion bodies were harvested and subsequently solubilized. The sample was centrifuged and the Langerin and DC-SIGN CRDs were refolded overnight via rapid dilution. Next, the sample was dialyzed overnight, centrifuged and purified via StrepTactin affinity chromatography (Iba). After an additional dialysis step overnight, the sample was centrifuged and the buffer was exchanged to 25 mM HEPES with 150 mM NaCl at pH 7.0 using 7 kDa size-exclusion desalting columns (Thermo Scientific) for ^19^F R_2_-filtered and ^15^N HSQC NMR experiments. The concentration of Langerin and DC-SIGN CRDs was determined via UV spectroscopy (A_280, 0.1%_ = 3.19 and A_280, 0.1%_ = 2.98). Purity and monodispersity of Langerin and DC-SIGN CRD samples were analyzed via SDS PAGE and DLS.

### ^19^F R_2_-filtered NMR

General remarks. ^19^F R_2_-filtered NMR experiments were conducted on a PremiumCompact 600 MHz spectrometer (Agilent). Spectra were processed in MestReNova and data analysis was performed with OriginPro^70,71^. Experiments with the Langerin ECD were performed at a receptor concentration of 50 μM in 25 mM Tris with 10% D_2_O, 150 mM NaCl and 5 mM CaCl_2_ at pH 7.8 and 25° C. Experiments with the DC-SIGN CRD were performed at a receptor concentration of 50 μM in 25 mM HEPES with 10% D_2_O, 150 mM NaCl and 5 mM CaCl_2_ at pH 7.0 and 25°C. TFA served as an internal reference at a concentration of 50 μM. Apparent relaxation rates R_2,obs_ for the reporter ligand were determined using the CPMG pulse sequence as previously published^50,72,73^.

#### Assay development for DC-SIGN

The ^19^F R_2_-filtered NMR reporter displacement assay for DC-SIGN was developed following the procedure previously published for Langerin^50^. Briefly, the K_D_ value and the relaxation rate in bound state R_2,b_ were determined at five concentrations [L]_T_ of reporter ligand **24** in three independent titration experiments. Samples were prepared via serial dilution. The addition of 10 mM EDTA served to validate the Ca^2+^-dependency of the interaction between DC-SIGN and the reporter ligand. To ensure the validity of the equations for K_D_ and K_I_ determination, the chemical exchange contribution R2,ex was estimated by ^19^F NMR relaxation dispersion experiments at a reporter ligand concentration of 0.1 mM in presence of receptor.

#### K_I_ determination

KI values were determined as previously published for Langerin^50^. Briefly, titration experiments were conducted at a concentration of 0.1 mM of reporter ligand **24** at five competitor concentrations [I]_T_. Samples were prepared via serial dilution. For the acids GlcNS, GlcNAc-6-OS and GlcNS-6-OS the pH values were monitored and adjusted to 7.8 if necessary.

### ^15^N HSQC NMR

#### General remarks

^15^N HSQC NMR experiments were conducted on an Ascend 700 MHz spectrometer (Bruker)^74^. Spectra were processed in NMRPipe^75^. Data analysis was performed using CCPN Analysis, MatLab and OriginPro^71,76,77^. Experiments with the Langerin CRD were performed at a receptor concentration of 100 μM in 25 mM HEPES with 10% D_2_O, 150 mM NaCl and 5 mM CaCl_2_ at pH 7.8 and 25° C. DSS-*d6* served as an internal reference at a concentration of 100 μM. Spectra were referenced via the internal spectrometer reference. Spectra were acquired with 128 increments and 32 scans per increments for 150 μl samples in 3 mm sample tubes. The relaxation delay d1 was set to 1.4 s and the acquisition time t_acq_ was set to 100 ms. The W5 Watergate pulse sequence was used for solvent suppression^78^. The used resonance assignment for the Langerin CRD has been published previously^32^.

#### K_D_ determination

KD values were determined in titration experiments at six ligand concentrations [L]_T_. Samples were prepared via serial dilution. Chemical shift perturbations CSPs for Langerin CRD resonances in the fast or fast-to-intermediate exchange regime observed upon titration with ligand were calculated via Equation 1^79^.

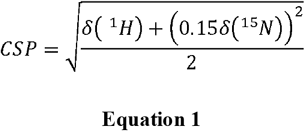

A standard deviation σ of 0.02 ppm was previously determined for the measurement of chemical shifts in ^15^N HSQC NMR experiments with the Langerin CRD^32^. Accordingly, only assigned resonances that displayed CSP values higher than a threshold of 2σ at the highest ligand concentration were selected for the determination of KD values via Equation 2 in a global two parameter fit^79^. Standard errors were derived directly from the fitting procedures. Additionally, resonances that displayed line broadening Δv_0.5_ larger than 10 Hz upon titration in either the ^1^H or the ^15^N dimension were not considered for the determination of K_D_ values. CSPmax represents the CSP value observed upon saturation of the binding site.

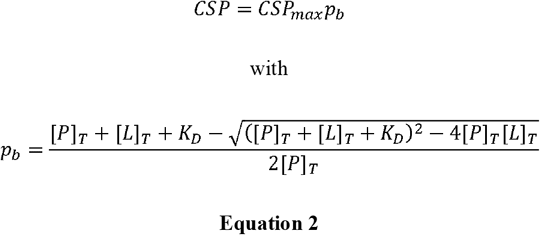

For resonances assumed to be in the slow exchange regime upon titration, K**D** values were derived from integrals Vb and Vf corresponding to the bound and free state of the Langerin CRD, respectively. V values served to calculate the bound fraction of the receptor pb via Equation 3. Integrals V were normalized via integral V of the *N*-terminal K347 and served to calculate the bound fraction of the receptor p_b_ via Equation 3. For these calculations, only resonances for which the bound state could be assigned were considered. Selected data points displaying a low SNR or issues with the baseline correction were treated as outliers and not considered for the determination of pb values. Next, a one parameter fit of Equation 3 to mean pb values served to determine KD values.

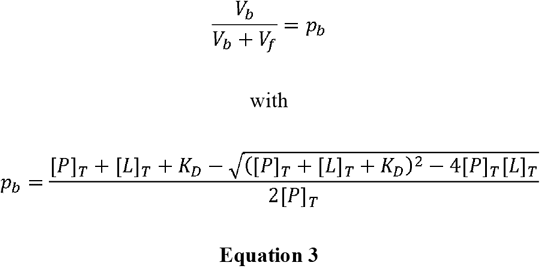

#### Binding mode analysis

Based on the resonance assignment, CSP values observed at maximal ligand concentrations [L]_T_ were mapped on the X-ray structure of the Langerin CRD (PDB code: 4N32) using Matlab’s Bioinformatics Toolbox via substitution of the B-factor values^38,80^. The CSP patterns obtained were visualized in MOE using Chain B of the Langerin CRD in complex with GlcNAc^81^. Model quality was maintained using MOE’s Structure Preparation followed by the simulation of protonation states and the hydrogen bond network of the complex with MOE’s Protonate 3D. Receptor surfaces were visualized in Connolly representation^82^.

### STD NMR

#### General remarks

STD NMR experiments were conducted on a PremiumCompact 600 MHz spectrometer (Agilent)^83^. Spectra were processed in MestReNova and data analysis was performed with OriginPro^70,71^. Experiments with the Langerin ECD were conducted at a receptor concentration of 50 µM in 25 mM Tris-d_11_ (Eurisotope) with 100% D_2_O, 150 mM NaCl and 5 mM CaCl_2_ at pH 7.8 and 25° C. Experiments were repeated in absence of receptor to exclude STD effects due to direct saturation of ligands. Residual H_2_O or TSP-*d*_*6*_ at 0.1 mM served as an internal reference. Spectra were recorded in 5 mm sample tubes at sample volumes of 500 μl. Saturation was implemented via a train of 50 ms Gauss pulses at varying saturation times t_sat_. The on-resonance irradiation frequency ν_sat_ was set to 0.0 ppm and the off-resonance irradiation frequency ν_ref_ was set to 80.0 ppm. The acquisition time t_acq_ was set to 2.0 s and the DPFGSE pulse sequence was utilized for solvent suppression^84^. Receptor resonances were suppressed using a T_1,rho_ filter at a relaxation time τ of 35 ms.

#### Epitope mapping

The binding epitope for **16** was determined at a concentrations of 500 µM. For each spectrum 512 scans were recorded. The relaxation delay d_1_ was set to 6 s and spectra were recorded at 5 different saturation time t_sat_ varying from 0.25 to 6.00 s. Equation 4 served to derive the STD effect STD for each analyzed resonance from the corresponding on- and off-resonance spectra^85^. I_0_ represents the integral of a resonance in the off-resonance spectrum and I_sat_ represents the integral of a resonance in the on-resonance spectrum.

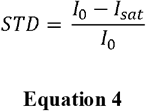

The apparent saturation rate k_sat_ and the maximal STD effect STD_max_ were derived from Equation 5 in a two parameter fit^86^. Standard errors were derived directly from the fitting procedures. These parameters were used to calculate the initial slope of the STD build-up curves STD’_0_ via Equation 6. STD’_0_ values were normalized and mapped on the corresponding ligand structure. Only resonances for which at least part of a multiplet was isolated were considered for the epitope mapping.

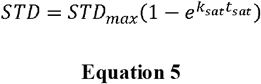

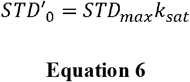

### Molecular Modelling

#### General remarks

Molecular modelling procedures were performed in MOE^81^. Deviations from default options and parameters are noted. The AMBER10:EHT force field was selected for the refinement of docking poses and the hydrogen bond network while the MMFF94x force field was utilized for the generation conformers^87–89^. Receptor surfaces were visualized in Connolly representation^82^.

#### Development of the pharmacophore model and preparation of the Langerin complex

A structural alignment of Langerin carbohydrate binding sites in complex with GlcNAc was performed (PDB codes: 4N32)^38^. Based on this visualization, a pharmacophore model was defined with features for O3, O4 and O5 of the Glc scaffold. The spatial constraint on the O3 and O4 was defined by a sphere with a radius r of 0.5 Å while the position of O5 was constrained by a sphere with a radius r of 1.0 Å. Chain B of the Langerin CRD in complex with GlcNAc served as the structural basis for the conducted molecular docking study. Additionally, an alternative conformation for K313 observed for the Langerin complex with Gal-6-OS was modeled and included in the study^90^. Overall model quality and protein geometry were evaluated and maintained using MOE’s Structure Preparation. Next, protonation states and the hydrogen bond network of the complex were simulated with MOE’s Protonate 3D followed by the removal of all solvent molecules.

#### Molecular docking

Conformations for **16** were generated utilizing MOE’s Conformation Import. A pharmacophore-based placement method was utilized to generate docking poses that we scored using the London ΔG function. Highly scored poses were refined utilizing molecular mechanics simulations, rescored via the GBIV/WSA ΔG function, filtered using the pharmacophore model and written into the output database^91^. Conformational flexibility of the carbohydrate binding site was accounted for by introducing B-factor-derived tethers to side chain atoms. Refined docking poses were ranked according to their the GBIV/WSA ΔG score and evaluated visually in the context of the conducted ^15^N HSQC and STD NMR experiments.

### Liposomal Formulation

PEGylated liposomes were prepared via thin film hydration and subsequent pore extrusion as previously published^92^. Briefly, non-targeted liposomes were formulated from a mixture of DSPC:cholesterol:PEG-DSPE:Alexa Fluor 647-PEG-DSPE (57:38:4.75:0.25). For targeted liposomes, the PEG-DSPE was substituted with glycolipids **22** or **23** at varying ratios. PEG-DSPE (3.0 kDa, NOF Europe), **22** or **23** were dissolved in DMSO, added to a round bottom flask and lyophilized. Next, DSPC (NOF Europe) and cholesterol (Sigma Aldrich) were dissolved in chloroform, added to the test tube and the solvent was removed *in vacuo*. The residue was dissolved in PBS and the mixture was vortexed and sonicated repeatedly until a homogeneous suspension was obtained. Unilamellar liposomes were prepared using a pore extruder (Avanti Polar Lipids) with polycarbonate membranes of 800, 400, 200 and finally 100 nm pore size (Avanti Polar Lipids). Liposomes were characterized by DLS and electrophoresis experiments to determine their Ζ potential. Liposomes were stored at 4° C. Non-targeted liposomes and liposomes **23** were characterized in ^1^H NMR experiments at a total lipid concentration [Lipid]_T_ of 1.5 mM in PBS at 25° C.

### Lipid-ELLA

1 mg•l^−1^ of PEG-DSPE (3.0 kDa, NOF Europe), **22** or **23** or unconjugated PEG-DSPE in 100 mM Tris (50 μl per well) at pH 8.9 and 4° C were added to a 384 well MaxiSorp plate (Nunc) and incubated overnight to immobilize the PEGylated lipids. After removal of the supernatant, the wells were blocked with 2% BSA in 25 mM Tris with 150 mM NaCl and 0.1% Tween-20 (70 μl per well) at pH 7.6 and room 25° C for 1 h. Next, the wells were washed three times using 25 mM HEPES with 150 mM NaCl, 5 mM CaCl_2_ and 0.01% Tween-20 at pH 7.6 and room temperature (100 μl per well). The wells were incubated with Langerin ECD in 25 mM HEPES with 150 mM NaCl, 5 mM CaCl_2_ and 0.01% Tween-20 (50 μl per well) at pH 7.6 and room temperature for 4 h at 10 different concentrations. Subsequently, the wells were washed and incubated with HRP conjugate and 2% BSA in 25 mM HEPES with 150 mM NaCl, 5 mM CaCl_2_ and 0.01% Tween-20 (50 μl) at pH 7.6 and room temperature for 45 min. The wells were washed and developed with TMB solution (Rockland) (50 μl per well). The reaction was quenched after 5 min by addition of 0.18 M sulfuric acid (50 μl per well). Binding of the Langerin ECD was detected via absorbance A_450_ measurements at 450 nm using a SpectraMax spectrometer (Molecular Devices) in three independent titrations.

### Establishment of C-type lectin^+^ model cells

#### Lentivirus production

TrueORF sequence-validated cDNA clones of human Langerin, human DC-SIGN and murine Dectin-1 (Sinobiologicals) were amplified from a pcDNA5/FRT/V5-His-TOPO TA expression vector (Life Technologies) by PCR (forward primer: 5′-CTGGCTAGCGTTTAAACTT AAG -3′, reverse primer: 5′-CAATGGTGATGGTGATGATG -3′) using Phusion polymerase (Thermo Fisher Scientific). The PCR amplicons were cloned in a BIC-PGK-Zeo-T2a-mAmetrine:EF1A construct by Gibson assembly (NEB) according to the manufacturer’s protocol. The destination vector was linearized by PCR (forward primer: 5′-GAGCTAGCAGtaTTAATTAACCACCCTGGCTAGCGTTT AAACTTAAG-3′; reverse primer: 5′-GTACCGGTTAGGATGCATGCCAATGGTGATGGTGATG ATG -3′) using Phusion Polymerase. Together with third-generation packaging vectors pVSV-G, pMDL and pRSV, the constructs were transfected into HEK293T cells (ATCC) using the Mirus LT1 reagent (Sopachem) for production of the lentivirus as previously published93. After 3 to 4 days the supernatant containing the viral particles was harvested and to −80ºC to kill any remaining HEK293T cells. This supernatant was used to transduce Langerin, DC-SIGN and Dectin-1 into Raji and COS-7 cells (ATCC).

#### Lentiviral transduction

50,000 Raji or COS-7 cells were transduced by spin infection for 2 h at 1000 g and 33°C using 100 µl virus-containing supernatant supplemented with 8 µg/ml polybrene (Santa Cruz Biotechnology). Complete medium (Thermo Fisher Scientific) was added after centrifugation. 2 to 3 days post-transduction mAmetrine expression was measured by flow cytometry to confirm integration of the construct. Cells were selected 3 days post-infection by 100 µg/ml or 200 µg/ml zeocin (Gibco) for COS-7 and Raji cells, respectively. After selection, 95% of cells were mAmetrine^+^ and the expression of Langerin, DC-SIGN or Dectin-1 was validated by flow cytometry.

### Preparation of Epidermal and Whole Skin Cell Suspensions

Healthy human skin samples were collected after informed consent and approval by the local ethics committee (AN 5003 360/5.22 of the 15.04.2016). Subcutaneous fat was removed with a scalpel. For preparation of epidermal cell suspension, skin pieces were incubated in RPMI1640 medium (Lonza) containing 1.5 U/ml dispase II (Roche, Switzerland) and 0.1% trypsin (Sigma Aldrich) overnight at 4°C. Next the epidermis was isolated, broken up into smaller pieces, and filtered through a 100 µm cell strainer (Thermo Fisher Scientific) to obtain a single cell suspension. For digestion of whole skin, skin pieces were incubated in RPMI1640 medium (Lonza) supplemented with 10% FCS (Pan-Biotech) containing 1 mg/ml collagenase IV (Worthington Biochemical Corporation) overnight at 37°C. Cell suspension was then filtered through a 100 µm cell strainer (Thermo Fisher Scientific).

### Flow Cytometry

#### C-type lectin receptor^+^ model cell lines

Raji cells were cultured in RPMI1640 medium (Sigma Aldrich) containing 10% FCS (Biochrom), 100 U*ml^−1^ penicillin and streptomycin (Life Technologies) and GlutaMax-I (Life Technologies) at 37° C and 5% CO_2_. To validate the expression of CLRs, 50,000 cells were incubated with 25 µl medium containing fluorophore-labeled primary antibodies for Langerin (clone DCGM4, Beckman Coulter), DC-SIGN (clone 9E9A8, Bio Legend) and Dectin-1 (clone BG1FPJ, eBioscience). Isotype-matched antibodies were used for control experiments. After incubation for 30 min at 4°C, cells were washed and CLR expression was analyzed by flow cytometry.

To monitor internalization and binding, liposomes and, in case of control experiments, 10 mM EDTA or 50 µg•ml^−1^ mannan in PBS were added to 96 well microtiter plates (Nunc). Cells were counted, centrifuged at 500 g for 3 min, aspirated and resuspended in culture medium at 37° C and 5% CO_2_. 50,000 cells were added to the 96 well microtiter plates (Nunc) to obtain a final volume of 100 µl. The plates were incubated for 1 h at 4° C and subsequently centrifuged at 500 g for 3 min. Cells were aspirated and resuspended in culture medium at 37° C and 5% CO_2_. Internalization and binding of liposomes were analyzed by flow cytometry. Following the initial optimization of the liposome concentration, all experiments were conducted at a total lipid concentration [Lipid]_T_ of 16 µM.

To evaluate cytotoxic effects induced by liposomes, cells were stained with the early apoptotic marker Annexin-V and the late apoptotic marker 7-AAD. 10,000 cells were incubated with at different total lipid concentration [Lipid]_T_ for 24 h or at an [Lipid]_T_ of 16 µM for various times at a final volume of 50 µl at 37°C and 5% CO_2_. Incubation with 50% DMSO for 3 min at the same condition was used as a positive control. Following incubation, the cells were washed and resuspended in 25 µl 10 mM HEPES with 140 mM NaCl and 2.5 mM CaCl_2_ containing Annexin-V-FITC (1:100 dilution, Adipogen) at pH 7.4. Next, the cells were protected from light and incubated for 10 min at room temperature. After washing, the cells were resuspended in 100 µl 10 mM HEPES with 140 mM NaCl and 2.5 mM CaCl_2_ containing 7-AAD (1:100 dilution, Bio Legend) at pH 7.4, protected from light and incubated for 5 to 10 min at room temperature. Cytotoxic effects induced by liposomes were analyzed by flow cytometry. All flow cytometry experiments were conducted using Attune Nxt Flow Cytometer equipped with an autosampler (Life Technologies) and analyzed with FlowJo^94^. Mean fluorescence intensities were determined in three independent experiments.

#### Epidermal and whole skin cell suspensions

Liposomes were incubated with epidermal or whole skin cell suspensions at a total lipid concentration [Lipid]_T_ of 16 µM in HBSS with 2 mM CaCl_2_ (Biochrom) supplemented with 1% BSA (Serva Electrophoresis) for 1h at 37° C. For experiments analyzing the kinetics of liposome internalization, epidermal cell suspensions were incubated for varying times at 4° C or 37° C and endocytosis was abrogated by addition of 10 mM EDTA. Alternatively, liposomes were incubated for varying times at 4° C or 37° C with epidermal cell suspension in presence of 10 mM EDTA. To evaluate cytotoxicity or maturation effects induced by liposomes, epidermal cell suspensions were incubated with liposomes at an [Lipid]_T_ of 2.7 µM for 1h or 48h at 37°C in RPMI1640 medium (Lonza) supplemented with 10% FCS (Pan-Biotech), 2 mM L-glutamine (Lonza), 50 µg/ml gentamicin (Gibco) and 200U/ml GM-CSF (Leukine sargramostim, Sanofi).

Flow cytometry experiments were conducted on a FACS Canto II Flow Cytometer (BD Biosciences) and analyzed in FlowJo^94^. Non-specific FcR-mediated antibody staining was blocked by human FcR Blocking reagent (Miltenyi Biotec). Dead cells were excluded by the fixable viability dye eFluor 780 (eBioscience). Fluorophore-labeled primary antibodies for CD1a (clone HI149), CD14 (clone HCD14), HLA-DR (clone L243), CD45 (clone HI30), CD83 (clone HB15) (Bio Legend), CD80 (clone L307.4, BD Biosciences) and Langerin (clone MB22-9F5, Miltenyi Biotec) or isotype-matched control antibodies were used for gating. Immune cells were always pre-gated on viable CD45^+^ cells. All antibody incubation steps were performed for 15 min at 4° C. For intracellular staining of active caspase 3, staining was done according to the manufacturer’s protocol and the cells were incubated with an antibody for active caspase 3 (clone C92-605; BD Biosciences) for 30 minutes at room temperature.

### Confocal Microscopy

#### C-type lectin receptor^+^ model cell lines

Langerin^+^ COS-7 cells were cultured in DMEM supplemented with 10% FCS (Pan-Biotech) at 5% CO_2_ and 37°C. To analyze the co-localization of liposomes **22** with endosomal markers, cells were either stained by immunofluorescence in case of EEA1 or Rab5 or transfected with a YFP-Rab9 construct. The YFP-Rab9 construct was a kind gift of Dr. Oliver Rocks (Max Delbrück Centrum, Berlin)^95^. Cells were seeded on glass coverslips in a 12-well tissue culture dish () and incubated with liposomes **22** at a total lipid concentration [Lipid]_T_ of 16 µM in DMEM at 4°C for 2h. Incubation was started either 1 day after seeding for staining by immunofluorescence or 1 day after transfection. For the initial time point, cells were washed with PBS with 2 mM MgCl_2_ and 2 mM CaCl_2_ (Invitrogen) at 4°C and fixed in 4% Roti-Histofix (Roth) for 10 min at room temperature. For the remaining time points, cells were incubated at 37°C, washed with PBS with 2 mM MgCl_2_ and 2 mM CaCl_2_ (Invitrogen) and subsequently fixed. For immunofluorescence, cells were washed 2 times with PBS and permeabilized in PBS with 0.2% Triton-X-100 and 100 mM glycine for 10 min at room temperature. After washing two times and subsequent blocking with PBS with 3% BSA for 30 min at room temperature, cells were incubated with primary antibodies for EEA1 (rabbit, clone C45B10) and Rab5 (rabbit, clone C8B1) (Cell Signaling Technology) in PBS with 3% BSA for 1h at room temperature. After washing three times for 5 min with PBS, cells were incubated with fluorophore-labeled secondary antibody (Invitrogen) for 30 min at room temperature followed by three additional washing steps. DAPI (1:1000 dilution, Thermo Fisher Scientific) in PBS was incubated for 5 min at room temperature, following by washing with PBS for 5 min at room temperature. Coverslips were mounted using Roti-Mount (Roth). For Rab9 visualization, cells were transfected 1 day after seeding with 500 ng YFP-Rab9 per well using Lipofectamine 2000 (Invitrogen) according to the manufacturer’s protocol. After 1 day, cells were incubated with liposomes and fixed as described above. After fixation, cells were washed two times with PBS and mounted. The co-localization of liposomes with endosomal markers was visualized with an SP8 confocal microscope (Leica). **Epidermal cell suspensions.** For confocal microscopy, 100,000 cells obtained from an epidermal cell suspension in 200 µl of RPMI1640 medium (Lonza) supplemented with 10% FCS (Pan-Biotech), 2 mM L-glutamine (Lonza) and 50 µg/ml gentamicin (Gibco) were incubated with FITC-labeled CD1a (clone HI149, Bio Legend) in a micro-slide 8 well (Ibidi) for 15 minutes at 4° C. Next, the cells were incubated for 1 h with liposomes **22** at a total lipid concentration [Lipid]_T_ of 16 µM at 37° C. Internalization of liposomes was visualized with an AxioObserver Z1 confocal microscope (Carl Zeiss).

## Supplementary Information

Supplementary information, including supplementary methods, is available in the online version of the paper. Reprints and permissions information is available online. Correspondence and requests for materials should be addressed to C. R..

## Statistical Analysis

For all figures, error bars reflect the standard error of the mean. For Figure 4, the repeated One-Way ANOVA Test with a post-hoc Tukey’s Test was employed for statistical analysis. The analysis was conducted independently for the 1 h and 48 h time points.

### Acknowledgements

The work was supported by the DFG (RA1944/2-1) and the Max Planck Society. We thank Prof. Dr. Peter H. Seeberger for support and helpful discussions. R.D. and N.M.S. were supported by the Dutch Scientific Organization (VIDI 91713303).

## Author contributions

Conceptualization: E.-C.W., J.S., L.B., P.S. and C.R.; Methodology: E.-C.W, J.S., L.B., G.B., F.F.F., R.D., N.M.S., O.S., P.S. and C.R.; Formal Analysis: E.-C.W, J.S., L.B., P.S. and C.R.; Investigation: E.-C.W, J.S., L.B., P.S. and C.R.; Resources: E.-C.W, J.S., L.B., G.B., F.F.F., R.D., D.H., N.M.S., O.S., P.S. and C.R.; Writing – Original Draft: E.-C.W and C.R.; Writing – Review & Editing: E.-C.W, J.S., L.B., G.B., O.S., P.S. and C.R.; Visualization: E.-C.W, J.S., L.B., P.S. and C.R.; Supervision: P.S. and C.R.; Project Administration: P.S. and C.R; Funding Acquisition: N.S., O.S., P.S. and C.R.

