## Supplementary Materials for "A specific, glycomimetic Langerin ligand for human Langerhans cell targeting" (from another version of this preprint: version 1, 2018-03-21)

Eike-Christian Wamhoff[1,2] (orcid.org/0000-0002-8665-5512), Jessica Schulze[1,2], Lydia Bellmann[3] (orcid.org/0000-0002-1891-5319), Gunnar Bachem[4], Felix F. Fuchsberger[1,7], Juliane Rademacher[1], Martin Hermann[5] (orcid.org/0000-0003-2213-3448), Barbara Del Frari[6], Rob van Dalen[7] (orcid.org/0000-0002-0436-6048), David Hartmann[1], Nina M. van Sorge[7] (orcig.org/0000-0002-2695-5863), Oliver Seitz[4], Patrizia Stoitzner[3] (orcid.org/0000-0002-8488-6704) and Christoph Rademacher[1,2]* (orcid.org/0000-0001-7082-7239)

### Supplementary Information

| | |
|---|---|
| **Supplementary Notes** | **S2** |
| **Supplementary Schemes** | **S3** |
| **Supplementary Figures** | **S6** |
| **Supplementary Tables** | **S18** |
| **Supplementary Methods** | **S21** |
| **Supplementary References** | **S45** |

### Supplementary Notes

#### Supplementary Note 1

Upon titration with both GlcNS analogs **2** and **16** as well as Man analog **21**, additional CSPs were observed for residues remote from the carbohydrate binding site (Supplementary Figure 9). These residues, prominently featuring K257 and G259 in the short loop, were previously demonstrated to constitute an allosteric network involved in the regulation of $Ca^{2+}$ recognition of Langerin[1]. Hence, we hypothesize that this allosteric network might contribute to the recognition of glycans as well as glycomimetics[2].

While the CSP trajectories in the short loop were conserved between the Glc and the Man scaffold, residues associated with the carbohydrate binding site such as E285 and W252 displayed altered trajectories (Supplementary Figure 9). This observation potentially results from conserved $Ca^{2+}$-dependent interactions for both the Glc and Man scaffold which transduce allosteric effects. By contrast, changes in the chemical environment due to unique secondary interactions formed by e.g. **16** are locally constrained.

### Supplementary Schemes

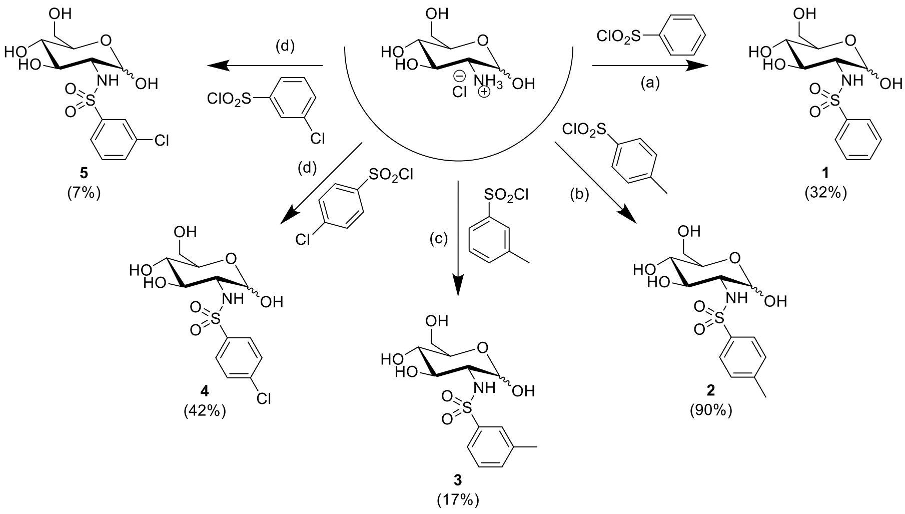


**Supplementary Scheme 1. Synthesis of GlcNS analogs 1 to 5.** Reaction conditions for the preparation of the library: (a) $Et_3N$, anhydrous MeOH, 0°C to room temperature; (b) acetone, room temperature; (c) DIPEA, DMSO, room temperature; (d) DIPEA, anhydrous MeOH, room temperature.

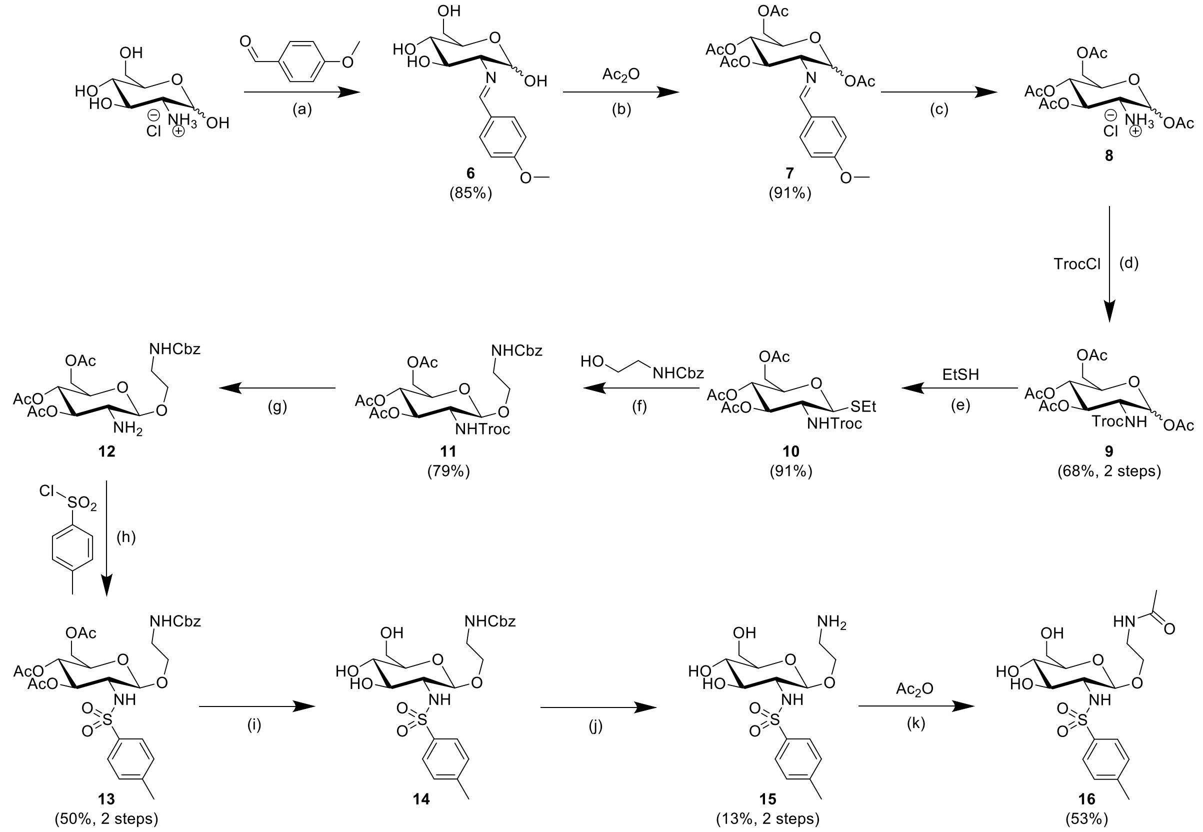


**Supplementary Scheme 2. Synthesis of GlcNS analogs 15 and 16.** Intermediate **10** was prepared as previously published[3]. Reaction conditions for the preparation of **16**: (a) 1 M aqueous NaOH, 0°C; (b) pyridine, 0°C to room temperature; (c) 5 M aqueous HCl, acetone, reflux; (d) pyridine, 0°C; (e) $BF_3{\cdot}OEt_2$, anhydrous DCM, 0°C to room temperature; (f) DMTST, anhydrous DCM, room temperature; (g) Zn, AcOH, room temperature; (h) pyridine, room temperature; (i) MeONa, anhydrous MeOH, room temperature; (j) $H_2$, Pd/C, EtOH, room temperature; (k) MeOH, room temperature.

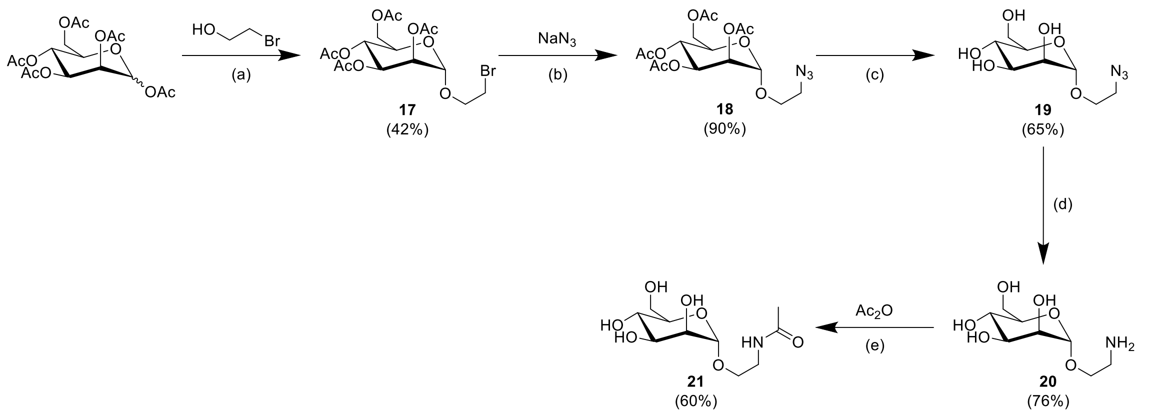


**Supplementary Scheme 3. Synthesis of Man analogs 20 and 21. 20** was prepared as previously published[4]. Reaction conditions for the preparation of **21**: (a) $BF_3{\cdot}OEt_2$, anhydrous DCM, room temperature; (b) anhydrous DMF, 60°C; (c) MeONa, MeOH, room temperature; (d) $H_2$, Pd/C, anhydrous MeOH, room temperature; (e) MeOH, room temperature.

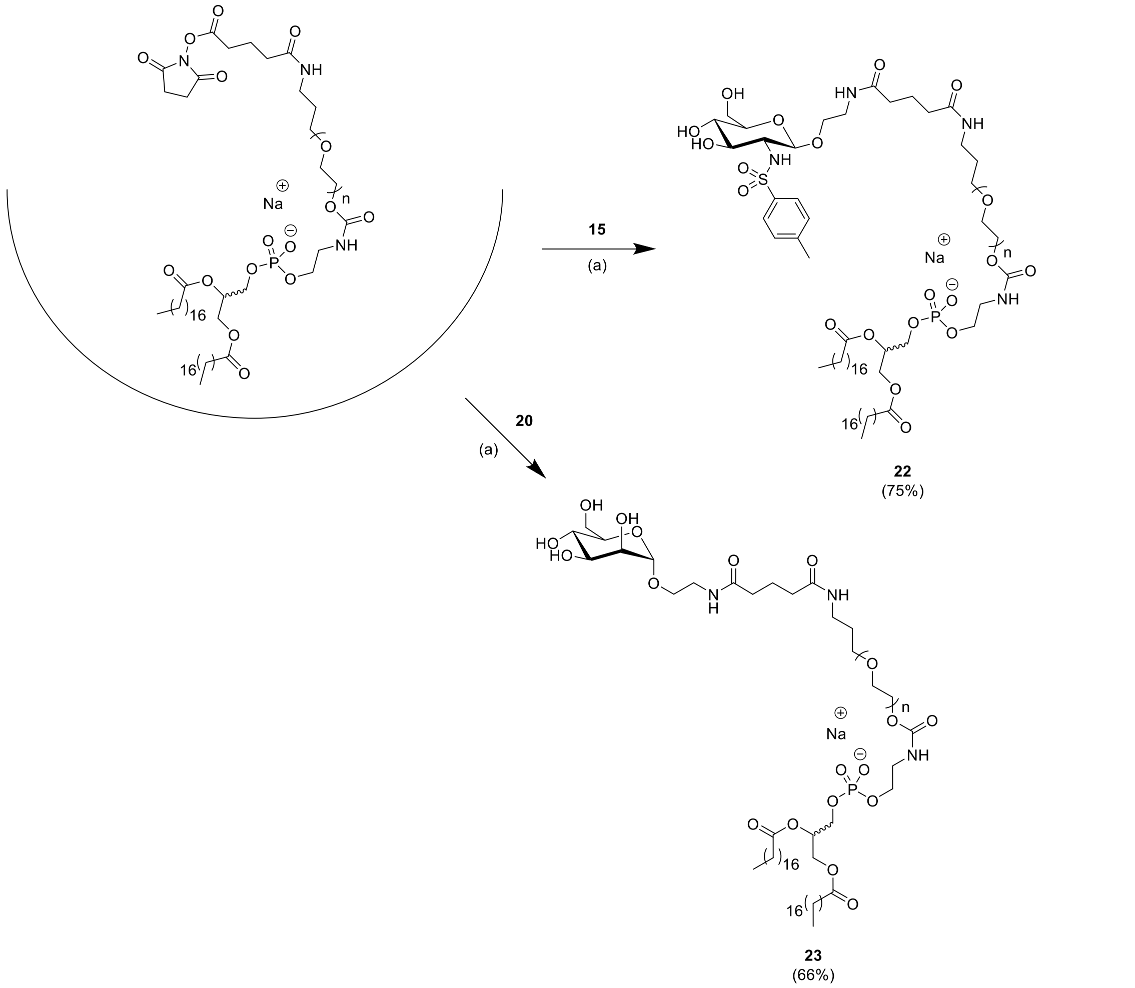


**Supplementary Scheme 4. Synthesis of glycolipids 22 and 23.** Reaction conditions for the preparation of **42** and **43**: (a) DMF:1M aqueous $NaHCO_3$ (1:10), pH 8.4, room temperature.

### Supplementary Figures

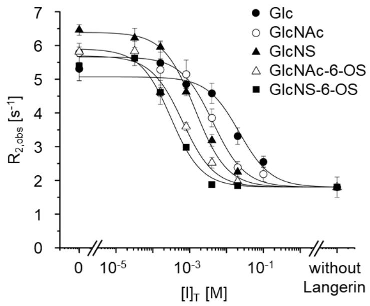


**Supplementary Figure 1. $K_I$ determination for sulfated GlcNAc derivatives.** The $K_I$ determination for heparin-derived GlcNAc derivatives *via* $^{19}F$ $R_2$-filtered NMR revealed the impact of sulfation patterns on monosaccharide affinity. Obtained $K_I$ values are given in Supplementary Table 1.

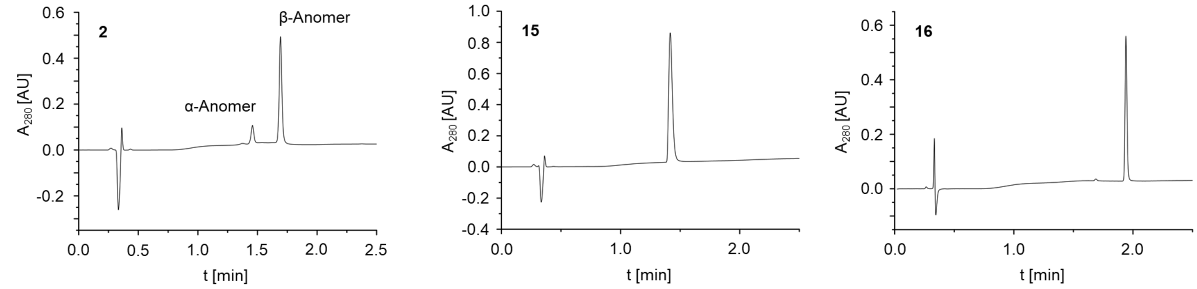


**Supplementary Figure 2. Analytical HPLC traces for GlcNS analogs 2, 15 and 16.** The purity of **2**, **15** and **16** was determined to be >95% using analytical reversed-phase HPLC (gradient: 3% to 30% acetonitrile in 0.1% TFA in $H_2O$ in 2.5 min). The assignment of the two peaks for **2** to the α- and the β-anomer were validated via $^1H$ NMR and ESI MS.

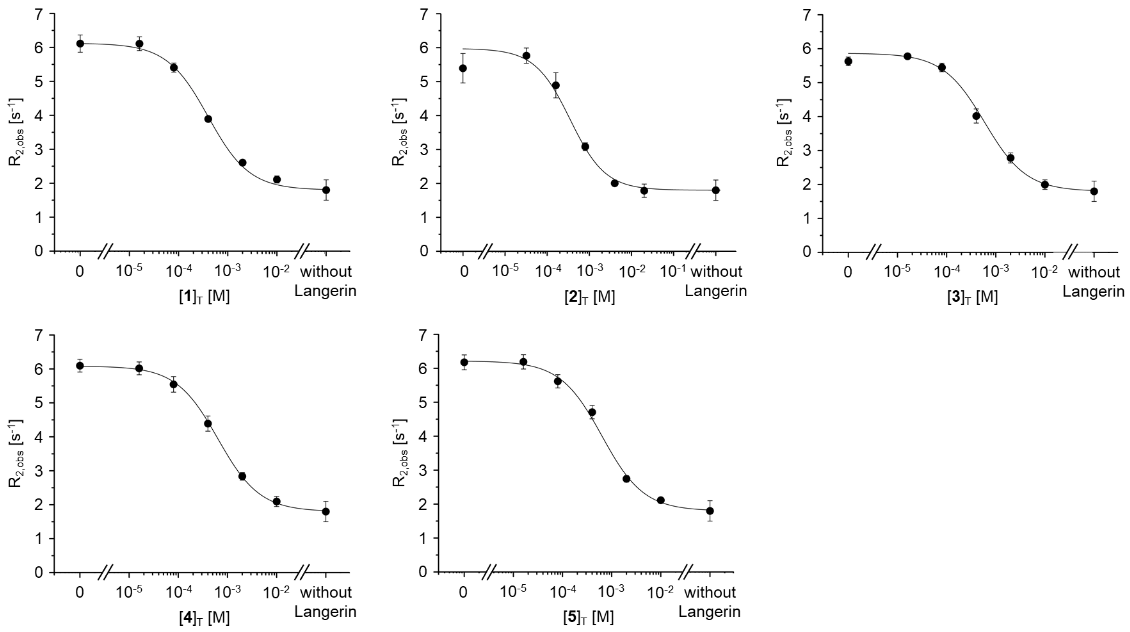


**Supplementary Figure 3. $K_I$ determination for GlcNS analogs 1 to 5.** Competitive binding experiments served to determine the affinities for the GlcNS analog library. Obtained $K_I$ values are given in Supplementary Table 2.

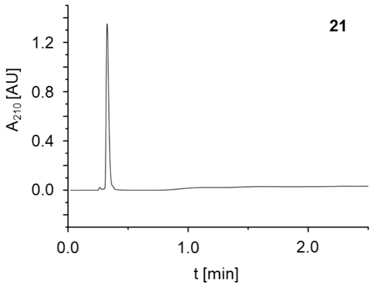


**Supplementary Figure 4. Analytical HPLC trace for Man analog 21.** The purity of **21** was determined to be >95% using analytical reversed-phase HPLC (gradient: 3% to 30% acetonitrile in 0.1% TFA in $H_2O$ in 2.5 min).

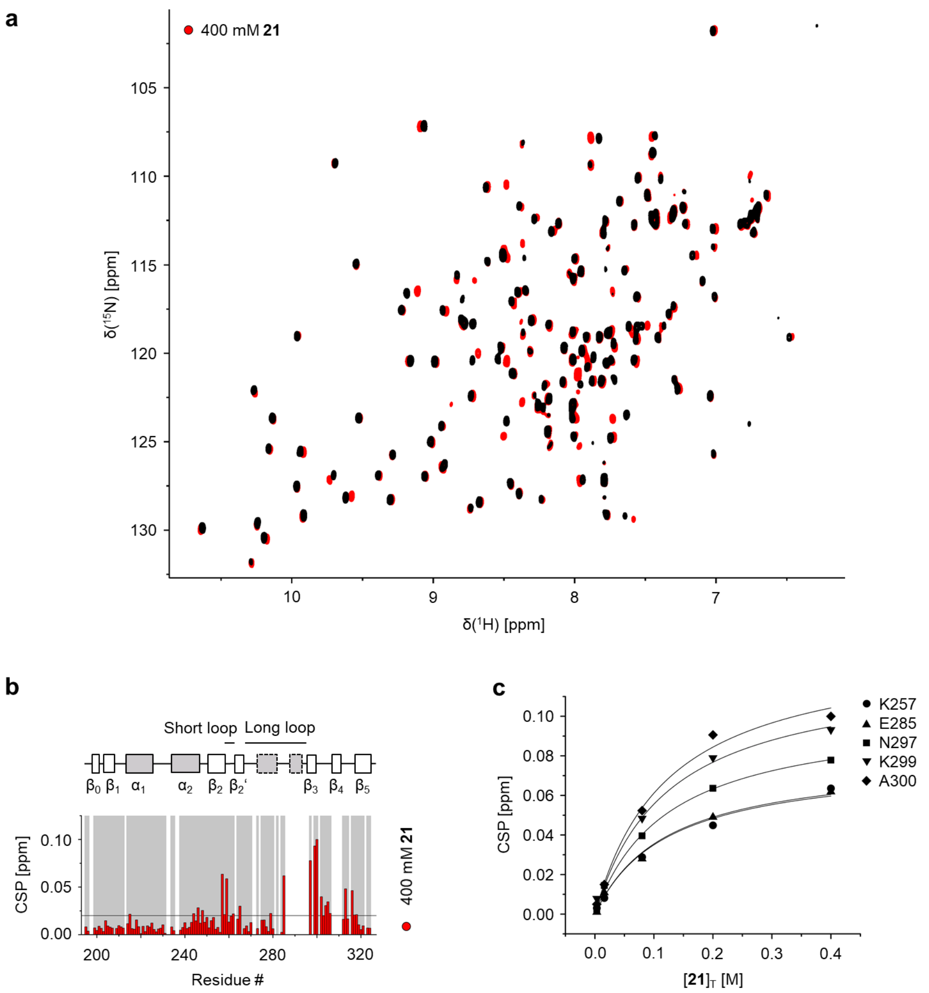


**Supplementary Figure 5. $K_D$ determination for Man analog 21. a and b.** $^{15}N$ HSQC NMR experiments served to validate the obtained $K_I$ value for **2**. Assigned resonances detected in the reference spectrum are highlighted (grey). **c.** Assigned resonances displaying fast chemical exchange and CSPs larger than 0.06 ppm were selected for the determination of $K_D$ values. The obtained $K_D$ value is given in Table 1.

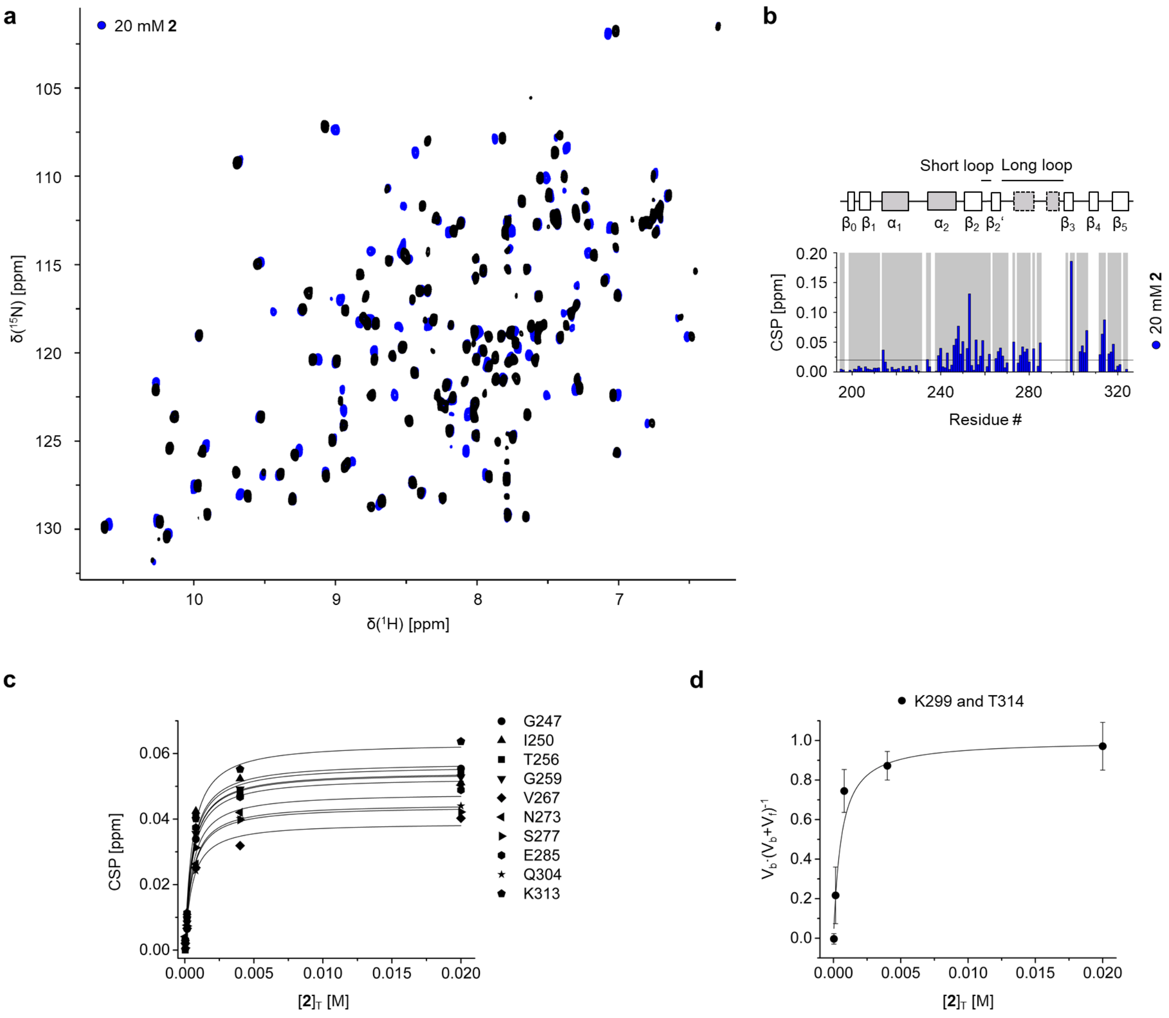


**Supplementary Figure 6. $K_D$ determination for GlcNS analog 2. a and b.** $^{15}$N HSQC NMR experiments served to validate the obtained $K_I$ value for **2**. Assigned resonances detected in the reference spectrum are highlighted (grey) **c.** Assigned resonances displaying fast chemical exchange and CSPs larger than 0.04 ppm were selected for the determination of $K_D$ values. **d.** Additionally, a set of residues including K299 and T314 displayed slow exchange phenomena. For these residues, integrals $V_f$ and $V_b$ of resonances corresponding to the free and the bound state of the Langerin were utilized to determine $K_D$ values. Obtained $K_D$ values are given in Table 1.

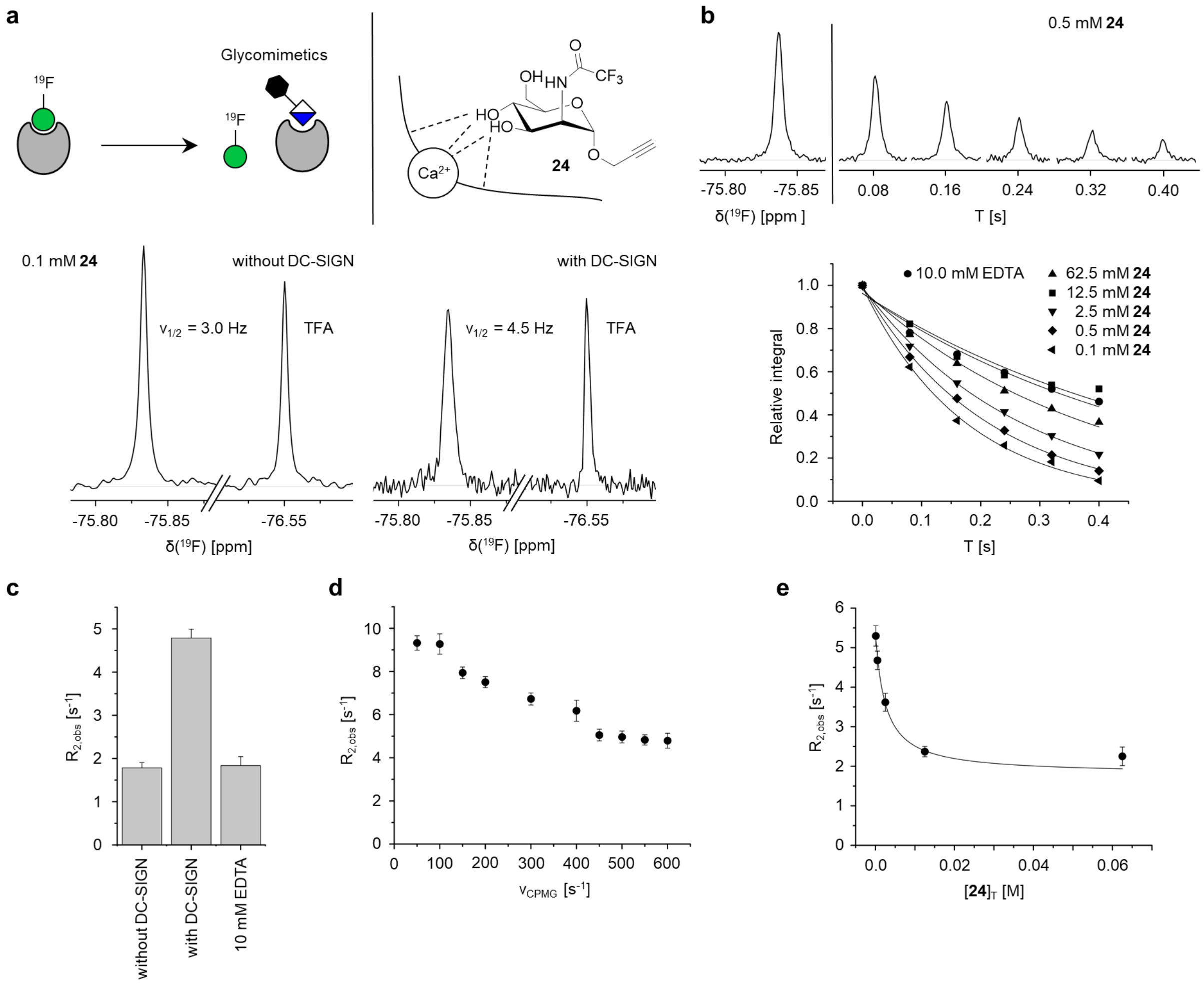


**Supplementary Figure 7. $^{19}$F $R_2$-filtered NMR assay development with DC-SIGN. a.** In presence of DC-SIGN, the $^{19}$F NMR resonance of the trifluoroacetamido group of reporter ligand **24** displayed substantial line broadening. This phenomenon was utilized to develop the $^{19}$F $R_2$-filtered NMR reporter displacement assay following the procedure previously published for Langerin[5]. The spectrum in absence of DC-SIGN is processed with an exponential apodization function at 1.8 Hz. **b.** The interaction between **24** and DC-SIGN was quantified via the transversal relaxation rate $R_{2,obs}$ using the CPMG pulse sequence[6,7]. Representative decay curves in the presence of DC-SIGN are shown. **c.** The $Ca^{2+}$-dependency of the interaction was validated via addition of EDTA. The $R_{2,obs}$ value in absence of DC-SIGN was determined from four independent experiments. The standard error for the experiments in presence of DC-SIGN and EDTA is derived directly from the fitting procedure. **d.** Relaxation dispersion experiments for **24** in presence of DC-SIGN indicate at negligible exchange contribution $R_{2,ex}$ at a $\nu_{CPMG}$ value of 500 Hz. In contrast to Langerin, a considerable exchange contribution is observed a lower $\nu_{CPMG}$ values. **e.** A representative binding isotherm for the affinity determination for **24** is shown. The $K_D$ value was determined from three independent titrations. The obtained $K_D$ and additional parameters from the fitting procedure are given in Supplementary Table 3.

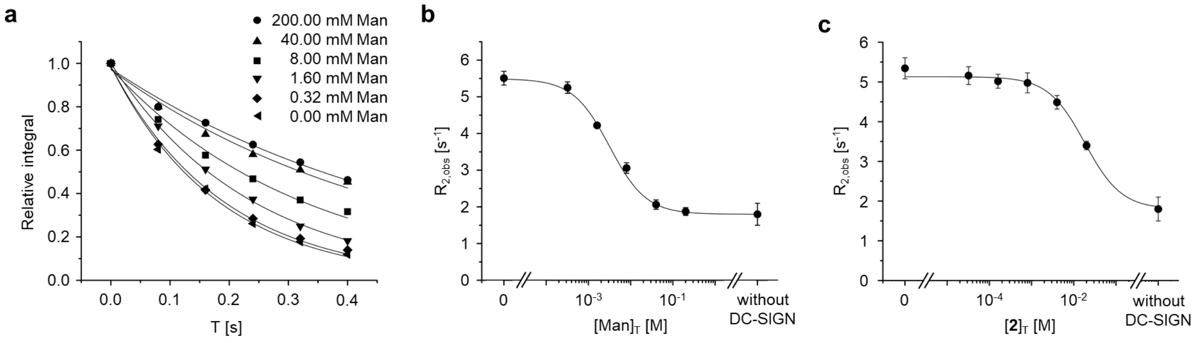


**Supplementary Figure 8. $K_I$ determination for Man and GlcNS analog 2 with DC-SIGN. a.** Representative decay curves from the competitive binding experiments with Man are shown. **b and c.** Titration with Man revealed an affinity consistent with literature values while the $K_I$ value determined for **2** indicated specificity against DC-SIGN[8]. Obtained $K_I$ values are given in Table 1.

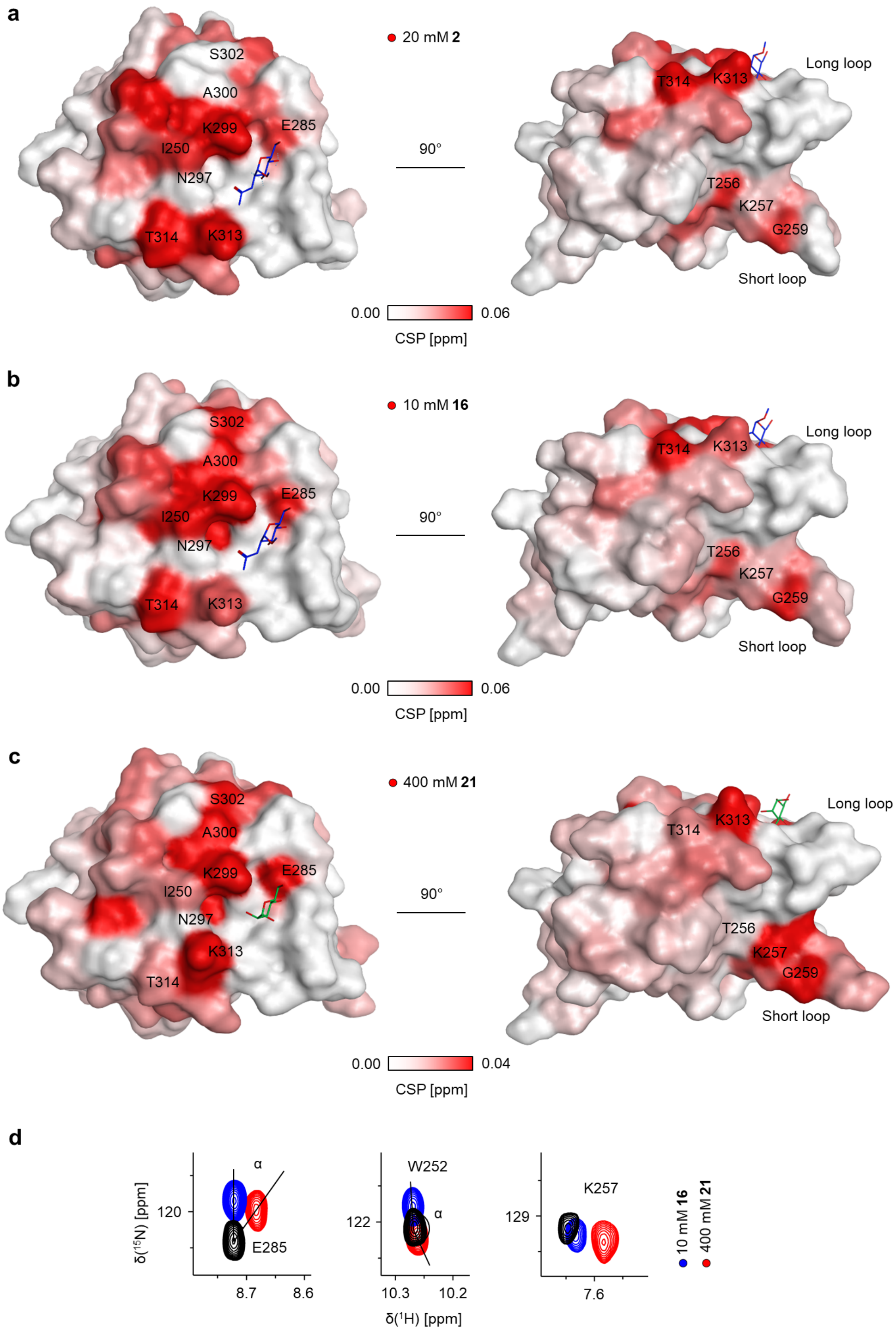


**Supplementary Figure 9. 15N HSQC NMR binding mode analysis for GlcNS analogs 2, 16 and Man analog 21. a to c.** The mapping of CSP values on the X-ray structure of Langerin (PDB code: 4N32 or 35PF) validated a $Ca^{2+}$-dependent binding mode for **2** and **16** as indicated by CSPs observed for E285 and K299[9,10]. Additionally, CSPs were observed for N297, A300 and S302 residues also affected upon recognition of Man or **21**. By contrast, Y251 and I250 displayed considerably increased CSP values compared to **21**, while a relative decrease was observed for K313. This decrease was accompanied by a relative increase for the proximal T314. Notably, residues that display considerably increased CSP values can predominantly associated with F315 and N307 which were not assigned. This also hold true for W252 and W306 that displayed smaller relative increases. Accordingly, the observed CSP pattern might be induced by interactions formed between **2** or **16** and F315 rather than K313. Similar to Man and **21**, CSPs were also observed in remote regions of the C-type lectin-like domain fold, particularly for K257 and G259 in the short loop region. This might indicate a modulation of the previously reported allosteric network[1]. **d.** A comparison of titrations with **16** and **21** revealed distinct CSP trajectories for residues associated with the carbohydrate binding site such as E285 or W252 while trajectories of residues located in remote regions of the C-type lectin-like fold such as K257 were conserved.

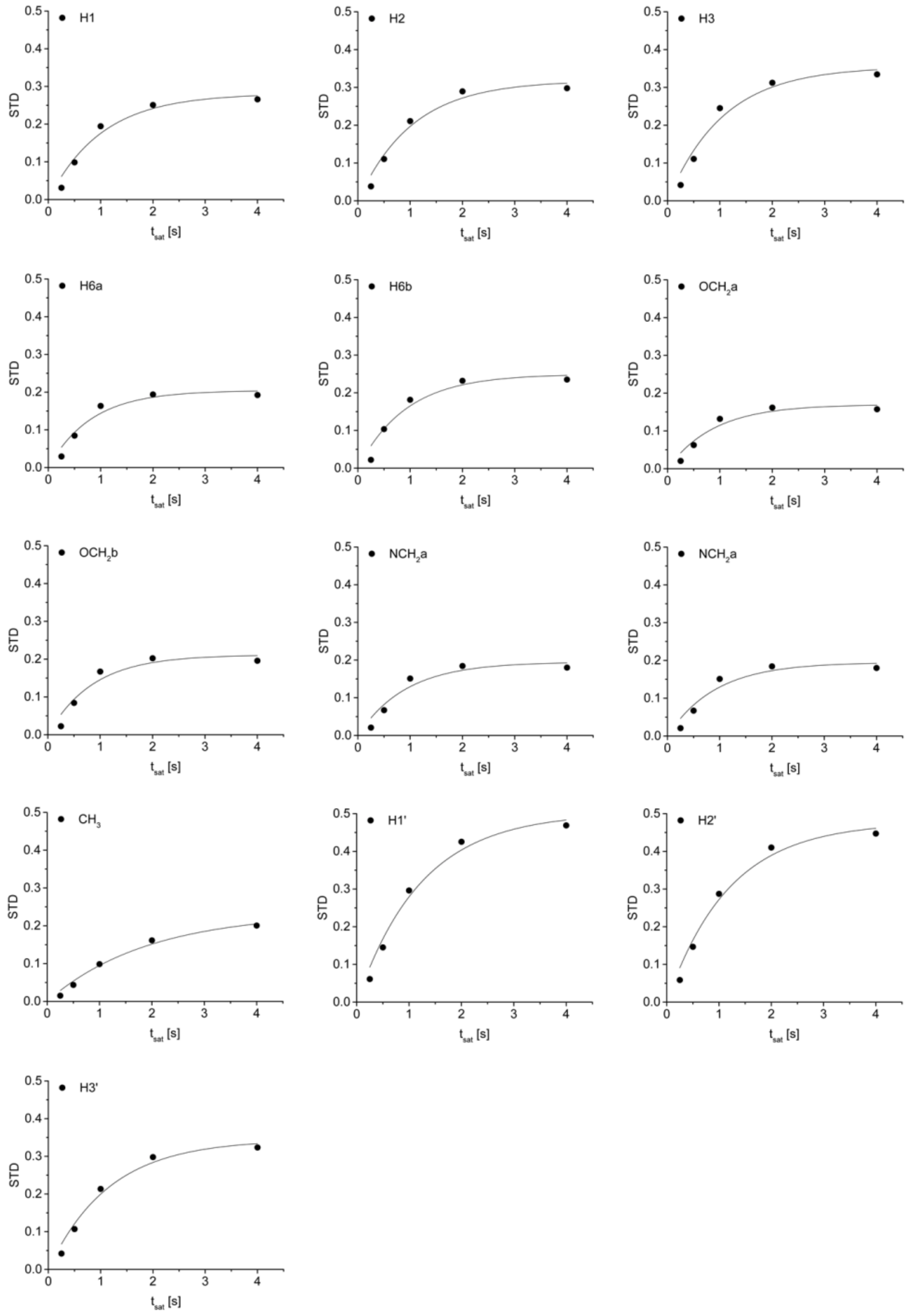


**Supplementary Figure 10. STD NMR build-up curves for GlcNS analog 16.** Equation 5 was fitted to STD values to calculate $STD_0$' values for the determination of the binding epitope of **16**.

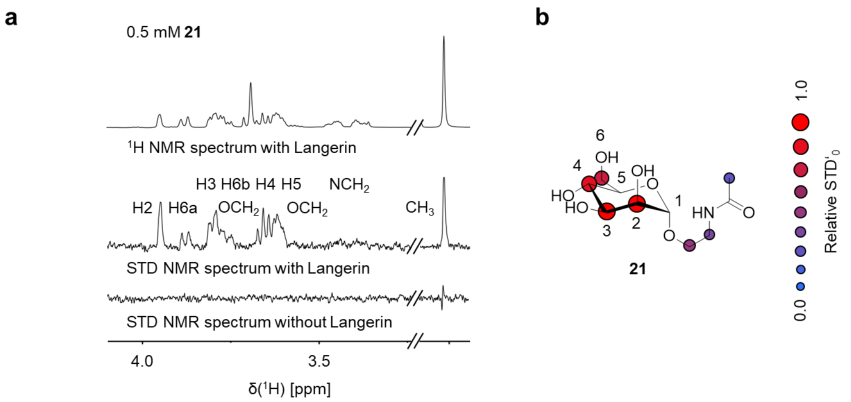


**Supplementary Figure 11. STD NMR epitope mapping for Man analog 21. a.** STD NMR experiments served to investigate the interaction of **21** with Langerin. STD NMR spectra were recorded at saturation times $t_{sat}$ of 0.4 s and are magnified 8-fold. **b.** The epitope for **21** was determined from build-up curves and suggests a solvent exposed orientation for acetylated ethylamino linker (Supplementary Figure 12).

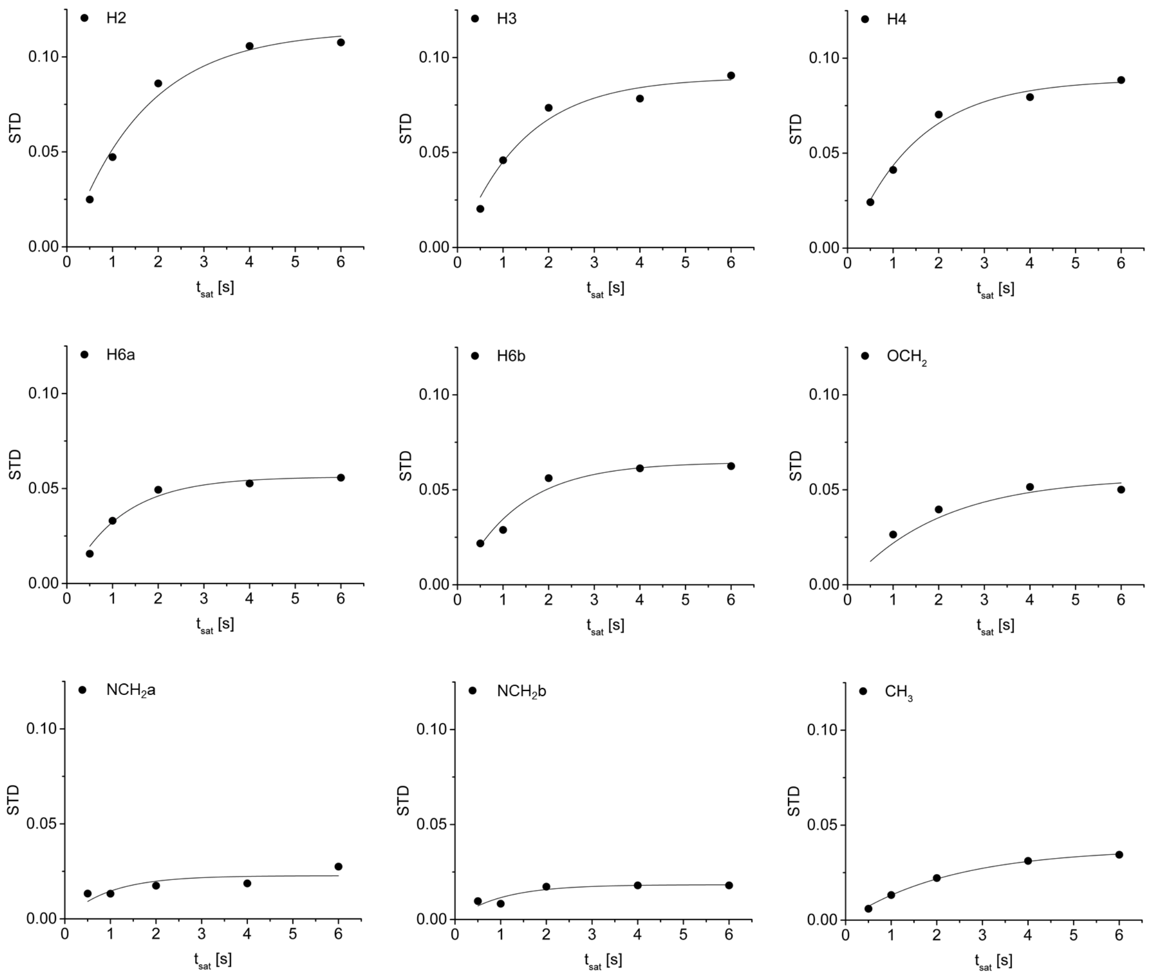


**Supplementary Figure 12. STD NMR build-up curves for Man analog 21.** Equation 5 was fitted to STD values to calculate $STD_0$' values for the determination of the binding epitope of **21**.

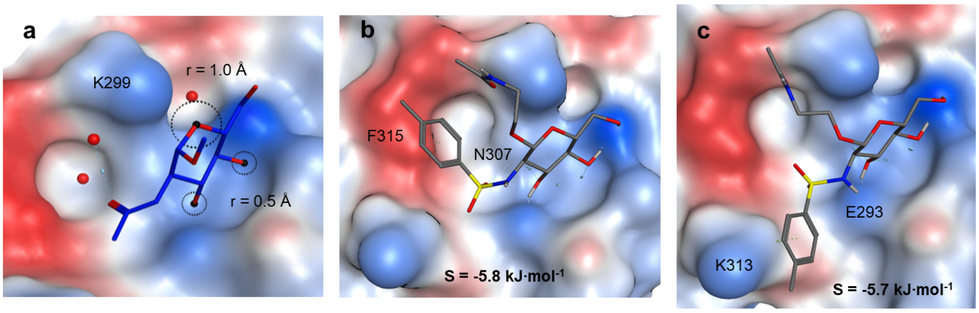


**Supplementary Figure 13. Molecular docking for GlcNS analog 16. a.** A pharmacophore model was defined to guide the initial placement of **16** in the carbohydrate binding site of Langerin (PDB code: 4N32) and to constrain the orientation of the Glc scaffold during the force field-based refinement of docking poses[9]. All features displayed require an oxygen atom within the indicated spheres. **b.** Four out of ten generated docking poses resemble the depicted conformation of **16**. The selected docking pose predicted the formation of π-π interactions between the phenyl ring and F315 as well as the formation of a hydrogen bond between the sulfonamide group and N307. The acetylated ethylamino linker displays high solvent exposure. Accordingly, this docking pose is consistent with both $^{15}$N HSQC and STD NMR experiments. **c.** The depicted alternative conformation of **16** is representative for three out of ten generated docking poses. The selected docking pose predicts the formation of cation-π interaction between the phenyl ring and K313 as well as the formation of a hydrogen bond between the sulfonamide and E293. The acetylated ethylamino linker displays high solvent exposure. However, this docking pose was less consistent with the $^{15}$N HSQC NMR results, particularly the relative decrease of CSP values for K313. The molecular docking study afforded three additional unique docking poses for **16** that were excluded due unfavorable dihedral angles for the sulfonamide linker. The receptor surface is colored according to its lipophilicity (lipophilic: red, hydrophilic: blue).

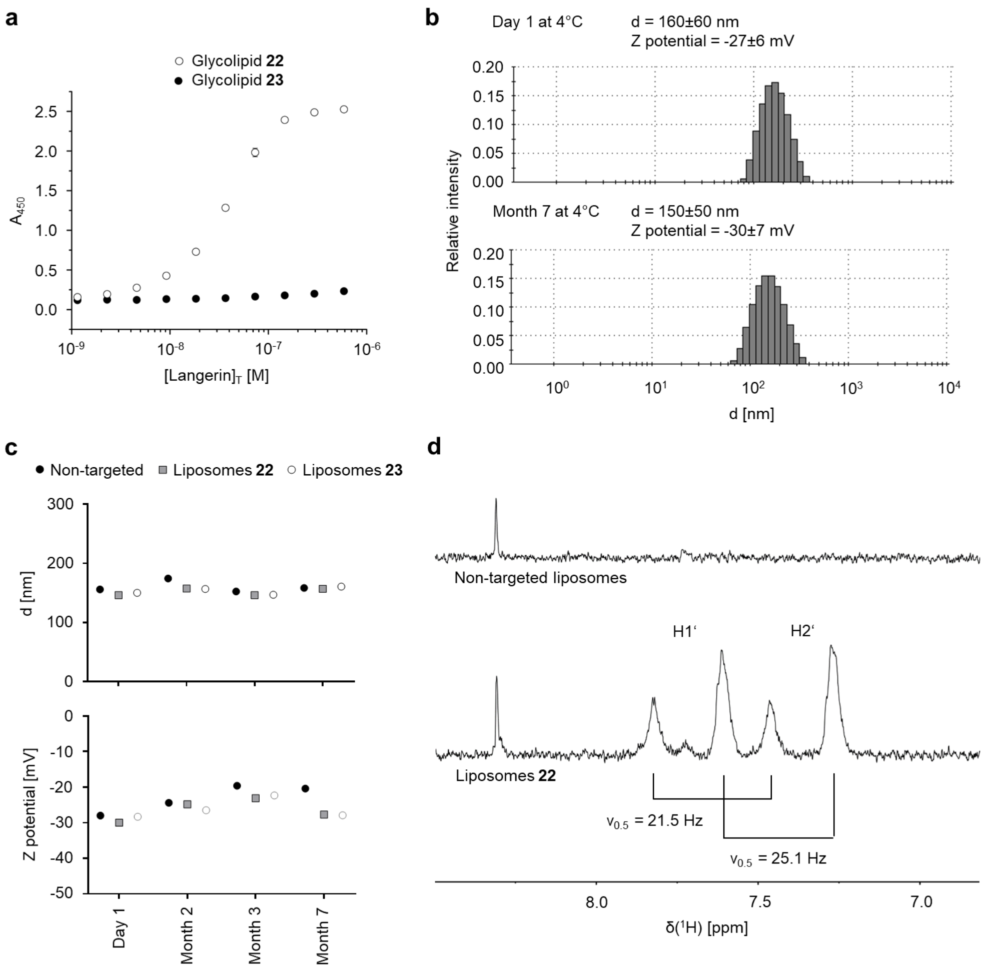


**Supplementary Figure 14. Characterization of liposomes 22 and 23. a.** Lipid-ELLA experiments served to evaluate the affinity of glycolipids **22** and **23** for the Langerin No binding was detected for glycolipid **23**, bearing Man analog **21**. In contrast, glycolipid **22**, bearing GlcNS analog **16** displayed a dose-dependent response validating the expected affinity increase over reference molecule **21**. **b. and c.** Dispersity and stability of the prepared liposomes were analyzed via DLS and electrophoresis experiments. Representative histograms are depicted for liposomes **22**. The liposomes are monodisperse (d = 160±60 nm) and stable (Z potential = -27±6 mV) for up to seven months when stored at 4°C in PBS. **d.** $^{1}$H NMR experiments with liposomes **22** revealed two states for **16** as observed for resonances corresponding to H1' and H2' of the phenyl ring. Both states display a linewidth $\nu_{0.5}$ smaller than 30 Hz, indicating accessibly and residual flexibility due to their presentation on extended PEG chains. The alternative state potentially corresponds to targeting ligands oriented towards the lumen of the liposomes.

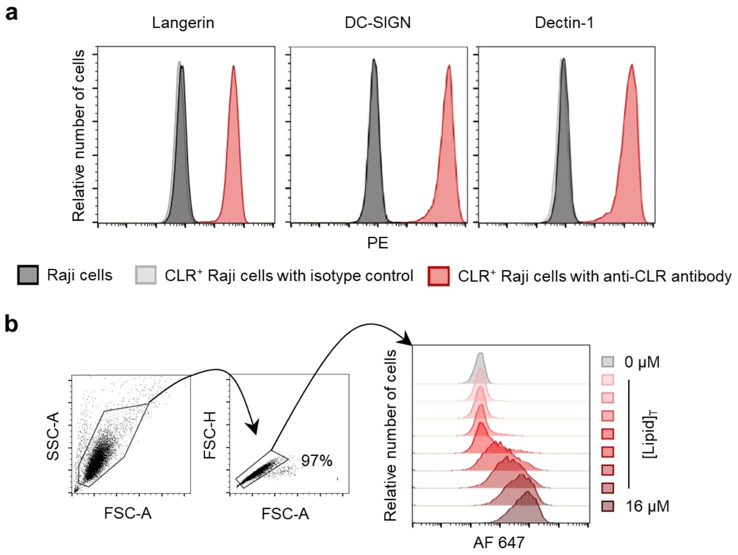

**Supplementary Figure 15.** ***In vitro*** **targeting of CLR⁺ Raji cells. a.** The expression of CLRs by the established Raji model cells was validated by flow cytometry using fluorophore-labeled anti-CLR antibodies. **b.** The gating strategy and the corresponding histograms for dose-dependent internalization and binding of liposomes to Langerin$^+$ Raji cells at 4°C are exemplarily shown.

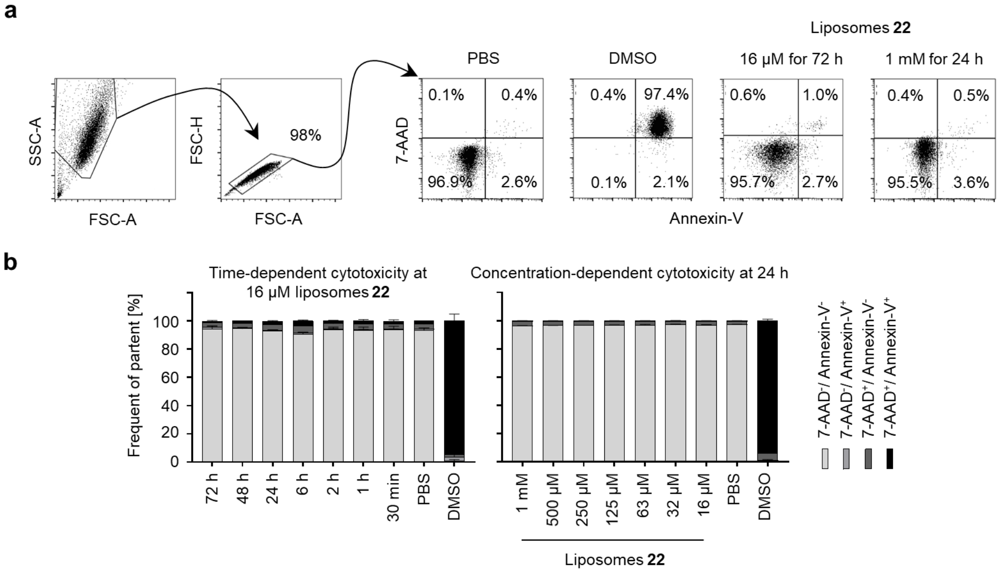


**Supplementary Figure 16. Induction of cytotoxicity by liposomes 22 with Langerin$^+$ Raji cells. a.** The gating strategy to evaluate the cytotoxicity of liposomes **22** is exemplarily shown for incubation of Langerin$^+$ Raji cells with liposomes **22** at total lipid concentrations $[Lipid]_T$ of 16 µM for 72 h and of 1 mM for 24 h. As a positive control, cells were exposed to 50% DMSO for 3 min following incubation for 72 h. **b.** Different incubation times and total lipid concentrations $[Lipid]_T$ were analyzed. Staining with the early and late apoptotic markers Annexin-V and 7-AAD demonstrated negligible cytotoxicity of liposomes **22** even at high concentrations and long-term exposure.

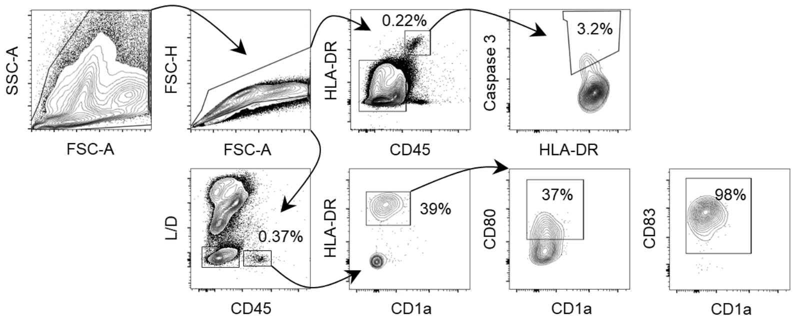


**Supplementary Figure 17. Induction of cytotoxicity and activation by liposomes 22 with LCs.** The gating strategy to evaluate the induction of cytotoxicity and activation by liposomes **22** with LCs is exemplarily shown after incubation at a total lipid concentration $[L]_T$ of 2.7 µM for 48 h. Using epidermal cell suspension, LCs were identified as HLA-DR$^+$-CD45$^+$-CD1a$^{high}$ cells with an additional viability gate to analyze their activation. Apoptosis was monitored by staining for active caspase 3. The activation of LCs was monitored via expression of CD80 and CD83.

### Supplementary Tables

**Supplementary Table 1. Structure-activity relationship of sulfated GlcNAc derivatives.**

| Name | Structure | $K_I$ [mM] | Relative potency[a] |
|---|---|---|---|
| Glc | 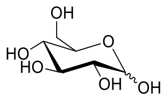 | 21±4 | 0.20 |
| GlcNAc | 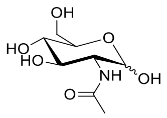 | 4.1±0.7 | 1.0 |
| GlcNS | 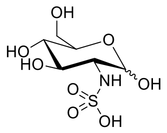 | 1.4±0.2 | 2.9 |
| GlcNAc-6-OS | 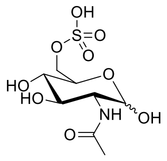 | 0.6±0.1 | 6.8 |
| GlcNS-6-OS | 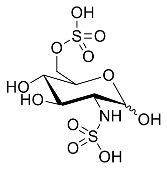 | 0.28±0.06 | 14 |

[a]The relative potency was calculated utilizing the $K_I$ value determined for GlcNAc.

**Supplementary Table 2. Structure-activity relationship of GlcNS analogs 1 to 5.**

| Name | Structure | R | $K_I$ [mM] | Relative potency[a] |
|---|---|---|---|---|
| **1** | 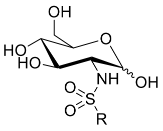 | 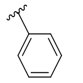 | 0.37±0.04 | 11 |
| **2** | | 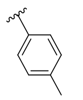 | 0.32±0.05 | 13 |
| **3** | | 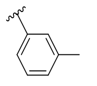 | 0.56±0.09 | 7.3 |
| **4** | | 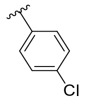 | 0.60±0.02 | 6.8 |
| **5** | | 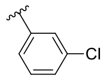 | 0.59±0.06 | 7.0 |

[a] The relative potency was calculated utilizing the $K_I$ value determined for GlcNAc.

**Supplementary Table 3. $^{19}$F $R_2$-filtered NMR assay development with DC-SIGN.**

| Parameter | CRD |
|---|---|
| $R_{2,f}$ [$s^{-1}$] | 1.8±0.3[a] |
| $R_{2,b}$ [$s^{-1}$] | 190±20[b] |
| $K_D$ [mM] for **24** | 2.3±0.5[b] |
| $K_I$ [mM] for Man | 3.0±0.3 |

[a] This value was determined from four independent experiments via $^{19}$F $R_2$-filtered NMR.
[b] This value was determined from three independent titration experiments via $^{19}$F $R_2$-filtered NMR.

### Supplementary Methods

#### Synthetic Chemistry – Procedures

**General remarks.** Reagents and solvents used were purchased from Sigma Aldrich unless indicated otherwise and used as supplied without any further purification. Anhydrous solvents were taken from an anhydrous solvent system (JC-Meyer Solvent Systems). Column chromatography was carried out using silica gel at a pore size from 40 Å to 60 Å (Machery Nagel). Reversed-phase column chromatography was carried out using Chromabond endcapped $C_{18}$ columns at a pore size of 60 Å (Machery Nagel or Merck Millipore). Analytical TLC was performed on glass plates coated with silica gel at a pore size of 60 Å (Machery Nagel). Compounds were detected via 3-methoxyphenol reagent (0.2% 3-methoxyphenol in EtOH: 2 N sulfuric acid in EtOH (1:1)), ninhydrin reagent (1.5 g ninhydrin in 15 ml acetic acid and 500 ml MeOH) or CAM reagent (1.0 g $Ce(SO_4)_2{\cdot}4H_2O$ and 2.5 g ammonium molybdate pentahydrate in 96 ml of $H_2O$ and 6 ml of concentrated $H_2SO_4$) upon heating or via UV adsorption ($\lambda$ = 254 nm). NMR experiments were either conducted on a PremiumCompact 400 MHz or 600 MHz spectrometer (Agilent) and an Avance II 500 MHz or an Avance III 400 MHz spectrometer (Bruker). Chemical shifts were referenced to the internal standards $CHCl_3$ ($\delta(^1H)$ = 7.26 ppm and $\delta(^{13}C)$ = 77.1 ppm), $H_2O$ ($\delta(^1H)$ = 7.26 ppm), MeOH ($\delta(^1H)$ = 4.87 ppm, $\delta(^{13}C)$ = 49.0 ppm). Coupling constants are reported in Hz and coupling patterns are indicated as s for singlet, d for doublets, dd for doublets of doublets, ddd for doublets of doublets of doublets, t for triplets, dt for doublets of triplets, td for triplet of doublets, q for quartets and m for multiplets. Signals were assigned by means of COSY, TOCSY, $^{13}C$ HSQC and H2BC NMR experiments[11]. Stereoselectivity at the anomeric position of the mannose and the glucosamine scaffold was analyzed by measuring $^1J_{C1,H1}$ coupling constants[12]. NMR spectra were processed with MestReNova[13]. ESI-MS analysis was conducted using an 1100 Series LC/MS coupled to a G1946D ESI-Q spectrometer (Agilent). HR ESI-MS analysis was conducted using an Acquity H-Class UPLC/MS coupled to a Xevo G2-S ESI-Q-TOF spectrometer (Waters). Preparative HPLC was performed on a 1100 Series LC/MS (Thermo Scientific) using a preparative Nuleodur C18 column (Machery Nagel). Analytical HPLC was performed on an Aqcuity UPLC system using an analytical BEH C18 column (Waters).

**2-deoxy-2-phenylsulfonylamido-D-glucopyranose**

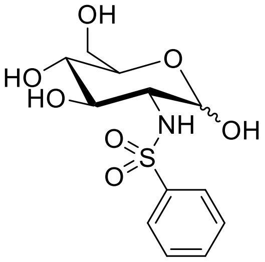

**1**

Glucosamine hydrochloride (50 mg, 232 µmol) was dissolved in anhydrous MeOH (2.3 ml) via the addition of $Et_3N$ (81 µl, 580 µmol). Benzenesulfonyl chloride (178 µl, 1.39 mmol) was added dropwise at 0°C under argon atmosphere. The reaction was allowed to heat up to room temperature and stirred for 2 h. More $Et_3N$ (32 µl, 230 µmol) and benzenesulfonyl chloride (89 µl, 695 µmol) were added dropwise at 0°C. The reaction was again allowed to heat up to room temperature and stirred for another 2h. Solvents were evaporated i*n vacuo* and the residue was purified via reversed-phase column chromatography (gradient: $H_2O$, 1%, 5% MeOH in $H_2O$ and elution with 10% MeOH in $H_2O$) and subsequently via column chromatography (gradient: DCM, 5% MeOH, 10% MeOH in DCM and elution with 20% MeOH in DCM). Silica particles were removed via filtration in MeOH with a cellulose acetate membrane at a pore size 0.2 µm to yield to yield an α/β-mixture of **1** (23.9 mg, 75 µmol, 32%) as a white solid.

The ratio of α- and β-anomer was determined to be 5:1 via $^1$H NMR. Here, only chemical shifts corresponding to the β-anomer are documented. As a result of the low signal intensities for the α-anomer the corresponding resonances were not assigned.

$^1$H NMR (400.0 MHz, MeOD, β-anomer): δ = 7.96 – 7.87 ppm, m, 2 H (aromatic H); δ = 7.64 – 7.45 ppm, m, 3 H (aromatic H); δ = 4.78 ppm, d, 1 H , J = 3.5 Hz (H1); δ = 3.76 – 3.68 ppm, m, 2 H (H6a, H5); δ = 3.69 – 3.55 ppm, m, 2 H (H6a, H3); δ = 3.30 – 3.24 ppm, m, 1 H (H4); δ = 3.13 ppm, dd, 1 H, J = 10.1, 3.5 Hz (H2).

$^{13}$C NMR (100.6 MHz, MeOD, β-anomer): δ = 143.4 ppm, 1 C (aromatic C); δ = 133.4 ppm, 1 C (aromatic C); δ = 130.0 ppm, 2 C (aromatic C); δ = 128.0 ppm, 2 C (aromatic C); δ = 92.9 ppm, 1 C (C1); δ = 72.8 ppm, 1 C (C5); δ = 72.5 ppm, 1 C (C3); δ = 72.2 ppm, 1 C (C4); δ = 62.6 ppm, 1 C (C6); δ = 59.9 ppm, 1 C (C2).

$R_f$ = 0.50 with 20% MeOH in DCM.

HR ESI-MS for $C_{12}H_{17}NO_7S$: $m \cdot z^{-1}(M+Na^+)_{calc}$ = 342.062; $m \cdot z^{-1}(M+Na^+)_{obs}$ = 342.063.

**2-deoxy-2-N-tosyl-D-glucopyranose**

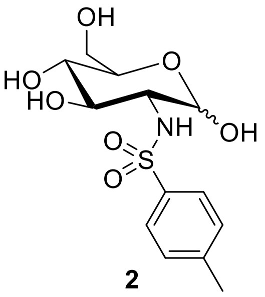

**2**

Glucosamine hydrochloride (3.0 g, 13.9 mmol) were dissolved in acetone (15 ml). Tosyl chloride (2.6 g, 13.9 mmol) was added and reaction was stirred for 4 h at room temperature. Solvents were removed *in vacuo* and the residue was purified via flash chromatorgraphy (15% MeOH in DCM) to yield an α/β-mixture of **2** (4.1 g, 12.5 mmol, 90%) as a white solid.

The ratio of α- and β-anomer was determined to be 10:1 via $^{1}$H NMR. Here, only chemical shifts corresponding to the β-anomer are documented. As a result of the low signal intensities for the α-anomer the corresponding resonances were not assigned.

$^{1}$H NMR (400.0 MHz, MeOD, β-anomer): δ = 7.82 – 7.77 ppm, m, 2 H (aromatic H); δ = 7.37 – 7.32 ppm, m, 2 H (aromatic H); δ = 4.77 ppm, d, 1 H , J = 3.5 Hz (H1); δ = 3.76 – 3.69 ppm, m, 2 H (H6a, H5); δ = 3.68 – 3.55 ppm, m, 2 H (H6a, H3); δ = 3.30 – 3.24 ppm, m, 1 H (H4); δ = 3.09 ppm, dd, 1 H, J = 10.1, 3.6 Hz (H2); δ = 2.41 ppm, s, 3 H ($CH_3$) .

$^{13}$C NMR (100.6 MHz, MeOD, β-anomer): δ = 144.4 ppm, 1 C (aromatic C); δ = 140.4 ppm, 1 C (aromatic C); δ = 130.6 ppm, 2 C (aromatic C); δ = 128.1 ppm, 2 C (aromatic C); δ = 92.9 ppm, 1 C (C1); δ = 72.8 ppm, 1 C (C5); δ = 72.5 ppm, 1 C (C3); δ = 72.2 ppm, 1 C (C4); δ = 62.6 ppm, 1 C (C6); δ = 59.8 ppm, 1 C (C2); δ = 21.5 ppm, 1 C ($CH_3$).

$R_f$ = 0.38 with 10% MeOH in DCM.

HR ESI-MS for $C_{13}H_{19}NO_7S$: $m{\cdot}z^{-1}(M+Na^{+})_{calc}$ = 356.078; $m{\cdot}z^{-1}(M+Na^{+})_{obs}$ = 356.077.

**2-deoxy-2-3'-(methyl)phenylsulfonylamido-D-glucopyranose**

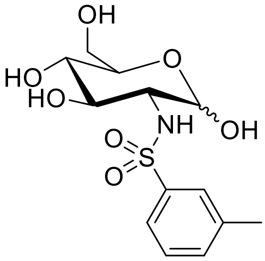


**3**

Glucosamine hydrochloride (49 mg, 227 µmol) was dissolved in DMSO (4.6 ml) via addition of DIPEA (240 µl, 1.39 mmol). Next, 3-methylphenylsulfonyl chloride (84 µl, 580 µmol) was slowly added and the reaction mixture was stirred for 2 h at room temperature. The reaction was quenched with MeOH and solvents were removed *in vacuo*. Residual DMSO was removed via lyophilization. The residue was purified via column chromatography (gradient: DCM, 5% MeOH in DCM and elution with 10% MeOH in DCM) and reversed-phase column chromatography (gradient: $H_2O$, 1%, 5% MeOH in $H_2O$ and elution with 10% MeOH in $H_2O$) to yield **3** (12.9 mg, 39 µmol, 17%) as a white solid after lyophilization.

The ratio of α- and β-anomer was determined to be 10:1 via $^1$H NMR. Here, only chemical shifts corresponding to the β-anomer are documented. As a result of the low signal intensities for the α-anomer the corresponding resonances were not assigned.

$^1$H NMR (400.0 MHz, MeOD, β-anomer): δ = 7.78 – 7.66 ppm, m, 2 H (aromatic H); δ = 7.45 – 7.38 ppm, m, 2 H (aromatic H); δ = 4.78 ppm, d, 1 H , J = 3.6 Hz (H1); δ = 3.76 – 3.68 ppm, m, 2 H (H6a, H5); δ = 3.68 – 3.56 ppm, m, 2 H (H6a, H3); δ = 3.30 – 3.24 ppm, m, 1 H (H4); δ = 3.12 ppm, dd, 1 H, J = 10.5, 3.6 Hz (H2); δ = 2.42 ppm, s, 3 H ($CH_3$) .

$^{13}$C NMR (100.6 MHz, MeOD, β-anomer): δ = 143.2 ppm, 1 C (aromatic C); δ = 140.4 ppm, 1 C (aromatic C); δ = 134.1 ppm, 1 C (aromatic C); δ = 129.9 ppm, 1 C (aromatic C); δ = 128.3 ppm, 1 C (aromatic C); δ = 125.1 ppm, 1 C (aromatic C); δ = 92.9 ppm, 1 C (C1); δ = 72.8 ppm, 1 C (C5); δ = 72.6 ppm, 1 C (C3); δ = 72.2 ppm, 1 C (C4); δ = 62.7 ppm, 1 C (C6); δ = 59.9 ppm, 1 C (C2); δ = 21.3 ppm, 1 C ($CH_3$).

$R_f$ = 0.34 with 10% MeOH in DCM.

HR ESI-MS for $C_{12}H_{19}NO_7S$: $m{\cdot}z^{-1}(M{+}Na^+)_{calc}$ = 356.078; $m{\cdot}z^{-1}(M{+}Na^+)_{obs}$ = 356.079.

**2-deoxy-2-4'-(chloro)phenylsulfonylamido-D-glucopyranose**

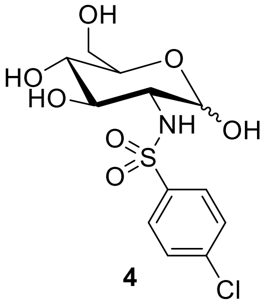

**4**

Glucosamine hydrochloride (51 mg, 234 mmol) was dissolved in anhydrous MeOH (2.3 ml) via the addition of DIPEA (240 µl, 1.39 mmol). 4-chlorophenylsulfonyl chloride (196 mg, 928 µmol) was dissolved in anhydrous MeOH (1.0 ml) and slowly added. The reaction mixture was stirred 4 h at room temperature. Solvents were removed *in vacuo* and the residue was purified via reversed-phase column chromatography (gradient: $H_2O$, 1% MeOH in $H_2O$ and elution with 5% MeOH in $H_2O$) to yield **4** (35 mg, 99 µmol, 42%) as a white solid after lyophilization.

The ratio of α- and β-anomer was determined to be 4:1 via $^1$H NMR. Here, only chemical shifts corresponding to the β-anomer are documented. As a result of the low signal intensities for the α-anomer the corresponding resonances were not assigned.

$^1$H NMR (400.0 MHz, MeOD, β-anomer): δ = 7.92 – 7.87 ppm, m, 2 H (aromatic H); δ = 7.56 – 7.51 ppm, m, 2 H (aromatic H); δ = 4.86 ppm, d, 1 H , J = 3.6 Hz (H1); δ = 3.77 – 3.69 ppm, m, 2 H (H6a, H5); δ = 3.68 – 3.55 ppm, m, 2 H (H6a, H3); δ = 3.29 – 3.23 ppm, m, 1 H (H4); δ = 3.14 ppm, dd, 1 H, J = 10.2, 3.5 Hz (H2).

$^{13}$C NMR (100.6 MHz, MeOD, β-anomer): δ = 142.3 ppm, 1 C (aromatic C); δ = 139.4 ppm, 1 C (aromatic C); δ = 130.1 ppm, 2 C (aromatic C); δ = 129.8 ppm, 2 C (aromatic C); δ = 93.2 ppm, 1 C (C1); δ = 72.8 ppm, 1 C (C5); δ = 72.5 ppm, 1 C (C3); δ = 72.3 ppm, 1 C (C4); δ = 62.6 ppm, 1 C (C6); δ = 59.9 ppm, 1 C (C2).

$R_f$ = 0.30 with 10% MeOH in DCM.

HR ESI-MS for $C_{12}H_{16}ClNO_7S$: $m \cdot z^{-1}(M+Na^+)_{calc}$ = 376.023; $m \cdot z^{-1}(M+Na^+)_{obs}$ = 376.024.

**2-deoxy-2-3'-(chloro)phenylsulfonylamido-D-glucopyranose**

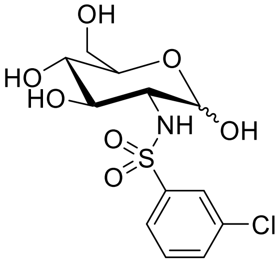


**5**

Glucosamine hydrochloride (50 mg, 234 µmol) was dissolved in anhydrous MeOH (2.3 ml) via addition of DIPEA (240 µl, 1.39 mmol). 3-chlorophenylsulfonyl chloride (130 µl, 930 µmol) was slowly added and the reaction mixture was stirred for 4 h at room temperature. Solvents were removed *in vacuo* and the residue was purified via reversed-phase column chromatography (gradient: $H_2O$, 1% MeOH in $H_2O$ and elution with 5% MeOH in $H_2O$) to yield **5** (6.0 mg, 17 µmol, 7%) as a white solid after lyophilization.

The ratio of α- and β-anomer was determined to be 8:1 via $^1$H NMR. Here, only chemical shifts corresponding to the β-anomer are documented. As a result of the low signal intensities for the α-anomer the corresponding resonances were not assigned.

$^1$H NMR (400.0 MHz, MeOD, β-anomer): δ = 7.97 – 7.79 ppm, m, 2 H (aromatic H); δ = 7.63 – 7.47 ppm, m, 2 H (aromatic H); δ = 4.87 ppm, m, 1 H (H1); δ = 3.78 – 3.69 ppm, m, 2 H (H6a, H5); δ = 3.69 – 3.55 ppm, m, 2 H (H6a, H3); δ = 3.30 – 3.24 ppm, m, 1 H (H4); δ = 3.12 ppm, dd, 1 H, J = 10.6, 3.5 Hz (H2).

$^{13}$C NMR (100.6 MHz, MeOD, β-anomer): δ = 145.5 ppm, 1 C (aromatic C); δ = 135.8 ppm, 1 C (aromatic C); δ = 133.2 ppm, 1 C (aromatic C); δ = 131.6 ppm, 1 C (aromatic C); δ = 128.0 ppm, 1 C (aromatic C); δ = 126.4 ppm, 1 C (aromatic C); δ = 93.2 ppm, 1 C (C1); δ = 72.8 ppm, 1 C (C5); δ = 72.5 ppm, 1 C (C3); δ = 72.3 ppm, 1 C (C4); δ = 62.6 ppm, 1 C (C6); δ = 60.0 ppm, 1 C (C2).

$R_f$ = 0.51 with 10% MeOH in DCM.

HR ESI-MS for $C_{12}H_{16}ClNO_7S$: $m \cdot z^{-1}(M+Na^+)_{calc}$ = 376.023; $m \cdot z^{-1}(M+Na^+)_{obs}$ = 376.024.

**(N-(Benzyloxycarbonyl)-2-aminoethyl)-2-deoxy-2-N-(2,2,2-trichlorethyloxy-carbonyl)-3,4,6-tri-O-acetyl-β-D-glucospyrannoside**

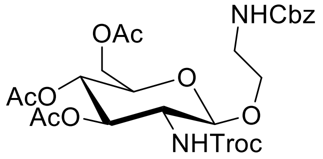

**11**

Intermediate **10** was prepared as previously published[3]. **10** (1.0 g, 1.0 mmol) and benzyl 2-hydroxyethylcarbamate (0.60 g, 3.1 mmol) were dissolved in DCM (40 ml) and stirred for 45 min at room temperature. DMTST (1.0 g, 3.8 mmol) was added and the reaction mixture was stirred overnight at room temperature. Solvents were removed *in vacuo* and the residue was purified via flash column chromatography (cyclohexane:EtOAc (1:1)) to yield **11** as a white solid (1.0 g, 1.4 mmol, 79%).

$^{1}$H NMR (300.0 MHz, $CDCl_3$): δ = 7.40 – 7.29 ppm, m, 5 H (aromatic H); δ = 5.35 – 5.16 ppm, m, 3 H (NH (NHCbz, NHTroc), H3); δ = 5.13 – 4.99 ppm, m, 3 H ($CH_2$ (NHCbz), H3); δ = 4.76 ppm, d, 1 H, J = 11.7 Hz (CHa (NHTroc)); δ = 4.58 ppm, d, 1 H, J = 8.3 Hz (H1); δ = 4.51 ppm, d, 1 H, J = 12.0 Hz (CHb (NHTroc)); δ = 4.24 ppm, dd, 1 H, J = 4.9, 12.3 Hz (H6a); δ = 4.13 ppm, dd, 1 H, J = 2.1, 12.1 Hz (H6b); ); δ = 3.86 ppm, dd, 1 H, J = 3.9, 10.1 Hz (OC*Ha*$CH_2$NHCbz); δ = 3.70 - 3.64 ppm, m, 3 H (OC*Hb*$CH_2$NHCbz, H2, H5); δ = 3.53 - 3.41 ppm, m, 1 H (O$CH_2$C*Ha*NHCbz); ); δ = 3.34 ppm, m, 1 H (O$CH_2$C*Hb*NHCbz); δ = 2.06 ppm, s, 3 H ($OCOCH_3$); δ = 2.03 ppm, s, 3 H ($OCOCH_3$); δ = 2.02 ppm, s, 3 H ($OCOCH_3$).

$^{13}$C NMR (125.7 MHz, $CDCl_3$): δ = 170.77 ppm, 1 C (O*C*O$CH_3$); δ = 170.72 ppm, 1 C (O*C*O$CH_3$); δ = 169.46 ppm, 1 C (O*C*O$CH_3$); δ = 156.55 ppm, 1 C (N*C*OO (NHTroc)); δ = 154.29 ppm, 1 C (N*C*OO (NHCbz)); δ = 136.47 ppm, 1 C (aromatic C); δ = 128.59 ppm, 1 C (aromatic C); δ = 128.24 ppm, 1 C (aromatic C); δ = 128.22 ppm, 1 C (aromatic C); δ = 101.06 ppm, 1 C (C1); δ = 95.43 ppm, 1 C (*C*$Cl_3$); δ = 74.42 ppm, 1 C (*C*$H_2$ (NHTroc)); δ = 71.92 ppm, 2 C (C3, C5); δ = 69.33 ppm, 1 C (O*C*$H_2CH_2$NHCbz); δ = 68.45 ppm, 1 C (C4); δ = 66.84 ppm, 1 C ($CH_2$ (NHCbz)); δ = 61.96 ppm, 1 C (C6); δ = 56.23 ppm, 1 C (C2); δ = 40.73 ppm, 1 C (O$CH_2$*C*$H_2$NHCbz); 1 C (C2); δ = 20.74 ppm, 1 C (OCO*C*$H_3$); δ = 20.64 ppm, 2 C (OCO*C*$H_3$).

$R_f$ = 0.35 with cyclohexane:EtOAc (1:1).

ESI-MS for $C_{25}H_{31}Cl_3N_2O_{12}$: $m \cdot z^{-1}(M+Na^+)_{calc}$ = 679.1; $m \cdot z^{-1}(M+Na^+)_{obs}$ = 671.0.

**(N-(Benzyloxycarbonyl)-2-aminoethyl)-2-deoxy-2-N-tosyl-3,4,6-tri-O-acetyl-β-D-glucospyrannoside**

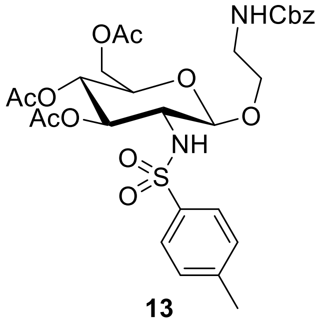

**13**

**11** (1.1 g, 2.0 mmol) was dissolved in acetic acid (60 ml). Activated zinc (20 g, 260 mmol) was added and the reaction mixture was stirred for 4 h at room temperature. Subsequently, the reaction mixture was filtered over Celite and solvents were removed *in vacuo*. The residue was dissolved in pyridine (4 ml) and tosyl chloride (110 mg, 0.60 mmol) was added slowly under argon atmosphere. The reaction mixture was stirred overnight at room temperature. Solvents were removed *in vacuo* and the residue was purified via flash chromatography (cyclohexane:EtOAc (1:1)) to yield **13** (640 mg, 1.0 mmol, 50%) as a white solid.

$^1$H NMR (500.0 MHz, $CDCl_3$): δ = 7.64 ppm, d, 2 H, J = 8.2 Hz (aromatic H (NHTs)); δ = 7.32 – 7.22 ppm, m, 5 H (aromatic H (NHCbz)); δ = 7.16 ppm, d, 2 H, J = 8.1 Hz (aromatic H (NHTs)); δ = 5.41 ppm, d, 1 H, J = 8.3 Hz (NH (NHTs)); δ = 5.25 ppm, m, 1 H (NH (NHCbz)); δ = 5.03 ppm, s, 2 H ($CH_2$ (NHCbz)); δ = 4.95 ppm, m, 2 H (H3, H4); δ = 4.27 ppm, d, 1 H, J = 8.2 Hz (H1); δ = 4.14 ppm, dd, 1 H, J = 4.8, 12.3 Hz (H6a); δ = 4.01 ppm, dd, 1 H, J = 2.2, 12.3 Hz (H6b); δ = 3.68 – 3.60 ppm, m, 1 H (OC*Ha*$CH_2$NHCbz); δ = 3.57 – 3.50 ppm, m, 1 H (H5); δ = 3.46 – 3.37 ppm, m, 1 H (H2); δ = 3.37 – 3.31 ppm, m, 1 H (OC*Hb*$CH_2$NHCbz); δ = 3.21 – 3.11 ppm, m, 1 H (O$CH_2$C*Ha*NHCbz); δ = 3.11 – 3.00 ppm, m, 1 H (O$CH_2$C*Hb*NHCbz); δ = 2.29 ppm, s, 3 H ($CH_3$ (NHTs)); δ = 1.98 ppm, s, 3 H (OCO$CH_3$); δ = 1.94 ppm, s, 3 H (OCO$CH_3$); δ = 1.81 ppm, s, 3 H (OCO$CH_3$).

$^{13}$C NMR (125.7 MHz, $CDCl_3$): δ = 171.41 ppm, 1 C (O*C*O$CH_3$); δ = 170.82 ppm, 1 C (O*C*O$CH_3$); δ = 169.45 ppm, 1 C (O*C*O$CH_3$); δ = 156.77 ppm, 1 C (N*C*OO (NHCbz)); δ = 143.51 ppm, 1 C (aromatic C (NHTs)); δ = 138.49 ppm, 1 C (aromatic C (NHTs)); δ = 136.72 ppm, 1 C (aromatic C (NHCbz)); δ = 129.55 ppm, 2 C (aromatic C (NHTs)); δ = 128.62 ppm, 3 C (aromatic C (NHCbz)); δ = 128.21 ppm, 2 C (aromatic C (NHCbz)); δ = 127.27 ppm, 2 C (aromatic C (NHTs)); δ = 101.50 ppm, 1 C (C1); δ = 72.95 ppm, 1 C (C3); δ = 71.85 ppm, 1 C (C5); δ = 69.51 ppm, 1 C (O*C*$H_2$$CH_2$NHCbz); δ = 68.40 ppm, 1 C (C4); δ = 66.79 ppm, 1 C ($CH_2$ (NHCbz)); δ = 61.96 ppm, 1 C (6); δ = 58.08 ppm, 1 C (C2); δ = 40.58 ppm, 1 C (O$CH_2$*C*$H_2$NHCbz); δ = 26.11 ppm, 1 C ($CH_3$ (NHTs)); δ = 20.82 ppm, 1 C (OCO*C*$H_3$); δ = 20.72 ppm, 1 C (OCO*C*$H_3$); δ = 20.70 ppm, 1 C (OCO*C*$H_3$).

$R_f = 0.27$ with 10% MeOH in DCM.

ESI-MS for $C_{29}H_{36}N_2O_{12}S$: $m\cdot z^{-1}(M+H^+)_{calc} = 637.2$; $m\cdot z^{-1}(M+H^+)_{obs} = 637.2$.

**Aminoethyl-2-deoxy-2-N-tosyl-β-D-glucospyrannoside**

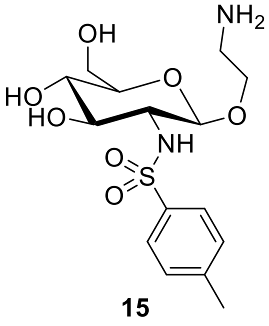


**15**

**13** (100 mg, 0.16 mmol) was dissolved in MeOH (5 ml) and 0.4 M sodium methanolate in MeOH (43 mg, 0.80 mmol) was added. The reaction mixture was stirred for 2 h at room temperature and solvents were removed *in vacuo*. The residue was dissolved in degassed ethanol and a catalytic amount of Pd/C (10%) was added. The reaction mixture was saturated with hydrogen at atmospheric pressure and stirred overnight at room temperature under hydrogen atmosphere. The reaction mixture was filtered over Celite, washed with EtOAc and solvents were removed *in vacuo*. The residue was purified by reversed-phase HPLC (gradient: 3% to 30% acetonitrile in 0.1% TFA in $H_2O$ in 30 min) to yield **15** (10 mg, 26 μmol, 16%) as a white solid after lyophilization.

$^1$H NMR (500.0 MHz, $D_2O$): δ = 7.75 ppm, d, 2 H, J = 8.4 Hz (aromatic H (NHTs)); δ = 7.39 ppm, d, 2 H, J = 8.5 Hz (aromatic H (NHTs)); δ = 4.41 ppm, d, 1 H, J = 8.4 Hz (H1); δ = 3.86 ppm, dd, 1 H, J = 2.0, 12.3 Hz (H6a); δ = 3.81 ppm, ddd, 1 H, J = 3.2, 8.2, 11.5 Hz (OC*Ha*$CH_2NH_2$); δ = 3.68 ppm, dd, 1 H, J = 5.4, 12.3 Hz (H6b); δ = 3.47 ppm, ddd, 1 H, J = 3.5, 5.6, 11.6 Hz (OC*Hb*$CH_2NH_2$); δ = 3.42 – 3.31 ppm, m, 3 H (H3, H4, H5); δ = 3.16 ppm, dd, 1 H, J = 8.4, 10.0 Hz (H2); δ = 3.02 ppm, ddd, 1 H, J = 3.2, 5.6, 13.6 Hz ($OCH_2C$*Ha*$NH_2$); δ = 2.71 ppm, ddd, 1 H, J = 3.5, 8.1, 13.5 Hz ($OCH_2C$*Hb*$NH_2$); δ = 2.40 ppm, s, 3 H ($CH_3$ (NHTs)).

$^{13}$C NMR (125.7 MHz, $D_2O$): δ = 144.48 ppm, 1 C (aromatic C (NHTs)); δ = 137.61 ppm, 1 C (aromatic C (NHTs)); δ = 129.59 ppm, 2 C (aromatic C (NHTs)); δ = 126.49 ppm, 2 C (aromatic C (NHTs)); δ = 101.17 ppm, 1 C (C1); δ = 75.58 ppm, 1 C (C4); δ = 73.91 ppm, 1 C (C3); δ = 69.71 ppm, 1 C (C5); δ = 65.65 ppm, 1 C (O*C*$H_2CH_2NH_2$); δ = 60.48 ppm, 1 C (C6); δ = 59.41 ppm, 1 C (C2); δ = 39.06 ppm, 1 C (O*C*$H_2CH_2NH_2$); δ = 20.54 ppm, 1 C ($CH_3$ (NHTs)).

HR ESI-MS for $C_{15}H_{24}N_2O_7S$: $m\cdot z^{-1}(M+H^+)_{calc} = 377.138$; $m\cdot z^{-1}(M+H^+)_{obs} = 377.152$.

**2-Acetamidoethyl-2-deoxy-2-N-tosyl-β-D-glucospyrannoside**

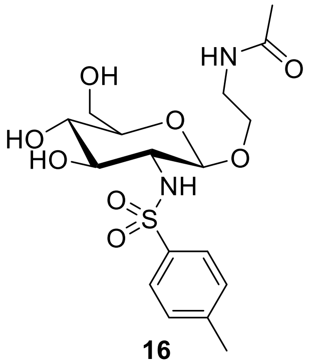


**16**

**15** (9.8 mg, 26 µmol) was dissolved in MeOH (0.7 ml) and acetic anhydride (50 µl, 490 µmol) was added slowly. The reaction mixture was stirred for 72 h at room temperature, quenched with diethlyl amine and solvents were removed *in vacuo*. The residue was purified by reversed-phase HPLC (gradient: 3% to 40% acetonitrile in 0.1% TFA in $H_2O$ in 30 min) to yield **16** (6 mg, 14 µmol, 54%) as a white solid after lyophilization.

$^1$H NMR (400.1 MHz, $D_2O$): δ = 7.69 ppm, d, 2 H, J = 8.2 Hz (aromatic H, in proximity of $NHSO_2$ (NHTs)); δ = 7.35 ppm, d, 2 H, J = 8.4 Hz (aromatic H, in proximity of $CH_3$ (NHTs)); δ = 4.33 ppm, d, 1 H, J = 8.2 Hz (H1); δ = 3.82 ppm, dd, 1 H, J = 1.5, 12.4 Hz (H6a); δ = 3.64 ppm, m, 1 H (H6b); δ = 3.60 ppm, m, 1 H (OC*Ha*$CH_2NHCOCH_3$); δ = 3.38 ppm, m, 1 H (H3); δ = 3.35 – 3.29 ppm, m, 2 H (H4, H5); δ = 3.23 ppm, m, 1 H (OC*Hb*$CH_2NHCOCH_3$); δ = 3.08 ppm, dd, 1 H, J = 8.4, 10.1 Hz (H2); δ = 2.99 ppm, ddd, 1 H, J = 4.1, 5.8, 14.5 Hz ($OCH_2C$*Ha*$NHCOCH_3$); δ = 2.80 ppm, ddd, 1 H, J = 3.9, 7.1, 14.3 Hz ($OCH_2C$*Hb*$NHCOCH_3$); δ = 2.35 ppm, s, 3 H ($CH_3$ (NHTs)): δ = 1.91 ppm, s, 3 H ($OCH_2CH_2NHCOC$*$H_3$*).

$^{13}$C NMR (100.6 MHz, $D_2O$): δ = 173.84 ppm, 1 C ($OCH_2CH_2NH$*C*$OCH_3$); δ = 144.28 ppm, 1 C (aromatic C (NHTs)); δ = 137.64 ppm, 1 C (aromatic C (NHTs)); δ = 129.50 ppm, 2 C (aromatic C (NHTs)); δ = 126.37 ppm, 1 C (aromatic C (NHTs)); δ = 101.06 ppm, 1 C (C1); δ = 75.56 ppm, 1 C (C5); δ = 74.17 ppm, 1 C (C3); δ = 69.71 ppm, 1 C (C4); δ = 67.87 ppm, 1 C (O*C*$H_2CH_2NHCOCH_3$); δ = 60.51 ppm, 1 C (C6); δ = 59.55 ppm, 1 C (C2); δ = 39.04 ppm, 1 C ($OCH_2$*C*$H_2NHCOCH_3$); δ = 21.64 ppm, 1 C ($OCH_2CH_2NHCO$*C*$H_3$); δ = 20.53 ppm, 1 C ($CH_3$ (NHTs)).

HR ESI-MS for $C_{15}H_{24}N_2O_7S$: $m \cdot z^{-1}(M+Na^+)_{calc}$ = 441.131; $m \cdot z^{-1}(M+Na^+)_{obs}$ = 441.131.

**2-Acetamidoethyl-α-D-mannopyrannoside**

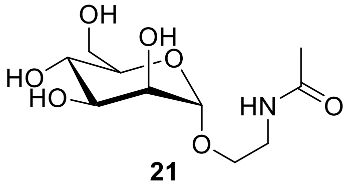


**21**

Intermediate **20** was prepared as previously published[4]. **20** (23 mg, 103 µmol) was dissolved in MeOH (700 µl) and acetic anhydride (10 µl, 110 µmol) was added slowly. The reaction mixture was stirred for 24 h at room temperature, quenched with 25 % (w/v) sodium methoxide in methanol and solvents were removed *in vacuo*. The residue was purified by reversed-phase HPLC (gradient: 0% to 10% acetonitrile in 0.1% TFA in $H_2O$ in 40 min) to yield **21** (17 mg, 26 µmol, 25%) as a white solid after lyophilization.

$^1$H NMR (400.1 MHz, $D_2O$): δ = 4.80 ppm, d, 1 H, J = 1.8 Hz (H1); δ = 3.88 ppm, dd, 1 H, J = 1.8, 3.3 Hz (H2); δ = 3.82 ppm, dd, 1 H, J = 2.2, 12.3 Hz (H6a); δ = 3.74 ppm, m, 1 H (H3); δ = 3.71 ppm, m, 1 H (OC*Ha*$CH_2$NHCO$CH_3$); δ = 3.67 ppm, m, 1 H (H6b); δ = 3.59 ppm, m, 1 H (H4); δ = 3.55 ppm, m, 1 H (H5); δ = 3.52 ppm, m, 1 H (OC*Hb*$CH_2$NHCO$CH_3$); δ = 3.52 ppm, ddd, 1 H, J = 3.7, 7.2, 14.4 Hz (O$CH_2$C*Ha*NHCO$CH_3$); δ = 3.31 ppm, ddd, 1 H, J = 3.9, 6.3, 14.4 Hz (O$CH_2$C*Hb*NHCO$CH_3$); δ = 1.95 ppm, s, 3 H (O$CH_2$$CH_2$NHCOC$H_3$).

$^{13}$C NMR (100.6 MHz, $D_2O$): δ = 174.17 ppm, 1 C (O$CH_2$$CH_2$NH*C*O$CH_3$); δ = 99.51 ppm, 1 C (C1); δ = 72.69 ppm, 1 C (C5); δ = 70.37 ppm, 1 C (C3); δ = 69.88 ppm, 1 C (C2); δ = 66.55 ppm, 1 C (C4); δ = 65.67 ppm, 1 C (O*C*$H_2$$CH_2$NHCO$CH_3$); δ = 60.77 ppm, 1 C (C6); δ = 38.90 ppm, 1 C (O$CH_2$*C*$H_2$NHCO$CH_3$); δ = 21.72 ppm, 1 C (O$CH_2$$CH_2$NHCO*C*$H_3$).

HR ESI-MS for $C_{15}H_{24}N_2O_7S$: $m \cdot z^{-1}(M\text{-}H^+)_{calc}$ = 264.108; $m \cdot z^{-1}(M\text{-}H^+)_{obs}$ = 264.109.

**16-PEG-DSPE**

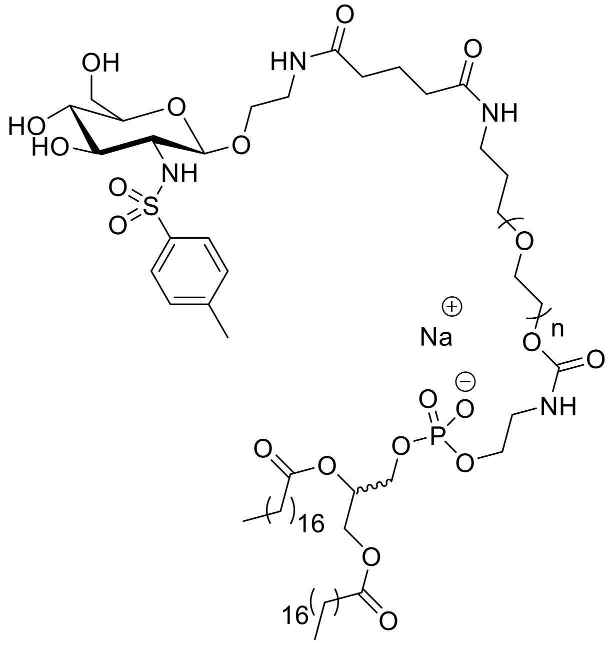

**22**

NHS-activated PEG-DSPE (3.1 kDa, NOF Europe) (2.8 mg, 900 nmol) was dissolved in DMF (140 µl) and slowly added to **16** (2.7 mg, 7.20 µmol) dissolved in 0.1 M $NaCO_3H$ at pH 8.4 (1.2 ml). The reaction mixture was stirred overnight at room temperature. Solvents were removed *in vacuo* and the residue was dissolved in 0.1 M $NaCO_3H$ at pH 8.4 (2.0 ml). Next, the reaction mixture was dialyzed three times against 0.1 M $NaCO_3H$ at pH 8.4 (5.0 ml) and three times against $H_2O$ (5.0 ml). After lyophilization, **22** (2.4 mg, 710 nmol, 79%) was obtained as a white solid. Product formation was analyzed via $^1H$ NMR and the yield was calculated via the integration of characteristic resonances.

$^1H$ NMR (400.0 MHz, DMSO-$d_6$, characteristic resonances): δ = 7.30 – 7.10 ppm, m, 2 H (aromatic H, **22**); δ = 0.85 ppm, t, 3 H (two times $CH_2CH_3$).

**21-PEG-DSPE**

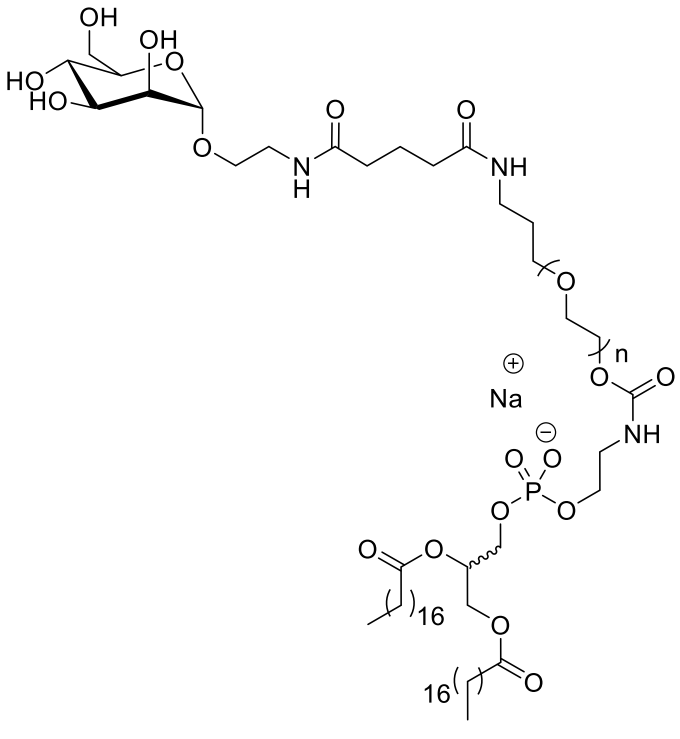

**23**

NHS-activated PEG-DSPE (3.1 kDa, NOF Europe) (2.5 mg, 810 nmol) was dissolved in DMF (130 µl) and slowly added to **21** (1.5 mg, 6.70 µmol) dissolved in 0.1 M $NaCO_3H$ at pH 8.4 (1.1 ml). The reaction mixture was stirred overnight at room temperature. Solvents were removed *in vacuo* and the residue was dissolved in 0.1 M $NaCO_3H$ at pH 8.4 (2.0 ml). Next, the reaction mixture was dialyzed three times against 0.1 M $NaCO_3H$ at pH 8.4 (5.0 ml) and three times against $H_2O$ (5.0 ml). After lyophilization, **23** (2.1 mg, 530 nmol, 66%) was obtained as a white solid. Product formation was analyzed via $^1H$ NMR and the yield was calculated via the integration of characteristic resonances.

$^1H$ NMR (400.0 MHz, DMSO-$d_6$, characteristic resonances): δ = 4.61 – 4.58 ppm, m, 1 H (H1, **12.29**): δ = 0.85 ppm, t, 3 H (two times $CH_2CH_3$).

**Alexa Fluor 647-PEG-DSPE**

Amino-PEG-DSPE (3.1 kDa, NOF Europe) (1.0 mg, 320 nmol) was dissolved in DMSO (500 µl) and slowly added to NHS-activated Alexa Fluor 647 (Thermo Scientific) (1.0 mg, 1.6 nmol) dissolved in DMSO (500 µl). The reaction mixture was stirred overnight at room temperature and protected from light. Solvents were removed by lyophilization and the residue was dissolved in 0.1 M $NaCO_3H$ at pH 8.4 (2.0 ml). Next, the reaction mixture was dialyzed three times against 0.1 M $NaCO_3H$ at pH 8.4 (5.0 ml) and three times against $H_2O$ (5.0 ml). After lyophilization, Alexa Fluor 647-PEG-DSPE (0.9 mg, 240 nmol) was obtained as a red solid.

#### Synthetic Chemistry – $^{1}$H and $^{13}$C NMR Spectra

##### 2-deoxy-2-phenylsulfonylamido-D-glucopyranose (1)

$^{1}$H NMR (400.0 MHz, MeOD)

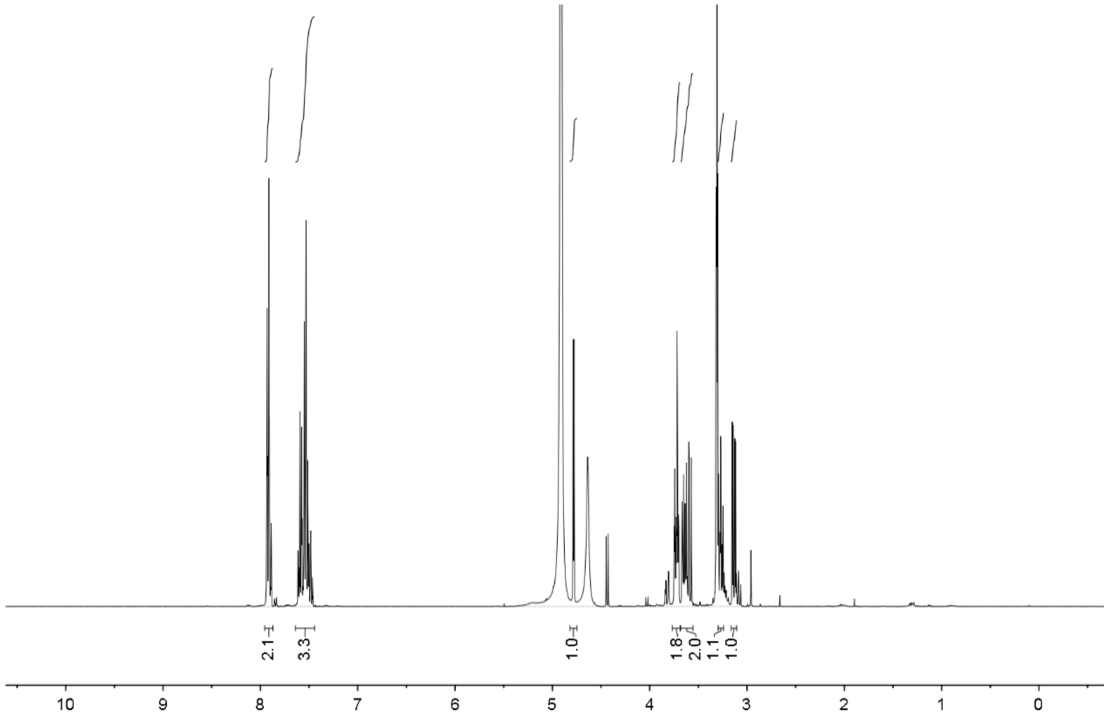


$^{13}$C NMR (100.6 MHz, MeOD)

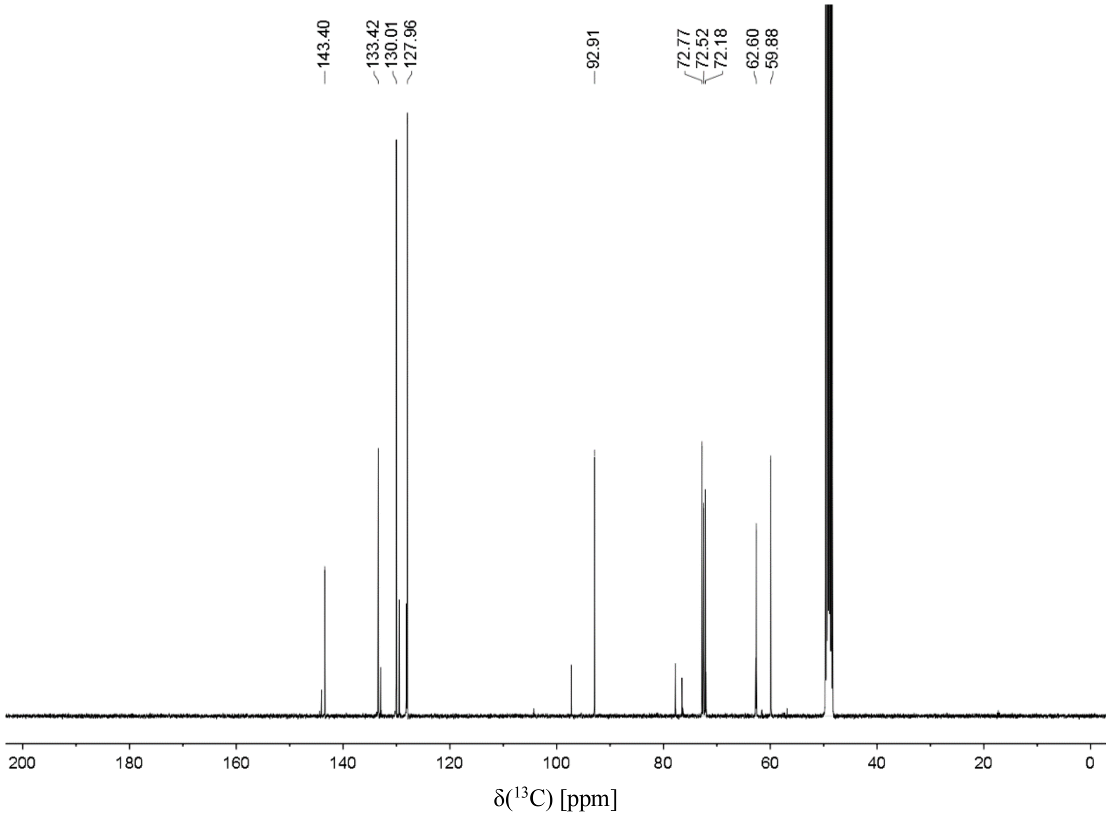

**2-deoxy-2-N-tosyl-D-glucopyranose (2)**

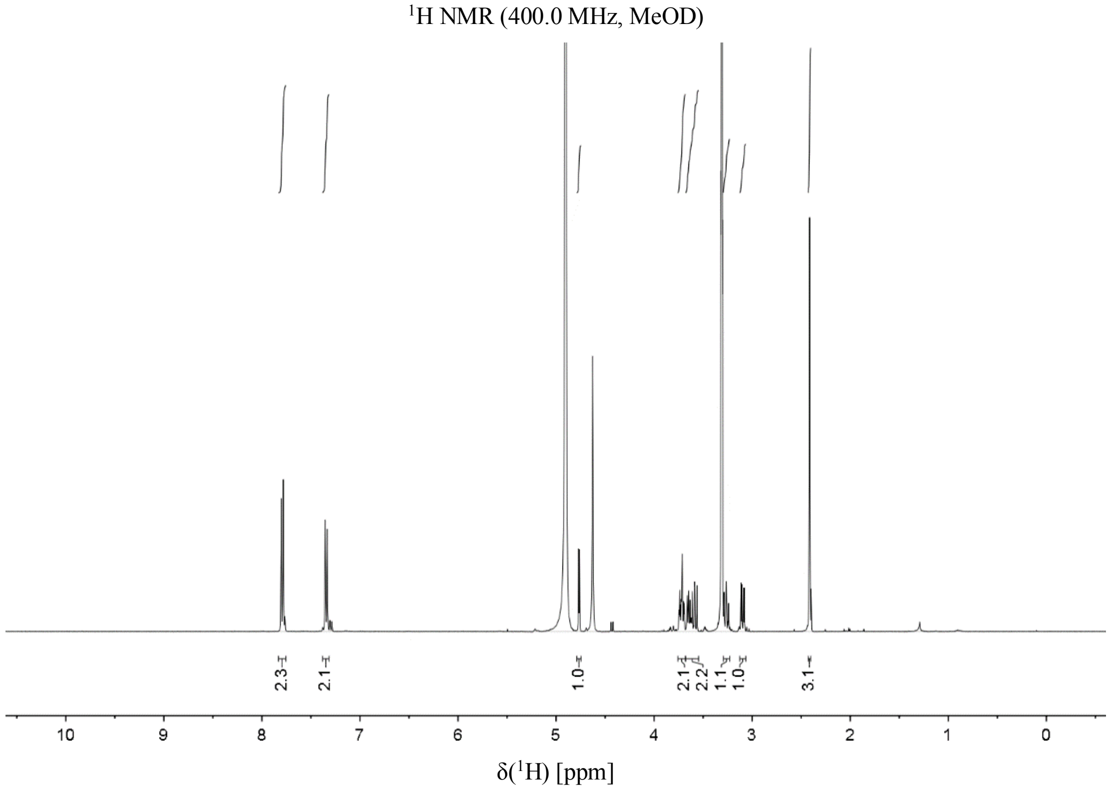


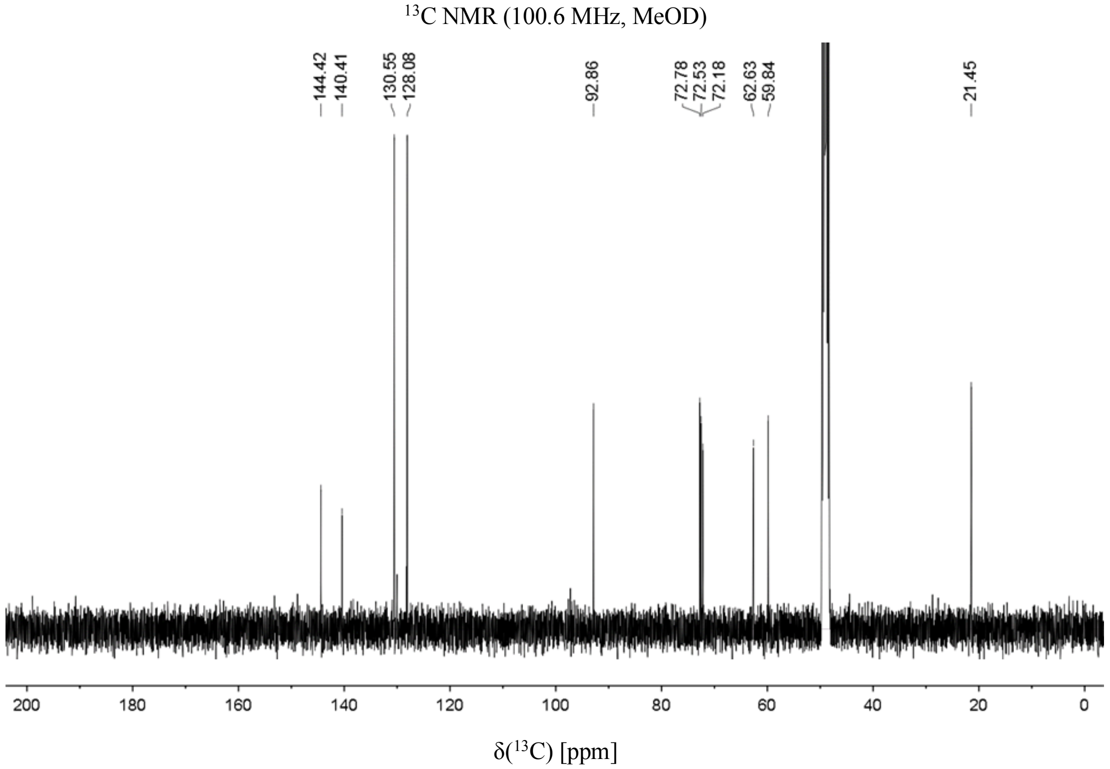

**2-deoxy-2-3'-(methyl)phenylsulfonylamido-D-glucopyranose (3)**

$^{1}H$ NMR (400.0 MHz, MeOD)

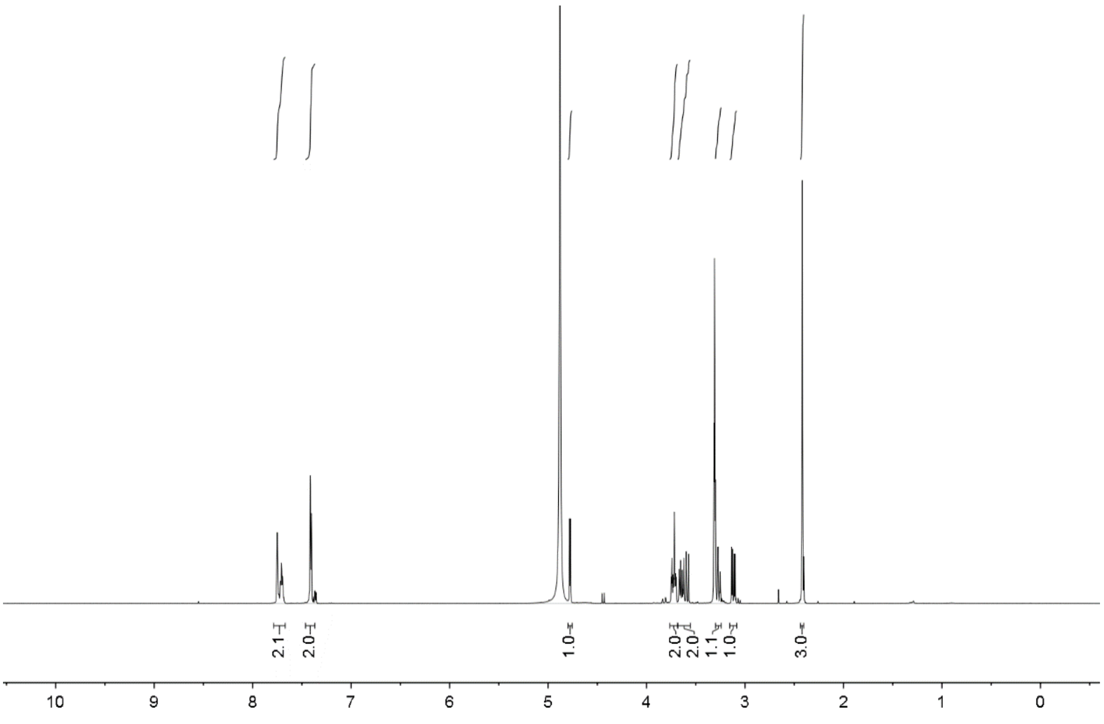


$^{13}C$ NMR (100.6 MHz, MeOD)

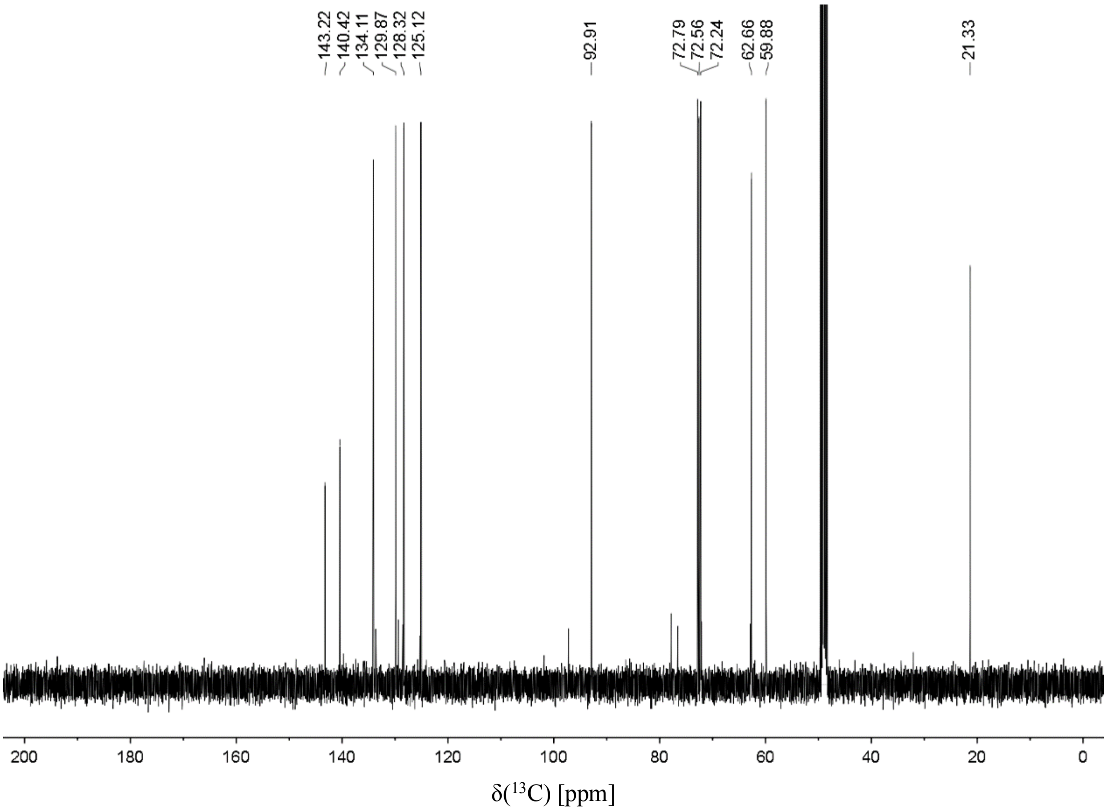

**2-deoxy-2-4’-(chloro)phenylsulfonylamido-D-glucopyranose (4)**

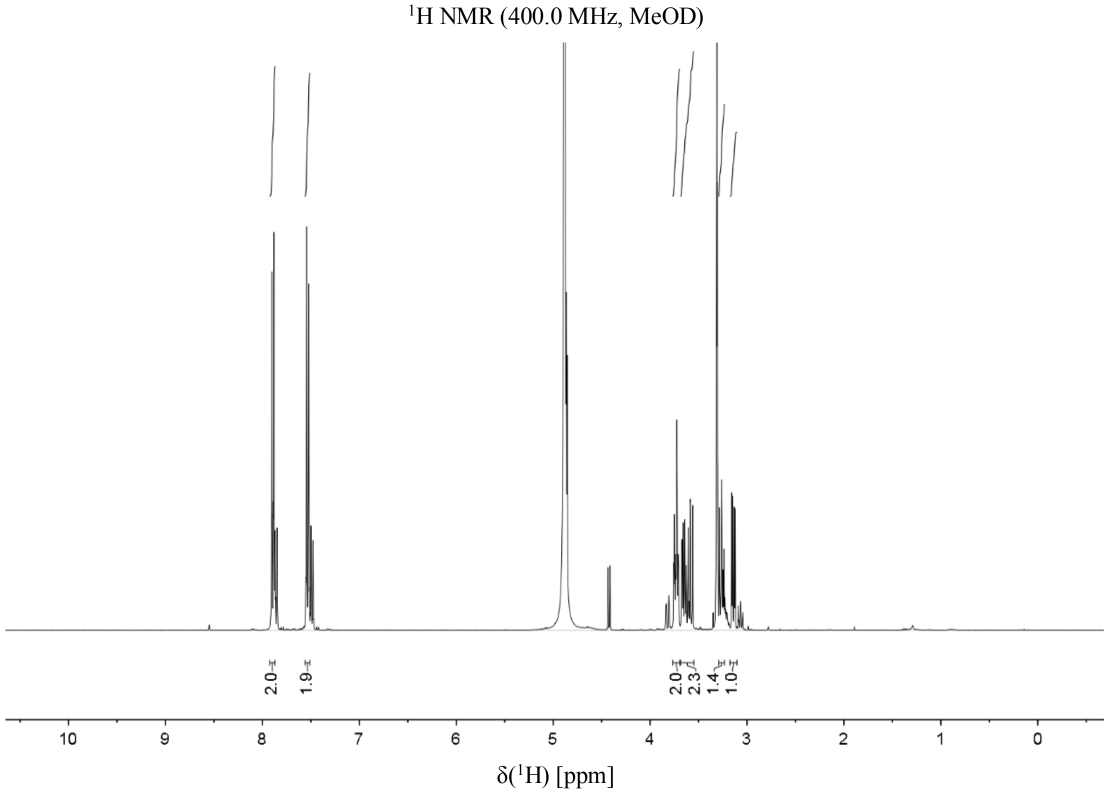


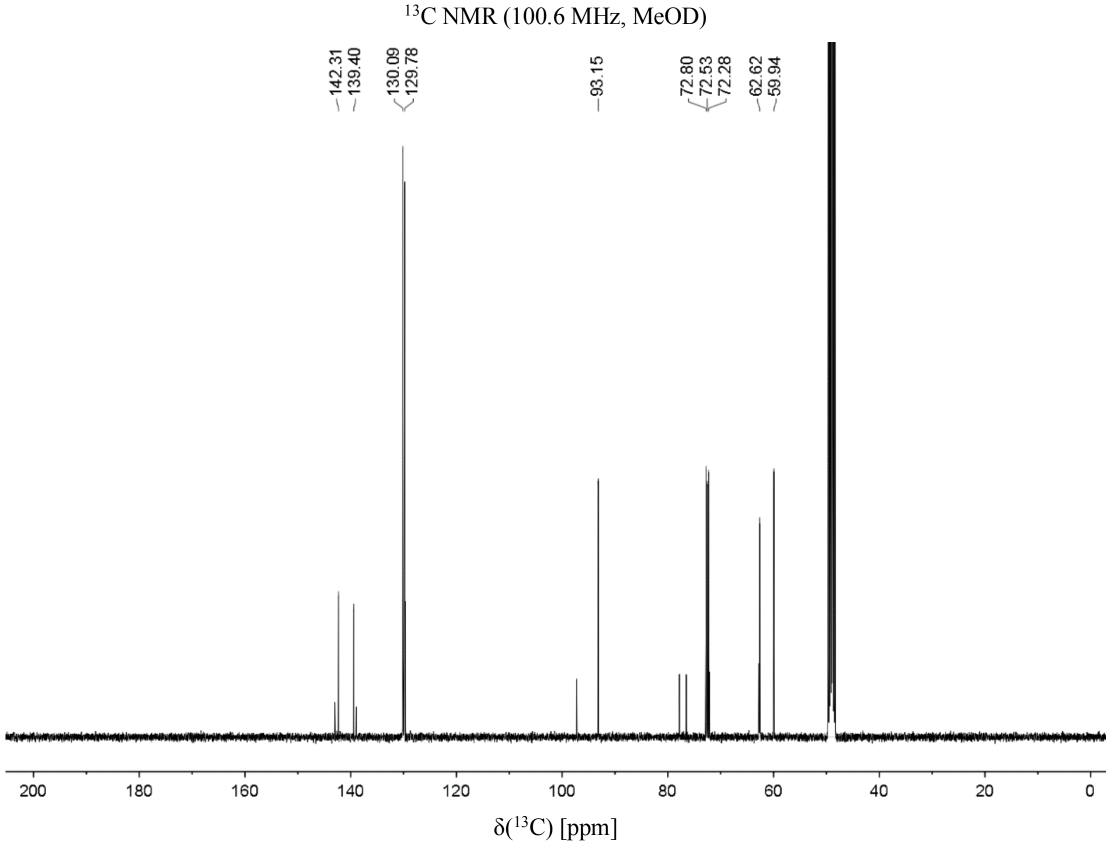

**2-deoxy-2-3'-(chloro)phenylsulfonylamido-D-glucopyranose (5)**

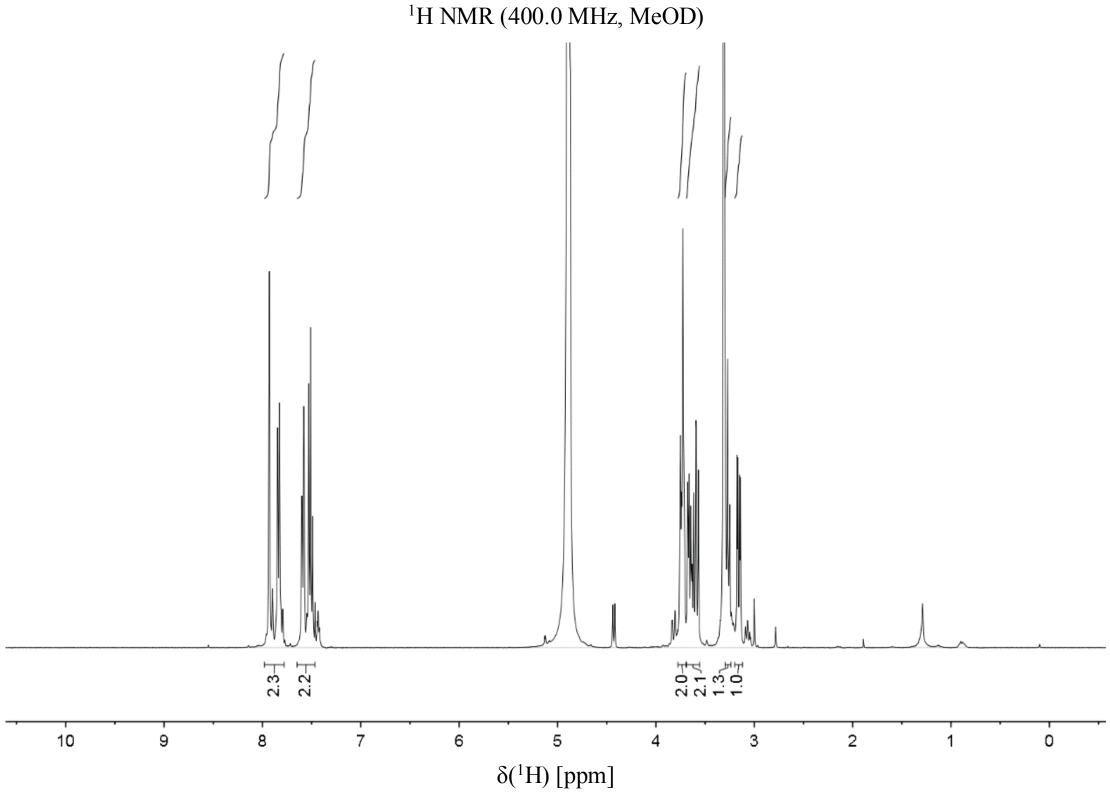


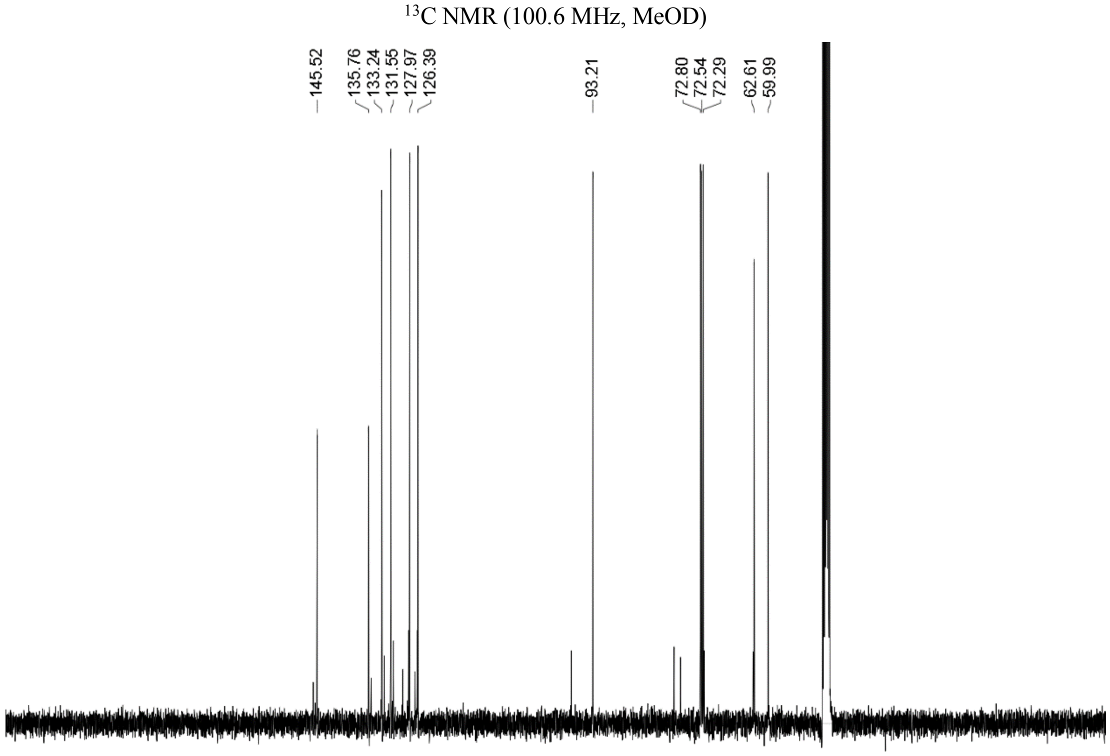

**(N-(Benzyloxycarbonyl)-2-aminoethyl)-2-deoxy-2-N-(2,2,2-trichlorethyloxy-carbonyl)-3,4,6-tri-O-acetyl-β-D-glucospyrannoside (11)**

$^{1}$H NMR (300.0 MHz, $CDCl_3$)

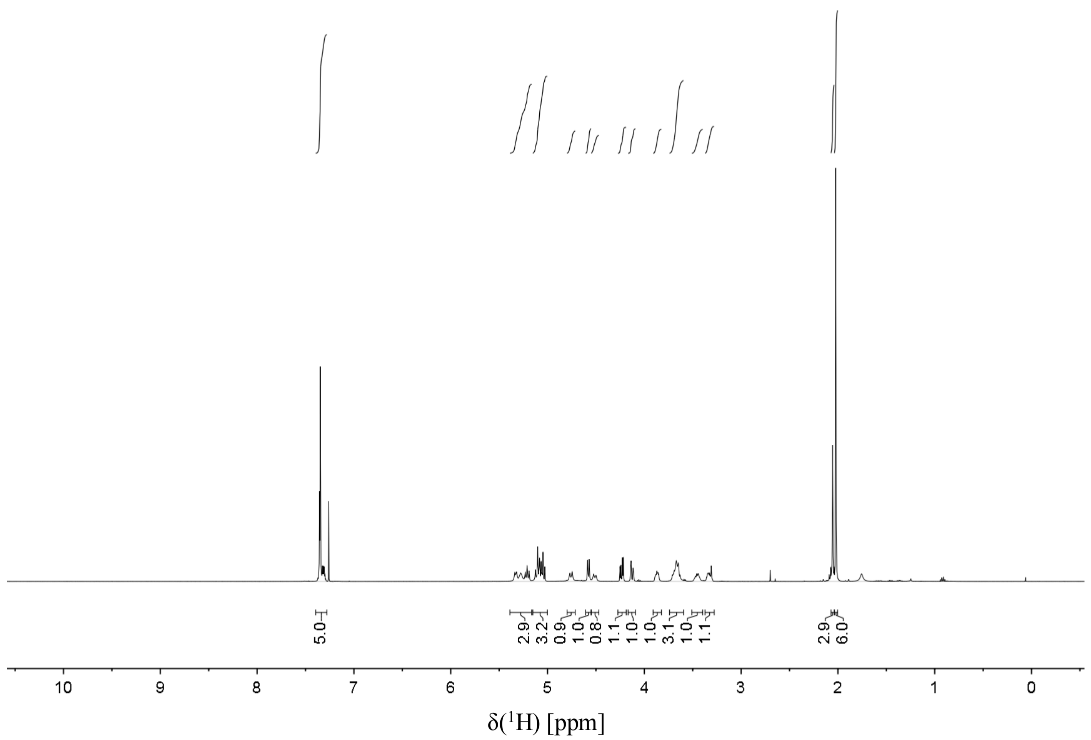


$^{13}$C NMR (125.7 MHz, $CDCl_3$)

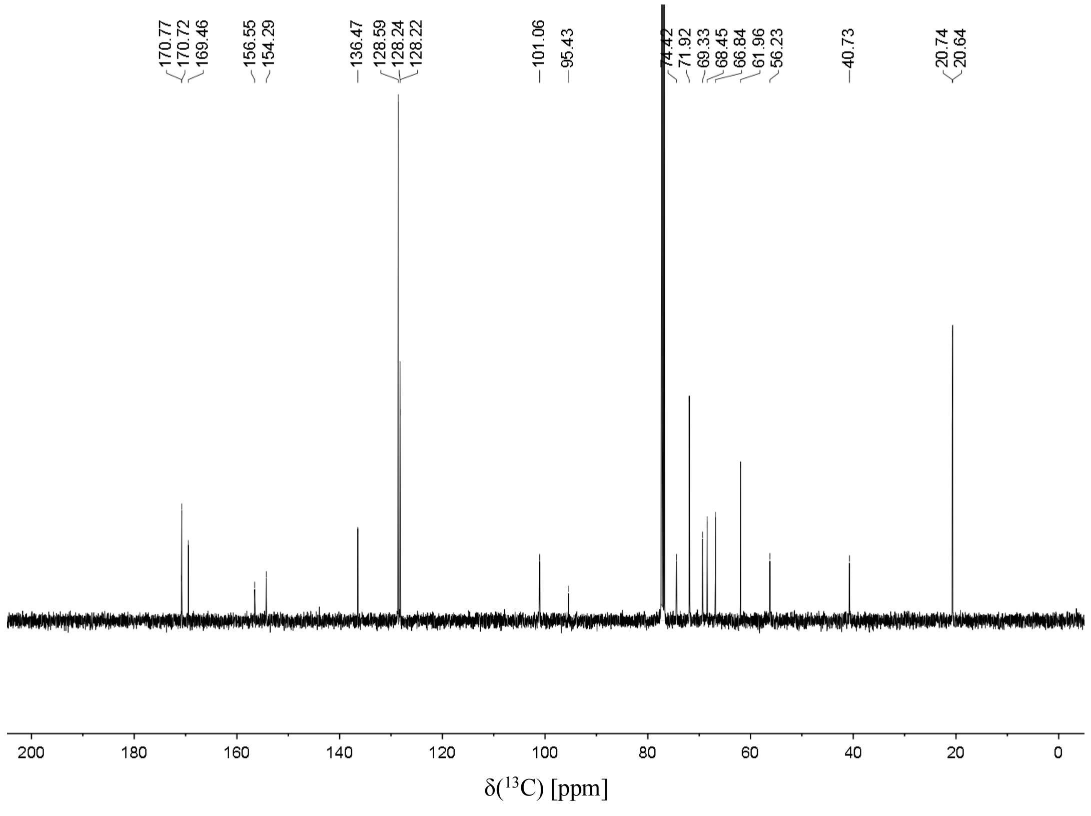

**(N-(Benzyloxycarbonyl)-2-aminoethyl)-2-deoxy-2-N-tosyl-3,4,6-tri-O-acetyl-β-D-glucospyrannoside (13)**

$^{1}H$ NMR (500.0 MHz, $CDCl_3$)

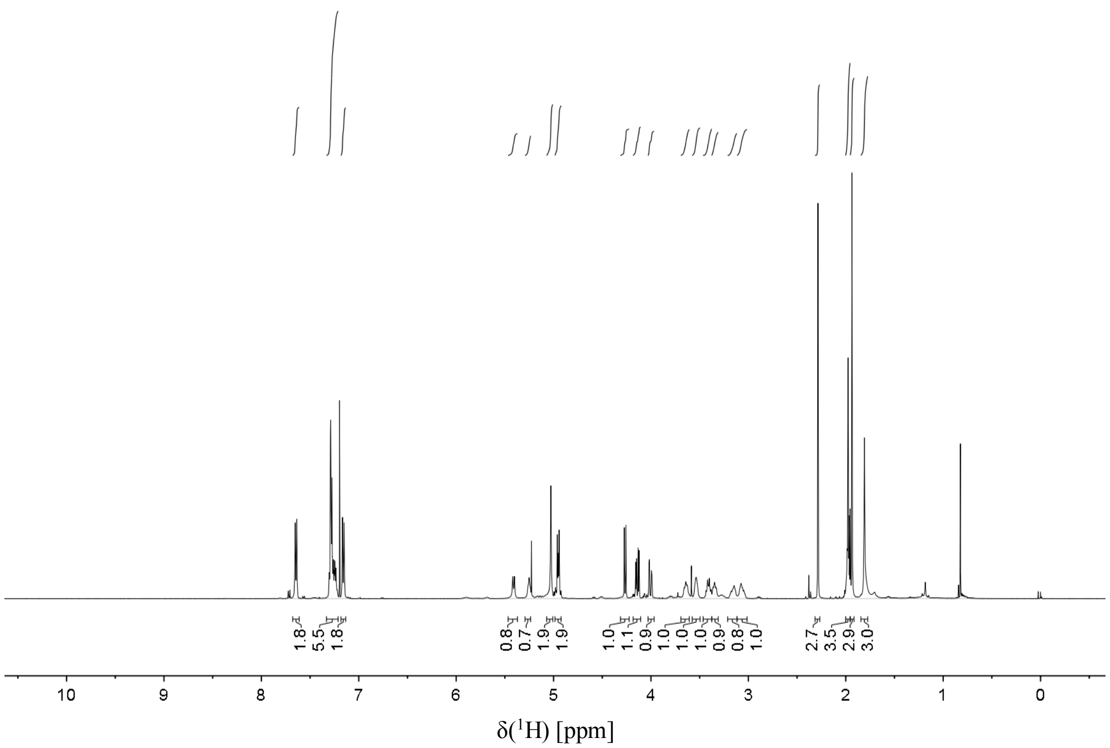


$^{13}C$ NMR (125.7 MHz, $CDCl_3$)

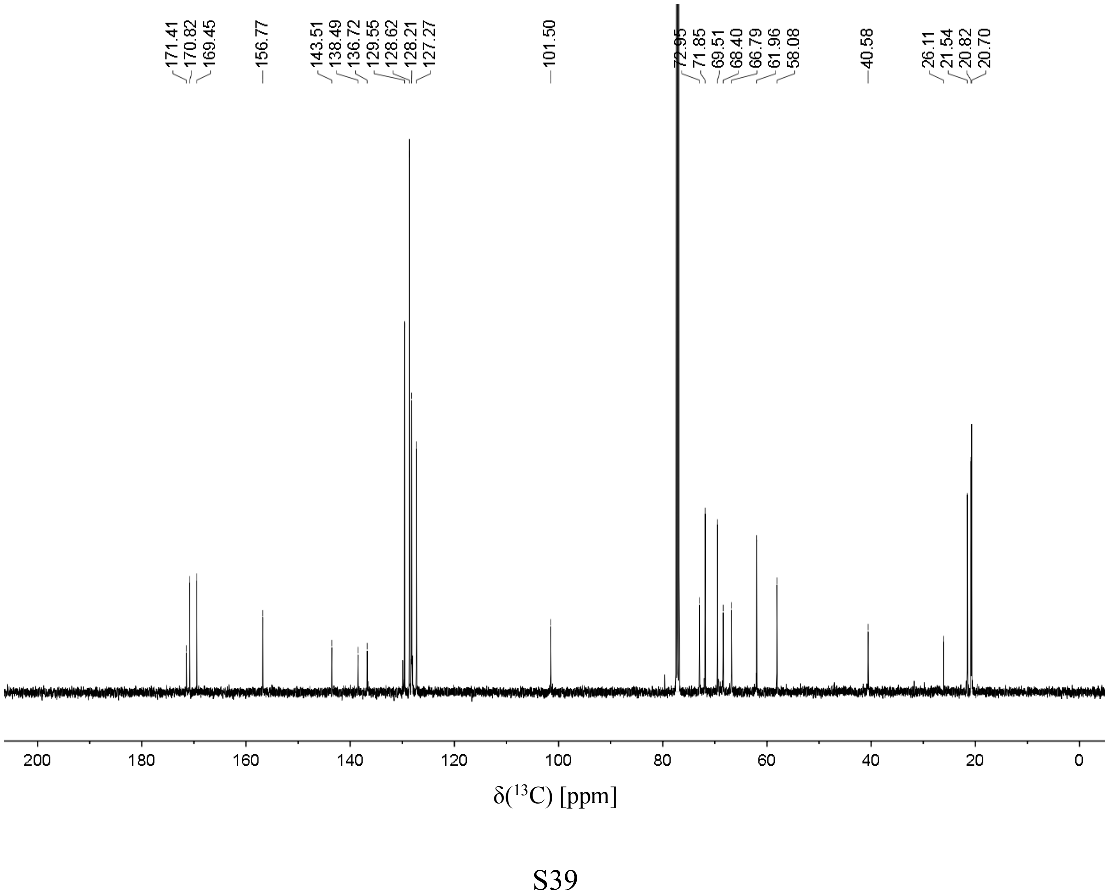

**Aminoethyl-2-deoxy-2-N-tosyl-β-D-glucospyrannoside (15)**

$^{1}H$ NMR (500.0 MHz, $D_2O$)

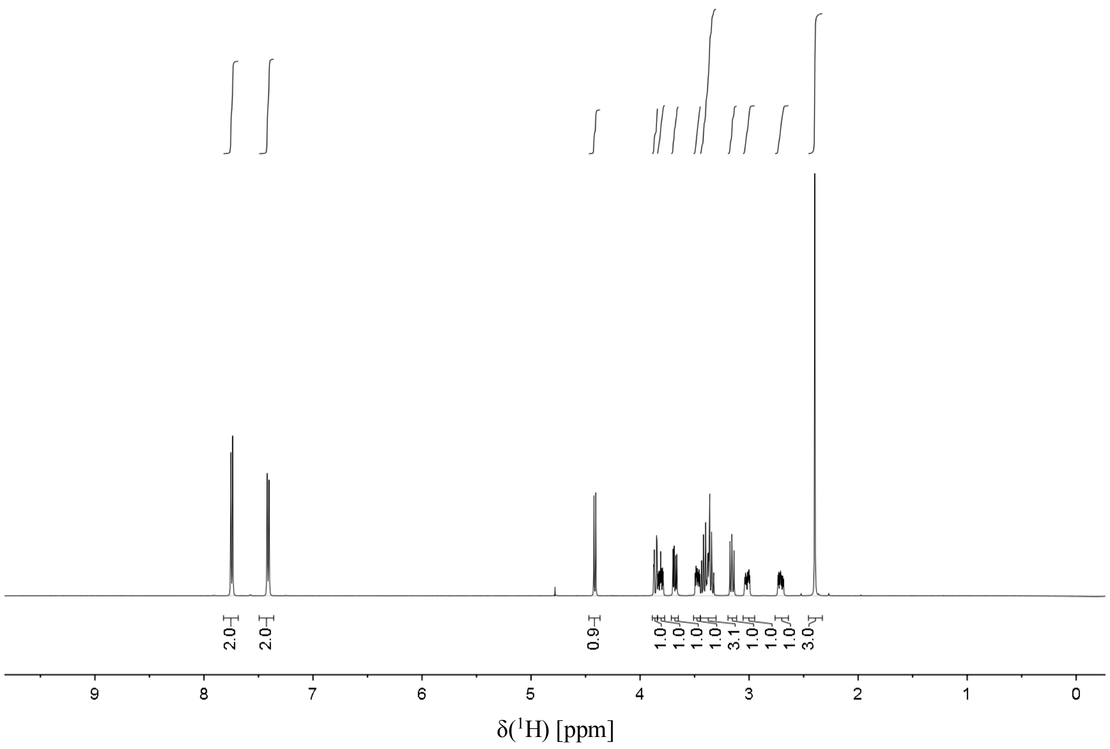


$^{13}C$ NMR (125.7 MHz, $D_2O$)

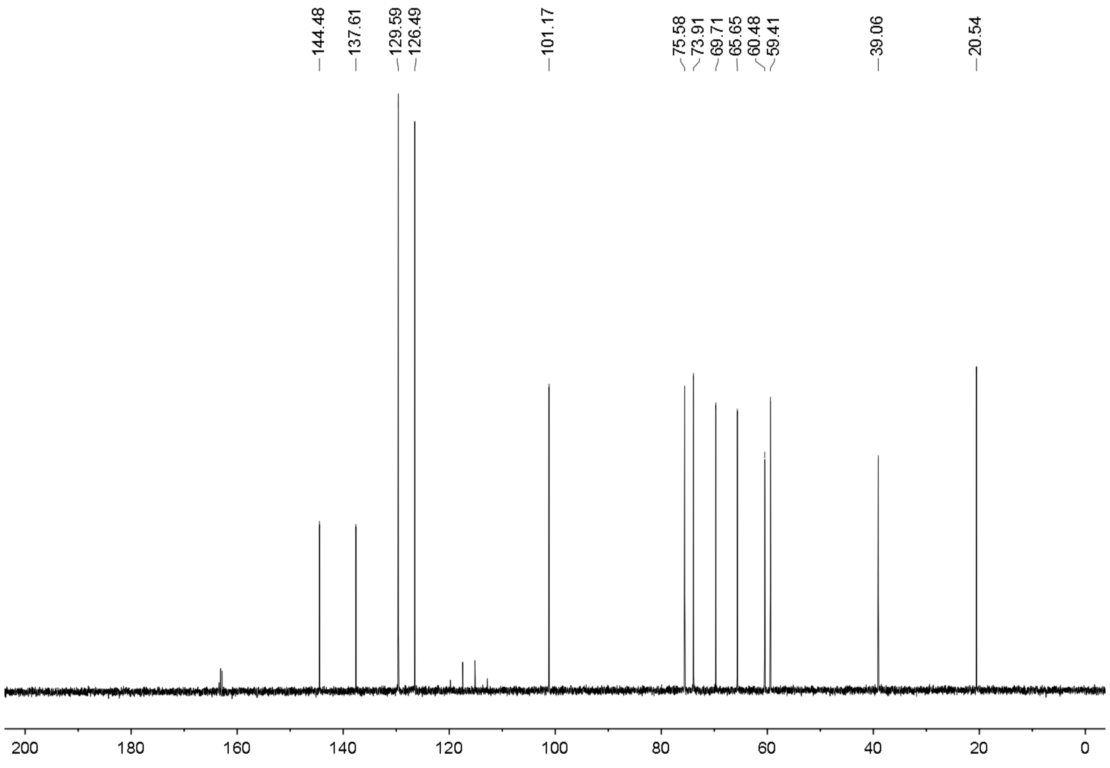

**2-Acetamidoethyl-2-deoxy-2-N-tosyl-β-D-glucospyrannoside (16)**

$^1$H NMR (400.1 MHz, $D_2O$)

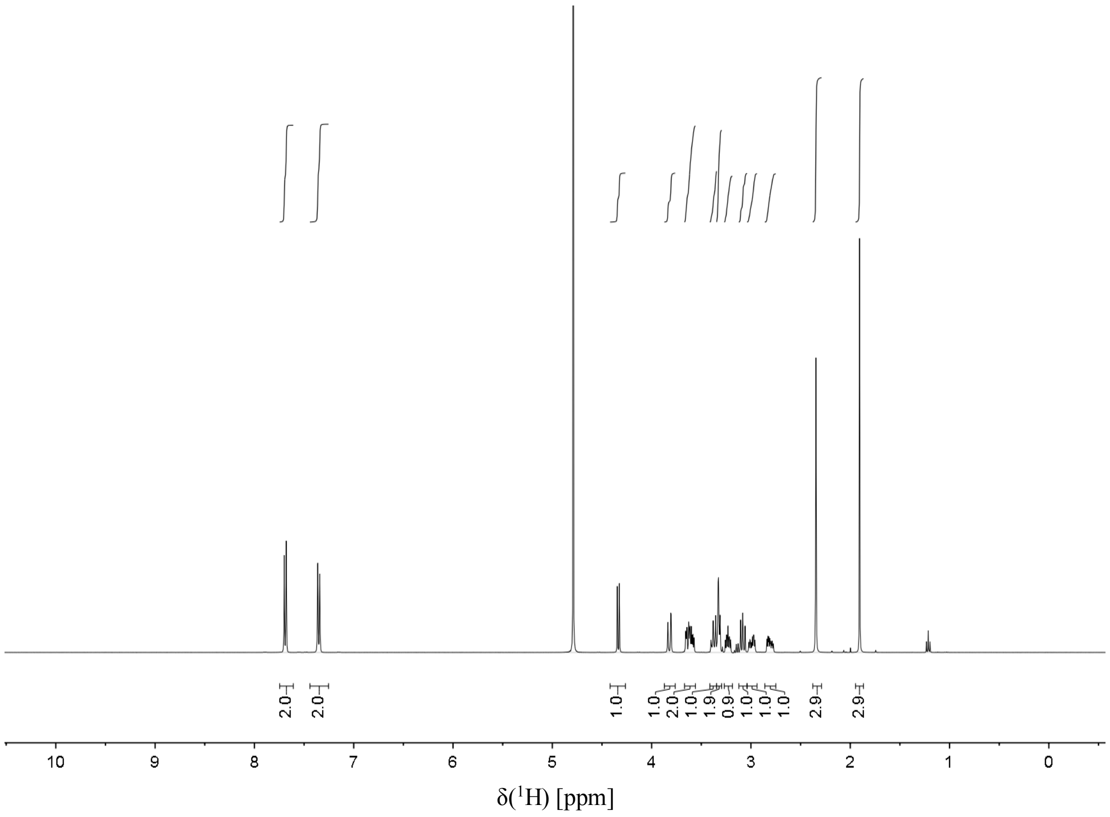


$^{13}$C NMR (100.6 MHz, $D_2O$)

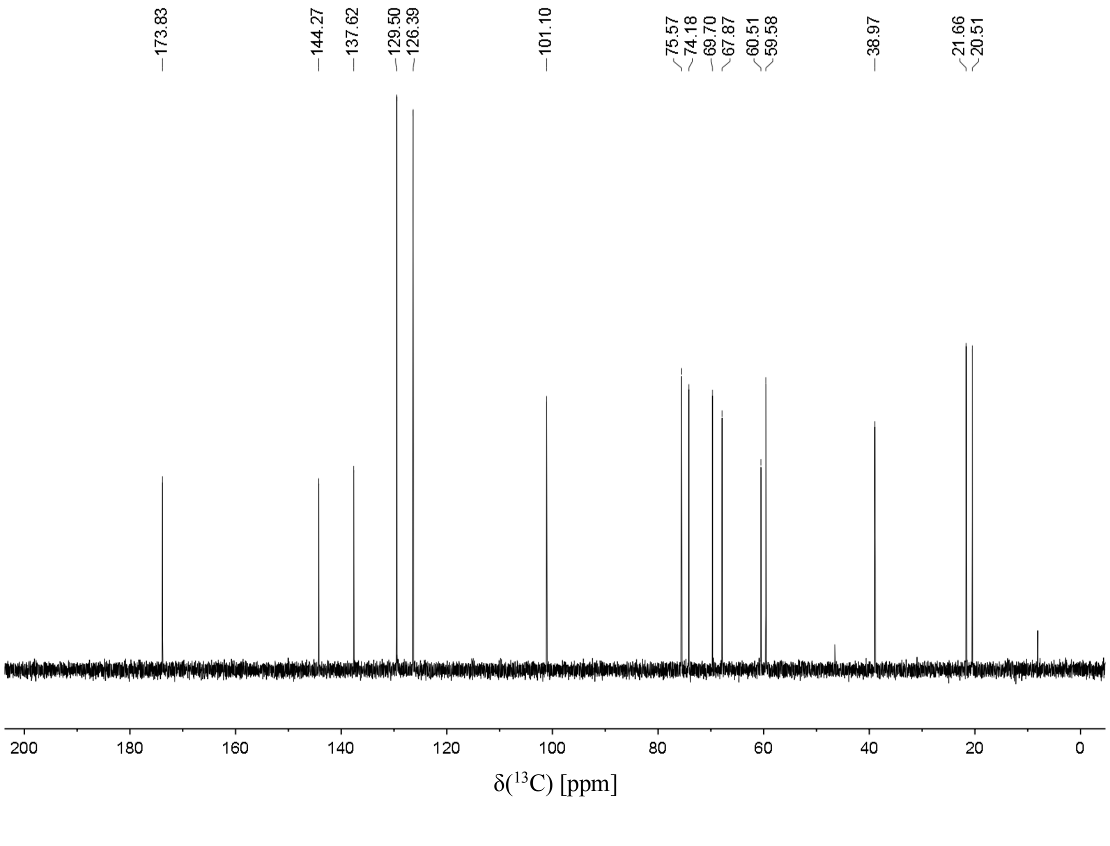

**2-Acetamidoethyl-α-D-mannopyrannoside (21)**

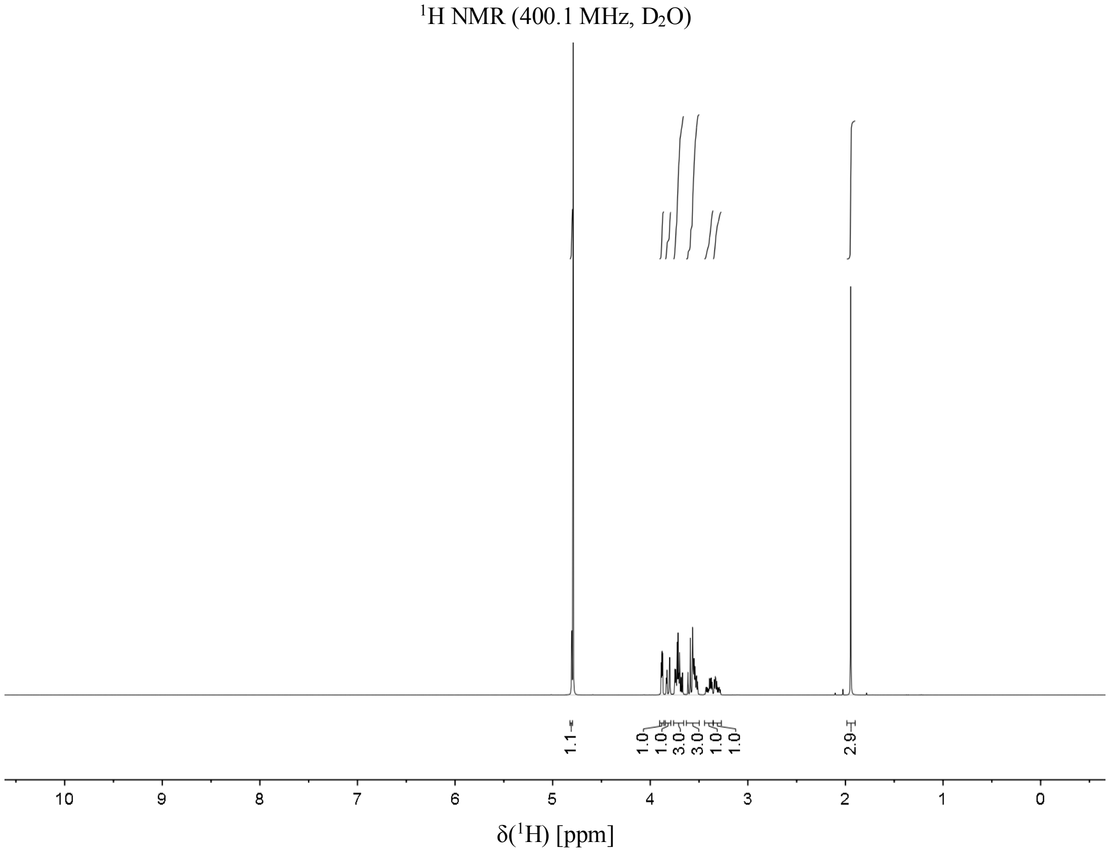


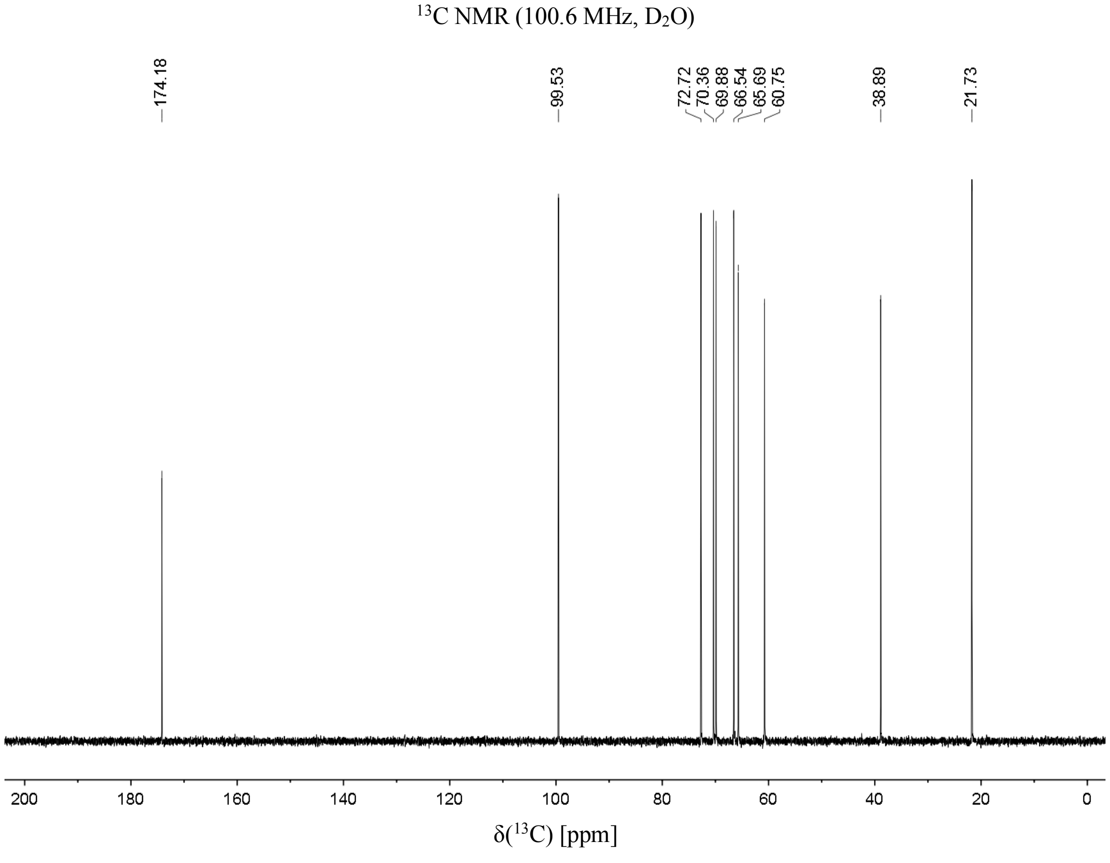

**16-PEG-DSPE (22)**

$^{1}$H NMR (400.0 MHz, DMSO-d$_6$)

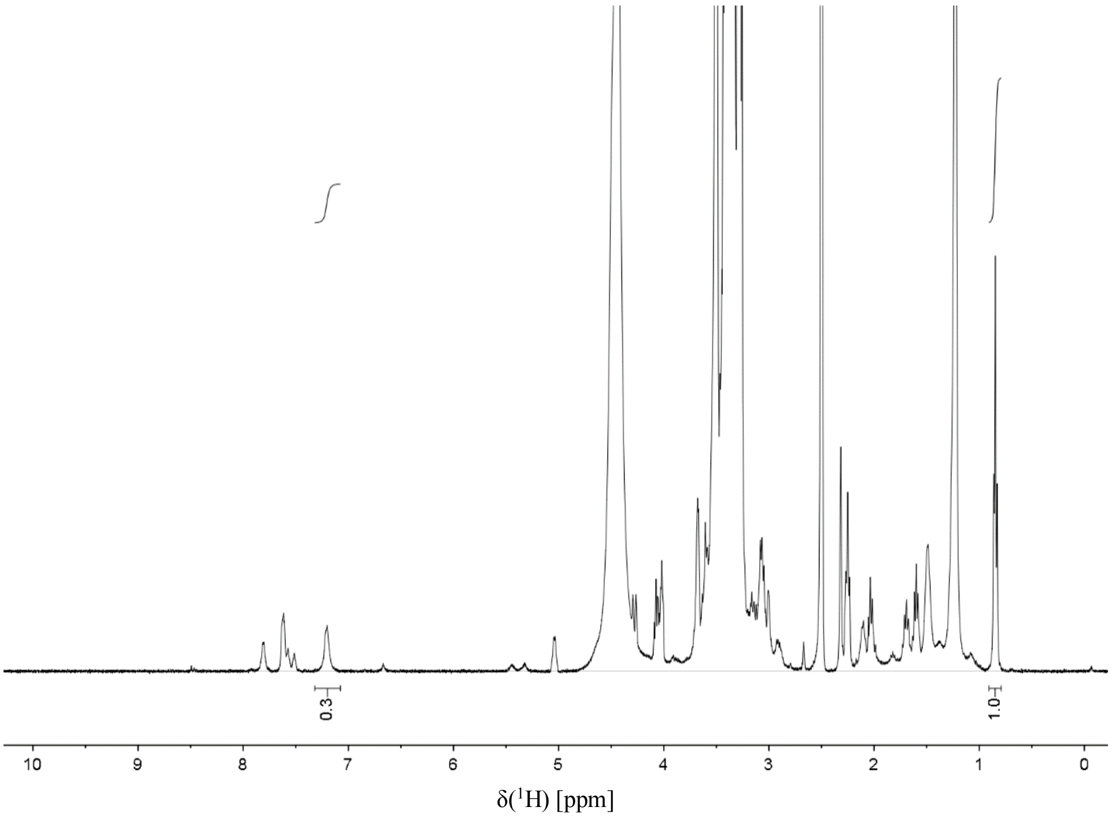


**21-PEG-DSPE (23)**

$^{1}$H NMR (400.0 MHz, DMSO-d$_6$)

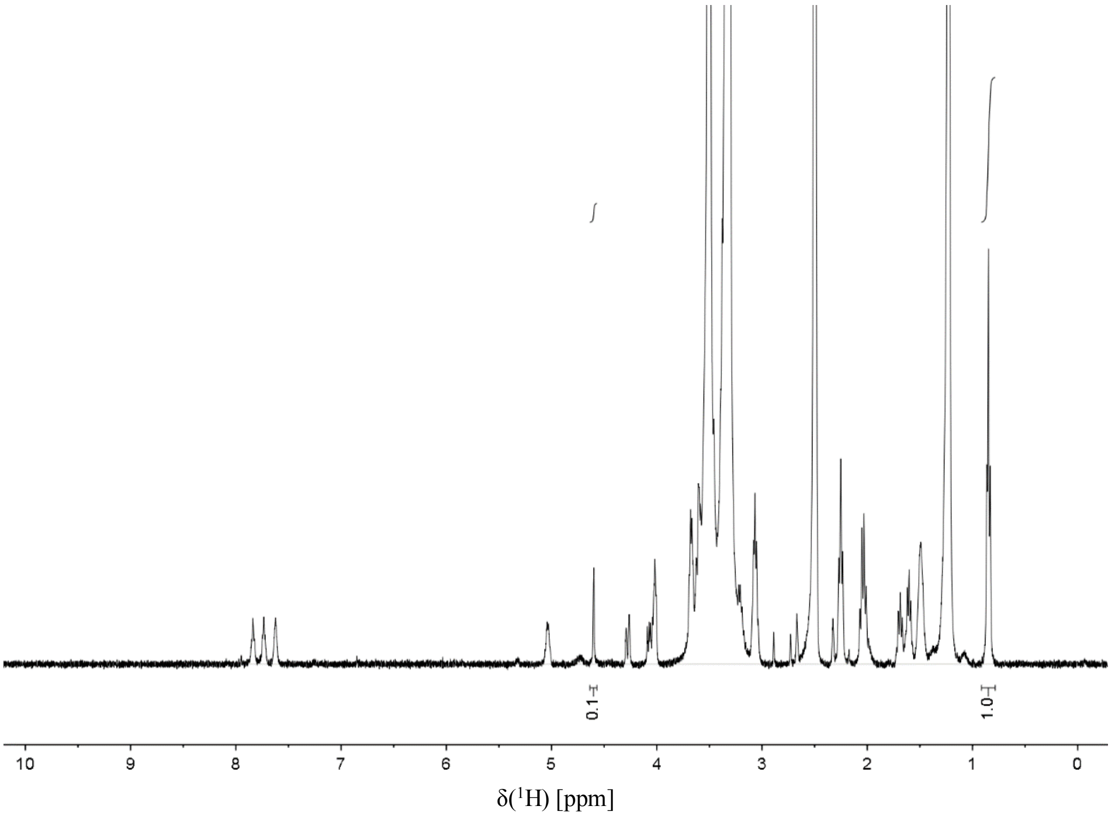
